# Phylogenetic and taxonomic insights into *Betula*: low-coverage whole genome sequencing and plastome analysis with focus on the rare Ukrainian endemic species *Betula klokovii* Zaverucha

**DOI:** 10.1101/2025.01.15.632994

**Authors:** Andrii Tarieiev, Kevin Karbstein, Oliver Gailing

## Abstract

*Betula klokovii* Zaverucha is a rare endemic species of Ukraine that is still not well taxonomically studied. In the current pilot study, we performed low-coverage whole genome sequencing for *B. klokovii*, related species (*B. pendula* Roth and *B. pubescens* Ehrh.) and the assumed hybrid *B. klokovii × B. pendula,* assessed the genomic structure of taxa with different mapping settings using UMAP non-linear dimension reduction algorithm, extracted and assembled whole plastomes.

Single Nucleotide Polymorphism (SNP) analysis based on low-coverage whole genome sequencing (LC-WGS) followed by UMAP visualization reveals the separation of *B. klokovii* from the other analysed taxa. The best taxonomic resolution was achieved with reads filtered for contamination. In contrast, the best result in obtaining complete plastome assemblies was achieved via NOVOPlasty pipeline and raw reads.

The size for the eight newly assembled plastid genomes ranges between 160,535 and 160,625bp, GC content is 36,1%. We annotated 130 genes (113 unique) for all eight assemblies. In addition, we investigated *B. klokovii’s* relationships with 20 other birch species and two intraspecific taxa by reconstructing plastome-based Bayesian inference and maximum likelihood phylogenies. Overall, plastome phylogeny provides a better resolution in comparison to phylogenies based on a few plastid or nuclear molecular markers. However, it could be affected by chloroplast capture, and some other factors like quality of sequencing and assembly, and is not suitable to detect hybrids when used alone.

In particular, we found that *B. klokovii* is likely a separate taxon that is closely related to *B. pubescens* but morphologically and genetically distinct. The study shows that genome-wide SNP data and plastome phylogenies have a certain potential for addressing issues with specific taxa within the genus *Betula* L. However, to fully leverage the potential of this approach, we suggest collecting a much larger number of birch plastome sequences to be sequenced and assembled. For a better understanding of birch phylogeny there is a need for reference-grade chromosome scale genome assemblies for polyploid species.

## Introduction

A better understanding of recent plant speciation is especially crucial for endangered endemic taxa and their preservation. Plant speciation occurs in different ways. Often it involves whole genome duplication (WGD) (Clark & Donoghue, 2018) usually followed by an extended rediploidization process (Tate & al., 2005; Marchant & al., 2016). In addition, hybridization also plays an important role in plant speciation (Abbott & al., 2013; Soltis, 2013) resulting in reticulate evolution of plant species (Linder & Rieseberg, 2004; Vriesendorp & Bakker, 2005; McBreen & Lockhart, 2006; Karbstein & al., 2022, 2024). Natural hybridization and polyploidy play an important role in the evolution of several plant genera including *Betula* and can impact adaptation processes, e.g., by the transfer of adaptive alleles between related species with different ecological preferences (Rieseberg & Willis, 2007; Strasburg & al., 2012; Abbott & al., 2013; Leroy & al., 2020).

The genus *Betula* L. is mostly represented by common pioneer trees and shrubs of temperate and boreal zones in the Northern Hemisphere. They are characterized by high vegetative variability, phenotypic plasticity, a set of different ploidies (from 2n=2x=28 to 2n=12x=168) (Ashburner & McAllister, 2016), and frequent hybridization and introgression between different species (Johnsson, 1945, 1949; Anamthawat-Jónsslon & Tómasson, 1999; Thórsson & al., 2001; Anamthawat-Jonsson, 2003; Palmé, 2003; Thórsson & al., 2010; Thomson & al., 2015b). Hence, the phylogeny of this group is very complex, and the estimated number of birch species in the world flora varies from 30 (Jong, 1993) to over 120 with an intermediate estimate of ca. 65 (Ashburner & McAllister, 2016). The respective estimates for Europe vary from 7 (de Jong, 1993) to ca. 30 according to the most recent data, including 5–7 species with hybrid origin and nearly 10 infraspecific taxa (Ashburner & McAllister, 2016; POWO, 2025). The number of existing taxonomic names (including all combinations and synonyms) is much greater – 529 specific and 514 infraspecific ones according to IPNI (IPNI, 2025). The genus has an internal structure with several subgenera and sections (Regel, 1865; Jong, 1993; Skvortsov, 2002; Tzvelev, 2002; Ashburner & McAllister, 2016). According to the most recent classification (Ashburner & McAllister, 2016), there are three subgenera and seven sections: subgenus *Betula* with sections *Costatae*, *Dauuricae*, *Apterocaryon,* and *Betula*; subgenus *Aspera* with sections *Asperae* and *Lentae*; and subgenus *Acuminata* with a single section *Acuminatae*, respectively.

There are several birch taxa in the European flora that are/were considered to be endemic or rare: *Betula aetnensis* Raf. (Rafinesque, 1814; Presl, & Presl, 1822). Deliciae pragenses, historiam naturalem spectantes (p. 141). Sumtibus Calve.), *B. borysthenica* Klokov (Klokov, 1946), *B. celtiberica* Rothm. & Vasc. (Rothmaler & Vasconcellos, 1940), *B. klokovii* Zaverucha (Zaverucha, 1964), *B. oycoviensis* Besser (Besser, 1809), a series of dark-barked birches (*B. atrata* Domin (Domin, 1927), *B. obscura* Kotula ex Fiek (Fiek, 1888), *B. kotulae* Zaverucha (Zaverukha & al., 1986)) and some others. They were described at different times and were initially identified based on morphological characters. Some of these taxa were included in more recent studies (e.g., *B. oycoviensis,* (Baláš & al., 2016; Vítámvás & al., 2020), and some dark-barked birches (Tarieiev & al., 2019)), some of them were reassessed (Ashburner & McAllister, 2016; Govaerts, 2024; POWO, 2025), but most rare endemic birch taxa remain understudied.

*Betula klokovii* is a Ukrainian strict endemic birch species (Fig. 1A–G). It was described in 1959 by the Ukrainian botanist Borys Zaverukha from two small mountains, Strakhova and Maslyatyn, that are close to Kremenets in the Ternopil region, Ukraine (Zaverucha, 1964). The species is growing in specific conditions on chalk-calcareous soil near the top of these mountains or hills (Fig. 1F, G).

**Fig. 1.**
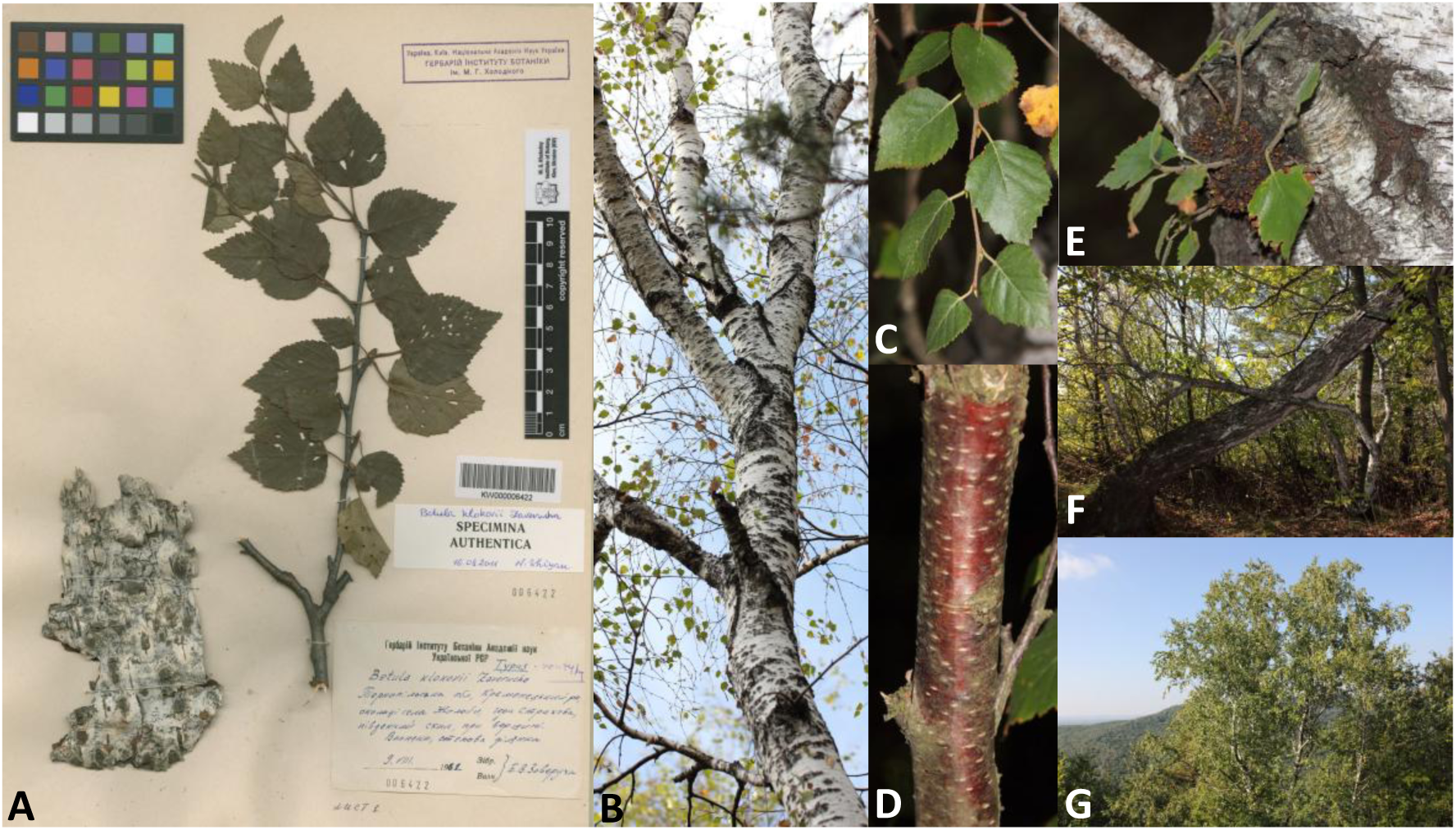
Original specimen of *Betula klokovii* Zaverucha from the National Herbarium of Ukraine (KW000006422) (**A**), stem of the tree with characteristic dotted pattern (**B**), young twig with leaves (**C**), bark on young twigs (**D**), area with dormant buds (**E**), and trees in natural habitat (**F, G**)

Klokov’s birch is a 5–10m tall tree, with 1–2 stems and white bark that does not peel off and has a specific dotted pattern on old trunks (dark grey dots, covering up to 50% of the surface). Young twigs are non-pendulous, with cherry-brown bark color, densely or sparsely pubescent. Buds are sticky, pointed, 4–6 mm long. Leaves are rhombic-ovate, or oblong-ovate with a cuneate base, 25–55 mm long, and 15–40 mm wide. The adaxial surface is dark green, glabrous, whereas the abaxial surface is pale greyish green, somewhat pubescent (Zaverucha, 1964).

*B. klokovii* is presumably tetraploid and closely related to tetraploid *B. pubescens* Ehrh. (Zaverucha, 1964; Tutin, 1993; Tzvelev, 2002). The type specimens (holotype KW000006416 (https://plants.jstor.org/stable/10.5555/al.ap.specimen.kw000006416); isotypes KW000006417 – KW000006421; original material (specimina authentica) KW000006422 – KW000006425) are stored in the National Herbarium of Ukraine (KW), M.G. Kholodny Institute of Botany, NAS of Ukraine (Fig. 1A, Olshanskyi & al, 2016).

Both the very low number of trees and a very small natural habitat pose a significant threat to this species. Consequently, it is protected on local, and state levels in Ukraine (Kagalo & al., 2009), and also included in the IUCN red list as a critically endangered species (Rivers & Tarieiev, 2015).

To improve the protection of this species for nature conservation, the trees from two known populations were morphologically identified and localized using GPS coordinates in 2014–2019 by Mr. Tarieiev and the staff of Natural Nature Park (NNP) “Kremenets Mountains”. In addition, the staff of NNP “Kremenets Mountains” established a nursery for planting young trees from seeds (Mr. Tarieiev contributed to planning of the nursery and taxonomic identification/expertise). The distribution of *B. klokovii* and related species, as well as the nursery and territory of NNP “Kremenets Mountains” are shown in Fig. 2.

**Fig. 2.**
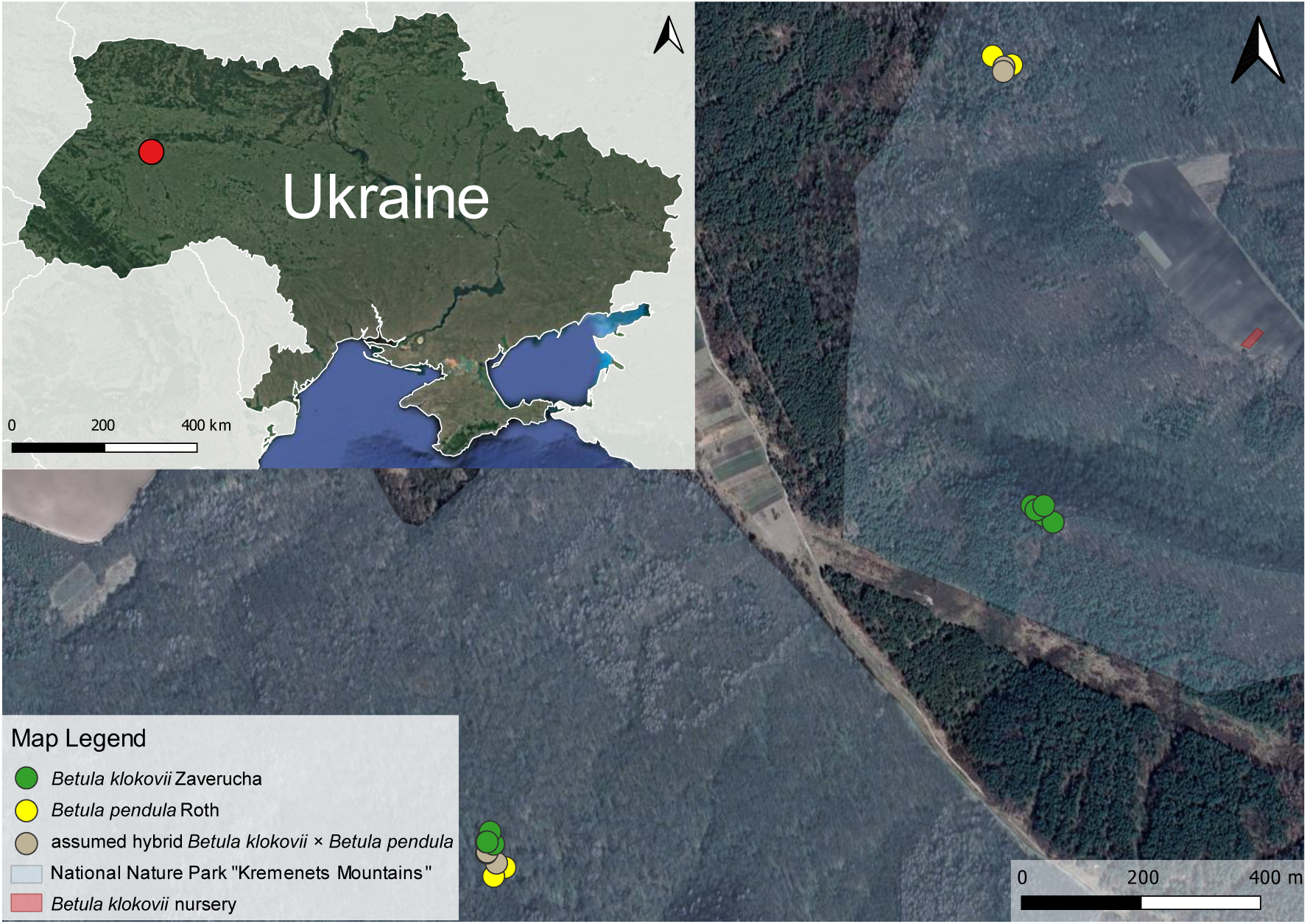
Distribution map of *Betula klokovii* Zaverucha in Ukraine (map created in QGIS 3.40 ‘Bratislava’ based on GoogleSatellite map)

Despite all morphological and distributional differences, traditional genetic markers (e.g., internal transcribed spacers, ITS (Tarieiev & al., 2021) do not resolve closely related tetraploid birch taxa including *B. klokovii*. Hence, we applied a genomic approach and performed low-coverage whole genome sequencing followed by genome-wide SNP analysis and plastome read extraction, assembly, and annotation. Whole-genome sequencing allows obtaining more data in comparison to the usage of the combination of several traditional genetic markers (McKain & al., 2018; Karbstein & al., 2024). However, there are only three birch genome assemblies available – *B. nana* L. (NCBI Accession number GCA_000327005.1)*, B. pendula* Roth (NCBI Accession number GCA_900184695.1), and *B. platyphylla* Sukaczev (Phytozome genome ID: 679). All of them represent only diploid species, and therefore the entire diversity of polyploid species is not covered. At the same time, both Klokov’s birch and downy birch are tetraploids.

Complete plastid genomes are easier to work with in terms of sequencing costs and bioinformatic analysis, and could provide valuable information about phylogenetic relationships and speciation processes inside the birch genus. There are several studies on assembling complete plastid genomes for different birch species (Hu & al., 2017; Wang & al., 2018; Lee & al., 2019; Wang & al., 2019; Yin & al., 2019; Hua & Zhu, 2020; Yin & al., 2020; Yoshida & al., 2020; Zhang & al., 2023), and some studies on Betulaceae (Yang & al., 2019b; Hu & al., 2020; Yang & al., 2021), which enable comparative plastome-based phylogenetic studies. In birch, plastids are maternally inherited and, applied alone, are not well-suited for detecting hybridization. Chloroplast DNA is non-recombining which allows treating the entire plastome as a single locus, reducing population size, and enabling recovery of ancestral lineages but also the (maternal) progenitor lineage of hybrids (Palmé, 2003; Palmé & al., 2003). Also, haploid plastid genomes make it easier to perform a phylogenetic analysis involving samples with different ploidies (Gitzendanner & al., 2018), which is highly relevant for the genus *Betula* L. The limitations of plastome-based studies include not only uniparental inheritance but also the fact that plastomes are more prone to stochastic events and there are known cases of chloroplast capture (Thomson & al., 2015a; Yang & al., 2021; Touchette & al., 2024) when plastids of one species are replaced by plastids from another one (usually as a part of cytoplasmic capture) as a result of interspecific hybridization/introgression that could have an impact on the phylogeny.

*<u>Objectives of the study</u>*: performing low-coverage short read whole genome sequencing on herbarium specimens of *B. klokovii, B. pubescens, B. pendula* and assumed hybrids *B. klokovii × B. pendula* followed by genome-wide SNP analysis, plastome assembly, and plastome phylogeny reconstruction with the aim to clarify phylogenetic position of *B. klokovii*.

## Material and methods

Based on the analysis of morphological traits, we suspected cases of hybridization between *B. klokovii* and *B. pendula* Roth, especially for the population located on Mt. Maslyatyn.

Birch samples (twigs with leaves) from *B. klokovii, B. pendula,* and their presumable hybrid were collected by Mr. Andrii Tarieiev and the staff of the NNP “Kremenets Mountains” in 2019. Samples from *B. pubescens* were collected by Dr. Serhii Panchenko in Sumy region, Ukraine (details about collection sites are given in Supporting information,Table S1).

### DNA isolation

DNA extraction was performed from dried leaf material using Qiagen DNeasy Plant Mini Kit (Hilden, Germany) according to the manufacturer’s instructions. The quantity and integrity of the DNA was checked by NanoDrop (ThermoScientific) (Supporting information, Table S2) and electrophoresis in a low-concentration (0.8 %) agarose gel.

### Low-coverage whole genome sequencing (LC-WGS)

Low-coverage whole genome sequencing (LC-WGS) was performed for eight different samples (two *B. klokovii*, two – *B. pendula*, two – prospective hybrids between them (*B. klokovii* × *B. pendula*) from the same location, and two samples of *B. pubescens* from another location in Ukraine). Library preparation, multiplexing, and sequencing (MiSeq 2×300PE) were conducted at the Competence Centre for Genomic Analysis (CCGA) Kiel within the DFG competence network as a pilot study.

### Bioinformatic analysis for LC-WGS

#### Quality assessment, trimming, filtering, and mapping to the references

Obtained reads were analyzed using several different approaches to check the quality, contamination, and to extract, and assemble plastid genomes. The analysis was mostly done on the high-performance scientific computing cluster of the GWDG (Göttingen, Germany). Scripts and details about the analysis are deposited on GitHub (https://github.com/AndriiTarieiev/Betula_klokovii_LC-WGS_data_analysis). Assemblies in CLC genomics workbench were performed locally.

First, we performed quality assessment of obtained reads in FastQC v0.11.4 (https://www.bioinformatics.babraham.ac.uk/projects/fastqc/), and trimming of reads and filtering adaptors in Trimmomatic v0.36 (Bolger & al., 2014). Screening for contamination and filtering was done in Kraken2 v2.1.2 (Wood & al., 2019; https://github.com/DerrickWood/kraken2) against the following newly created databases: archaea, bacteria, fungi, human, protozoa, viral, and univec. Mapping of the cleaned reads on three available genome assemblies (for *B. nana, B. pendula,* and *B. platyphylla*, respectively) was performed using bowtie2 (Langmead & Salzberg, 2012). For this purpose, custom indices were specially created for available genome assemblies of *B. nana* L. (NCBI Accession number GCA_000327005.1)*, B. pendula* (NCBI Accession number GCA_900184695.1), and *B. platyphylla* Sukaczev (Phytozome genome ID: 679; Chen & al., 2021), respectively. In addition, we tested several alignment options for the reads against references:

i. default end-to-end alignment using unfiltered reads;
ii. local alignment using unfiltered reads;
iii. local alignment based on filtered paired and unpaired reads;
iv. local alignment based on filtered paired and unpaired reads without unaligned reads (--local --nounal).

#### Genome-wide SNP calling

Picard v2.20.2 (https://broadinstitute.github.io/picard/) and samtools v1.6 (Danecek & al., 2021) were used to convert .sam to .bam files, and to index .bam files. Sambamba v1.1.0 was applied to mark and remove duplicates. To call variants, we ran GATK v4.4.0 HaplotypeCaller (McKenna & al., 2010; Poplin & al., 2017; Auwera & O’Connor, 2020) based on different ploidy level settings according the estimated ploidy of the given species. The individual g.vcf files for each genome reference and filtering settings were merged in a .vcf alignment file using gatk CombineGVCFs. Variants were genotyped using gatk GenotypeGVCFs, and filtered using gatk VariantFiltration and gatk SelectVariants (except for --max-nocall-fraction 0.13 to ensure meaningful phylogenetic comparisons with low levels of missing data) with default settings.

We were able to use only mapping to the *B. platyphylla* genome assembly since genome assemblies for *B. nana* and *B. pendula* are not chromosome-scale and contain too high numbers of contigs for GATK4.

No further filtering regarding allele depth and frequency was conducted to not bias the study of different mapping references and filtering settings. Obtained vcf files were converted into sample-wise genetic distance matrices across all SNP sites in R using the R packages vcfR v1.15.0 (Knaus & Grünwald, 2016, 2017), adegenet v2.1.10 (Jombart & Kamvar, 2007; Jombart & Ahmed, 2011), and ape v5.8 (Paradis & al., 2002, 2004).

#### Uniform Manifold Approximation and Projection (UMAP)

Plots based on different mapping settings (see above) were visualized using UMAP (k=3, epochs= 1000, dimensions=2) (McInnes & al., 2018) and the R packages uwot v0.2.2 (Melville, 2019) and ggplot 2 v3.5.1 (Wickham & al., 2007). UMAP, as a dimension reduction approach, is able to preserve both local and global non/linear structures, and is therefore highly suitable to visualize species-specific relationships (Hodač & al., 2023, 2024). Since UMAP is a stochastic algorithm that uses randomness to speed up approximation steps and to solve optimization problems, it could produce somewhat different results (https://umap-learn.readthedocs.io/en/latest/reproducibility.html). To overcome this, we introduced 1000 iterations of bootstrap resampling (Supporting information, Script S2; https://github.com/AndriiTarieiev/Betula_klokovii_LC-WGS_data_analysis).

### Plastome extraction and assembly

We attempted to obtain plastid genome assemblies in two different ways:

1. QIAGEN CLC Genomics Workbench v23.0.2 and 23.0.3 (https://digitalinsights.qiagen.com) with filtered and unfiltered reads (details in Supporting information, Table S3).
2. NOVOPlasty v4.3.1. (Dierckxsens & al., 2017) with different k-mer sizes (23, 33, and 45), and also with and without reference sequences to identify the optimal parameters of assembly (details in Supporting information, Table S4).

The reason behind using unfiltered reads for plastome assemblies is that some part of plastid and/or mitochondrial reads could be mistakenly filtered out as bacterial contamination (which was also true in our case, details in Table 1). That is why it is even recommended by NOVOPlasty developers to use unfiltered raw whole genome set of reads as input (https://github.com/ndierckx/NOVOPlasty).

**Table 1.**
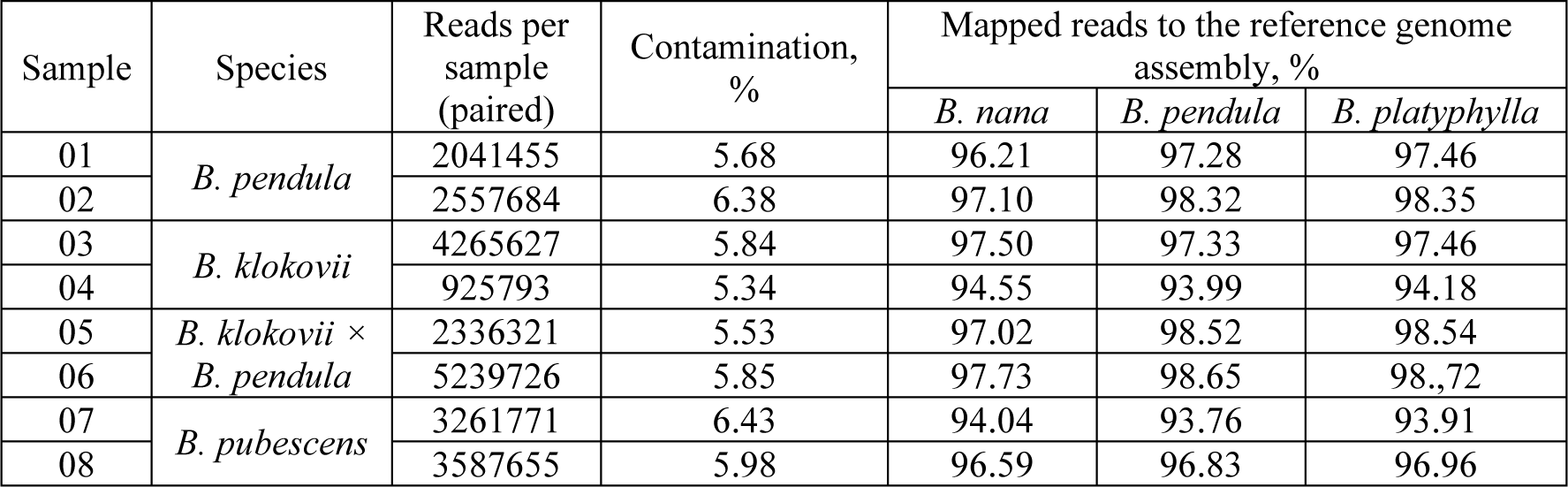
Basic statistics on LC-WGS data

Circularized plastome assemblies were annotated using GeSeq (Tillich & al., 2017) and the following references: NC_072281.1 and NC_044852.1 for annotation of *B. pendula* samples and the prospective hybrid *B. klokovii × B. pendula*, NC_039996.1 for annotation of *B. klokovii* and *B. pubescens*, respectively. To infer additional details (forward and reverse repeats, tandem repeats, and SSRs) visualization of annotated assemblies was performed in OrganellarGenomeDRAW (OGDRAW) v1.3.1 (Greiner & al., 2019).

GeSeq annotation was checked manually in GeneiousPrime, and compared to several reference sequences from NCBI. Annotation was converted to a feature table using GB2seqin file converter (Lehwark & Greiner, 2019) within CHLOROBOX (https://chlorobox.mpimp-golm.mpg.de/index.html) and edited manually to meet NCBI submission criteria.

### Plastome phylogeny

For plastome phylogeny reconstruction, we retrieved 66 complete plastid genome assemblies from NCBI GenBank and used them for an alignment together with our 8 assemblies for *B. klokovii, B. pendula*, prospective hybrid *B. klokovii* × *pendula*, and *B. pubescens*. As an outgroup, we used the sequence of *Alnus glutinosa* L. (Betulaceae; NC_039930.1), retrieved from NCBI GenBank. Alternatively, we removed all duplicates representing identical assemblies and retained only unique/nonredundant ones (8 own + 52 from NCBI Genbank + outgroup) for the alignment. All the analysis was performed twice – for the entire set of sequences and for the one containing non-duplicated ones only.

The specific problem with aligning circular DNA is that it can be linearized in several ways. For alignment and phylogeny reconstruction, there is a need to have all plastomes linearized in the same way with the same starting point. That is why we have developed our own approach, which includes the following steps:

In the first step, all plastome sequences were aligned using MAFFT v7.520 (Katoh, 2002; Katoh & Standley, 2013) in slow accurate mode (FFT-NS-i). Then a single unique conserved motif present in all the sequences was selected and used as a starting point. To achieve this, we developed an original small script based on a series of UNIX commands (Supporting information, Script S1, https://github.com/AndriiTarieiev/relinearization_circular_sequences/) that works with non-interleaved fasta formatted plastome multiple sequence alignment (MSA).

After getting MSA with the same starting point for all sequences (Supporting information, Alignment S1 with all the sequences and Alignment S2 with only unique ones; also deposited on GitHub: https://github.com/AndriiTarieiev/Betula_klokovii_LC-WGS_data_analysis), we proceeded with test runs for phylogeny reconstruction that were based on a single model assigned by ModelTest-NG 0.2.0 (Darriba & al., 2020) according to Bayesian information criterion (BIC), Akaike information criterion (AIC), and Akaike information criterion with small sample correction (AICc), respectively. Different criteria were used in order to test for differences between them (Susko & Roger, 2020). Identified models were used to reconstruct phylogenetic trees based on Bayesian Inference (GTR+I+G) and Maximum Likelihood (TVM+I+G).

Bayesian Inference reconstruction was performed in the parallel version of MrBayes 3.2.6 (Altekar & al., 2004; Ronquist & al., 2012) with 100,000,000 iterations, burnin 10%, and GTR+I+G model. Maximum Likelihood reconstructions were performed in RAxML-NG 1.2.0 (Kozlov & al., 2019) with 10000 bootstrap iterations and TVM+I+G model, IQ-TREE v1.6.12 and IQ-TREE v2.2.2.6 (Nguyen & al., 2015; Minh & al., 2020) with – 10,000 ultrafast bootstrap iterations with TVM+I+G models, respectively.

Conversion from fasta to nexus file format required for MrBayes and IQ-TREE was performed using seqmagick (https://github.com/fhcrc/seqmagick).

The next stage was phylogeny reconstruction based on plastome assemblies that were partitioned according to the annotation. To achieve that, we created a consensus sequence for plastome MSA and reannotated it using GeSeq (Tillich & al., 2017). Annotation was checked visually using GUI interfaces of CLC Genomics workbench 23.05 and GeneiousPrime 2023.2.1, manually corrected, and used for partitioning the alignment which was performed in Jalview2 (Waterhouse & al., 2009). Best evolutionary models were assigned to every defined part by ModelTest-NG according to BIC, AIC, and AICc in order to test for differences between different criteria (Susko & Roger, 2020). Information about the position of each part within MSA and the evolutionary model was used for partition assignment in MrBayes, RAxML-NG, and IQ-TREE, respectively. Due to limitations in the number of partitions that could be created and handled by the software, parts with the same evolutionary models were grouped together (partitioning schemes are presented in Supporting information, Partitioning S1–S9). Such analysis was performed separately for each criterion. Bayesian Inference phylogeny reconstructions in MrBayes were performed with 100,000,000 iterations and 10% burnin. Maximum Likelihood phylogeny reconstructions were performed in RAxML-NG with 10000 iterations of bootstrap, and in both versions of IQ-TREE with 10000 iterations of ultrafast bootstrap (UFB), respectively.

In addition, we reconstructed several types of phylogenetic networks in PopART (http://popart.otago.ac.nz; Leigh & Bryant, 2015) – Minimum Spanning Network, Median Joining Network (Bandelt & al., 1999) [both with epsilon=0], Integer NJ Net (French & al., 2014; Leigh & Bryant, 2015) with different tolerance to reticulation (0.5 and 0), and TCS (Clement & al., 2000, 2002).

Visualization of the trees was performed in TreeGraph2 (Stöver & Müller, 2010, http://treegraph.bioinfweb.info/), followed by coloring and minor adjustments in Inkscape 1.3 (https://inkscape.org/).

## Results

### Low-coverage genome sequencing

Basic LC-WGS read statistics per sample, total contamination rates, and mapping rates on three available genome assemblies using bowtie2 (Langmead & Salzberg, 2012) with “local” option and custom indices for *B. nana, B. pendula*, and *B. platyphylla* Sukaczev genome assemblies are shown in Table 1.

#### Genome-wide SNPs and UMAP results

GATK SNP-calling recovered 9,399 variants in the case of mapping raw reads (without filtering for contamination) of all the samples against the *B. platyphylla* genome assembly using bowtie2 default end-to-end alignment. 13,492 variants were recovered in the case of mapping raw reads (without filtering for contamination) for all samples using bowtie2 local alignment. 11,891 variants were recovered in the case of mapping reads that were filtered for contamination for all samples using local alignment considering paired and unpaired reads, and paired and unpaired reads without non-aligned ones (--local -- nounal), respectively. By repeating the analysis 10 times, we inferred that even 100 bootstrap iterations were sufficient for obtaining a reproducible result. However, increasing the bootstrap number to 1000 resulted in improved final clustering (Fig. 3, S1– S12).

**Fig. 3.**
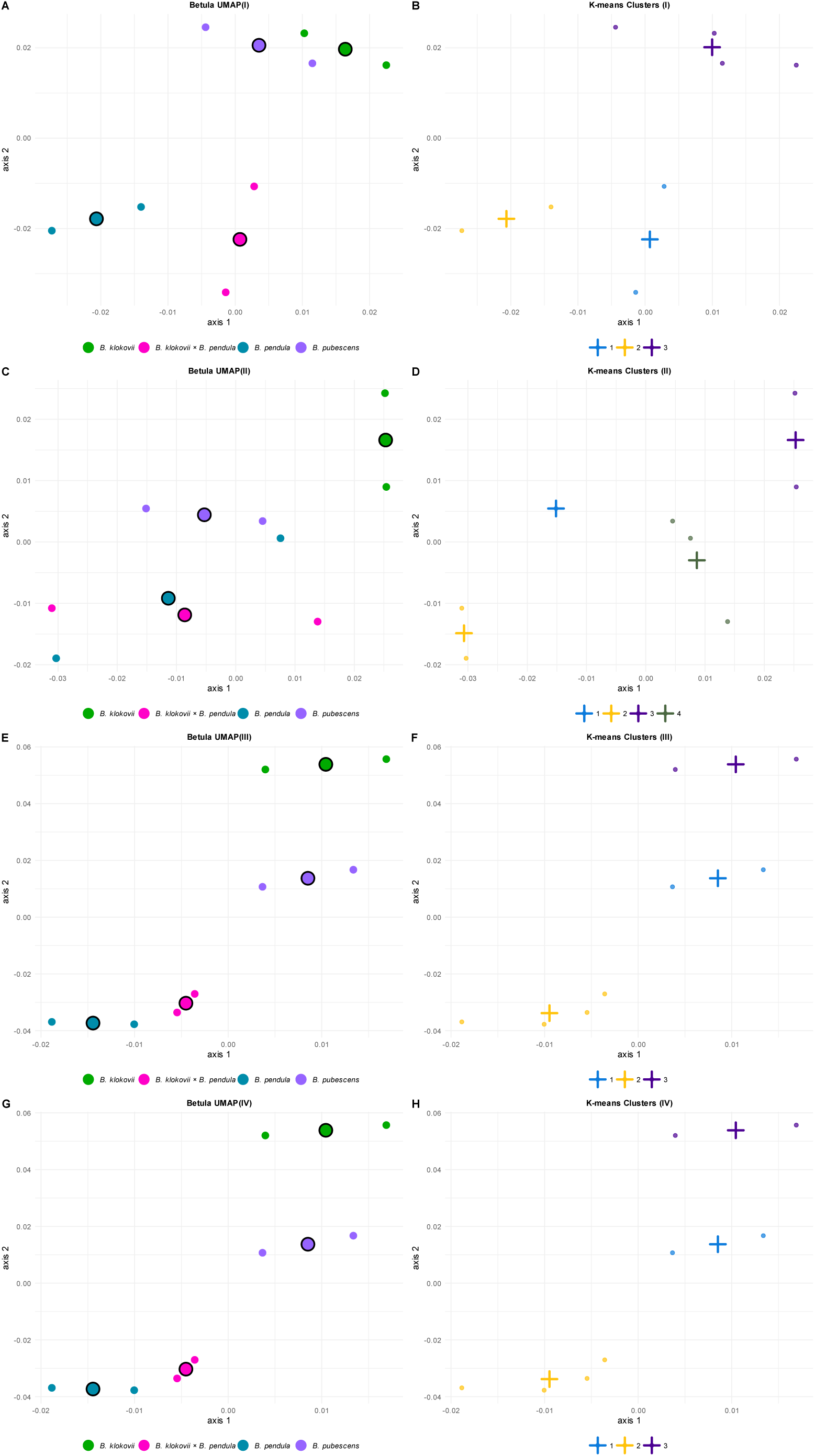
UMAP combined plot (K=3, n_epochs=1000 n_bootstraps=1000) based on genome-wide SNPs recovered under different mapping settings. Small circles represent individual samples; larger circles with black margins and crosses represent centroids. A (I) – UMAP graph based on genome-wide SNPs recovered from unfiltered reads mapped on *B. platyphylla* genome assembly using end-to-end bowtie2 alignment; B (I) – K-means clusters for UMAP based on genome-wide SNPs recovered from unfiltered reads mapped on *B. platyphylla* genome assembly using end-to-end bowtie2 alignment; C (II) – UMAP graph based on genome-wide SNPs recovered from unfiltered reads mapped on *B. platyphylla* genome assembly using local bowtie2 alignment; D (II) – K-means clusters for UMAP based on genome-wide SNPs recovered from unfiltered reads mapped on *B. platyphylla* genome assembly using local bowtie2 alignment; E (III) – UMAP graph based on genome-wide SNPs recovered from paired and unpaired reads after filtering for contamination mapped on *B. platyphylla* genome assembly using local bowtie2 alignment; F (III) – K-means clusters for UMAP based on genome-wide SNPs recovered from paired and unpaired reads after filtering for contamination mapped on *B. platyphylla* genome assembly using local bowtie2 alignment; G (IV) – UMAP graph based on genome-wide SNPs recovered from paired and unpaired reads after filtering for contamination and excluding unaligned ones mapped on *B. platyphylla* genome assembly using local bowtie2 alignment; H (IV) – K-means clusters for UMAP based on genome-wide SNPs recovered from paired and unpaired reads after filtering for contamination and excluding unaligned ones mapped on *B. platyphylla* genome assembly using local bowtie2 alignment.

Graphs based on SNPs recovered from mapping of the unfiltered reads based on end-to-end bowtie2 alignment (Fig.3A (I), S1–S3) show three clusters – one for *B. klokovii* and *B. pubescens* together, the second one for *B. pendula* and the third one for assumed hybrids *B. klokovii × B. pendula*.

UMAP graphs based on mapping the unfiltered reads using local bowtie2 alignment (Fig. 3C (II), S4–S6) show four clusters – a separate one for *B. klokovii*, another one with only one sample of *B. pubescens*, the third with one sample of *B. pendula*, and one sample of assumed hybrid *B. klokovii × B. pendula*. The last cluster contains single samples of *B. pubescens*, *B. pendula*, and assumed hybrids *B. klokovii × B. pendula*, respectively.

UMAP graphs based on SNPs recovered from mapping of reads after filtering for contamination using local bowtie2 alignment with paired and unpaired reads (Fig. 3E (III), S7–S9) and without unaligned reads (Fig. 3G (IV); S10–S12) are identical and show three clusters – two related clusters, *B. klokovii* and *B. pubescens*, and one for *B. pendula* together with assumed hybrids *B. klokovii × B. pendula.* We consider this clustering the most realistic from a biological point of view.

### Plastid genome assembly

Attempts to assemble plastid genomes using CLC genomic workbench resulted in contigs that were smaller than the average size of the complete plastid genome for birch. The details on assemblies are provided in Supporting information (Table S3).

In contrast, with NOVOPlasty we succeeded in obtaining six circularized plastid genome assemblies with k=23 and seven ones with k=33 and 45 (Supporting information, Table S4). The plastome assembly for sample 07 of downy birch was not recognized as circular by NOVOPlasty. However, the assembly contained a full-length contig that allowed us to proceed with it further. All plastome assemblies obtained with different k-mers were aligned together to check for consistency and fix ambiguous positions. The approach based on performing several separate assemblies with different k-mer lengths without and with different references for each sample is helpful for resolving ambiguous positions (Fig. 4).

**Fig.4.**
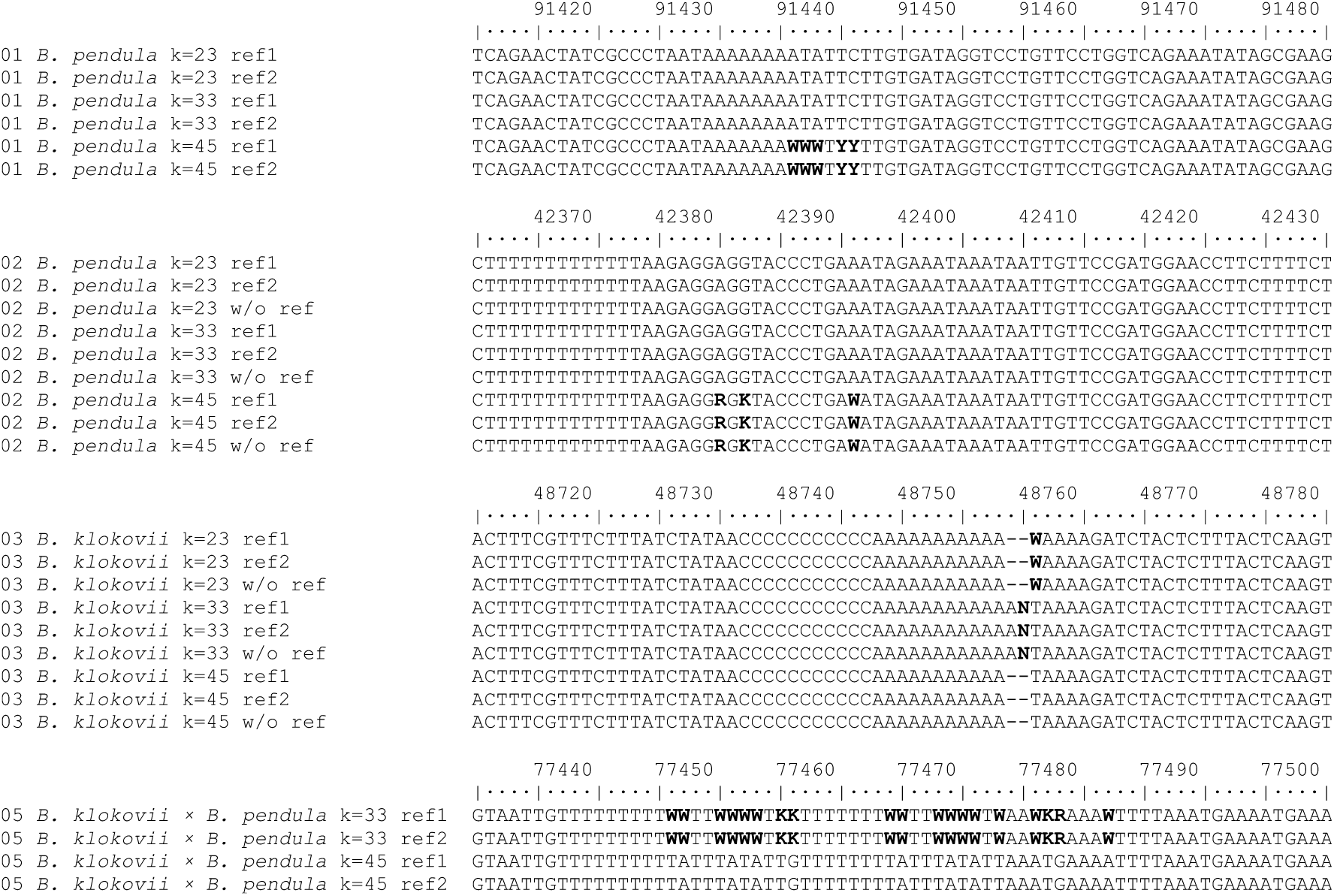
Presence of ambiguous positions (marked bold) in plastome assemblies depending on k-mer lengths k=23, k=33, and k=45.

As a result, we retrieved eight circularized consensus assemblies varying from 160,535 to 160,625 bp in length without ambiguous positions.

### Plastid genome annotations

Birch plastid genomes have a typical quadripartite structure containing two single-copy regions (small and large, respectively) and two inverted repeats (IRA and IRB) as in most angiosperms (Yang & al., 2019a, 2022). All newly produced assemblies were annotated using GeSeq followed by manual inspection/correction and contained 130 genes (113 unique), 85 CDSs, 8 rRNAs, 37 tRNAs, 49 exons (counted only for cases when one gene contains two or more exons separated by intron(s)), and 25 introns. In addition, two specific pseudogenes were detected: two copies of *ycf15* which was previously reported to be absent in Betulaceae (Yang & al., 2019a), and one or two copies *infA*.

Visualization of all annotated assemblies was performed in OGDraw 1.3.1 (Greiner & al., 2019); one for *B. klokovii* (sample 03, PP536053) is shown below (Fig. 5), all the rest are in the Supporting information (Fig. S13–S20).

**Fig. 5.**
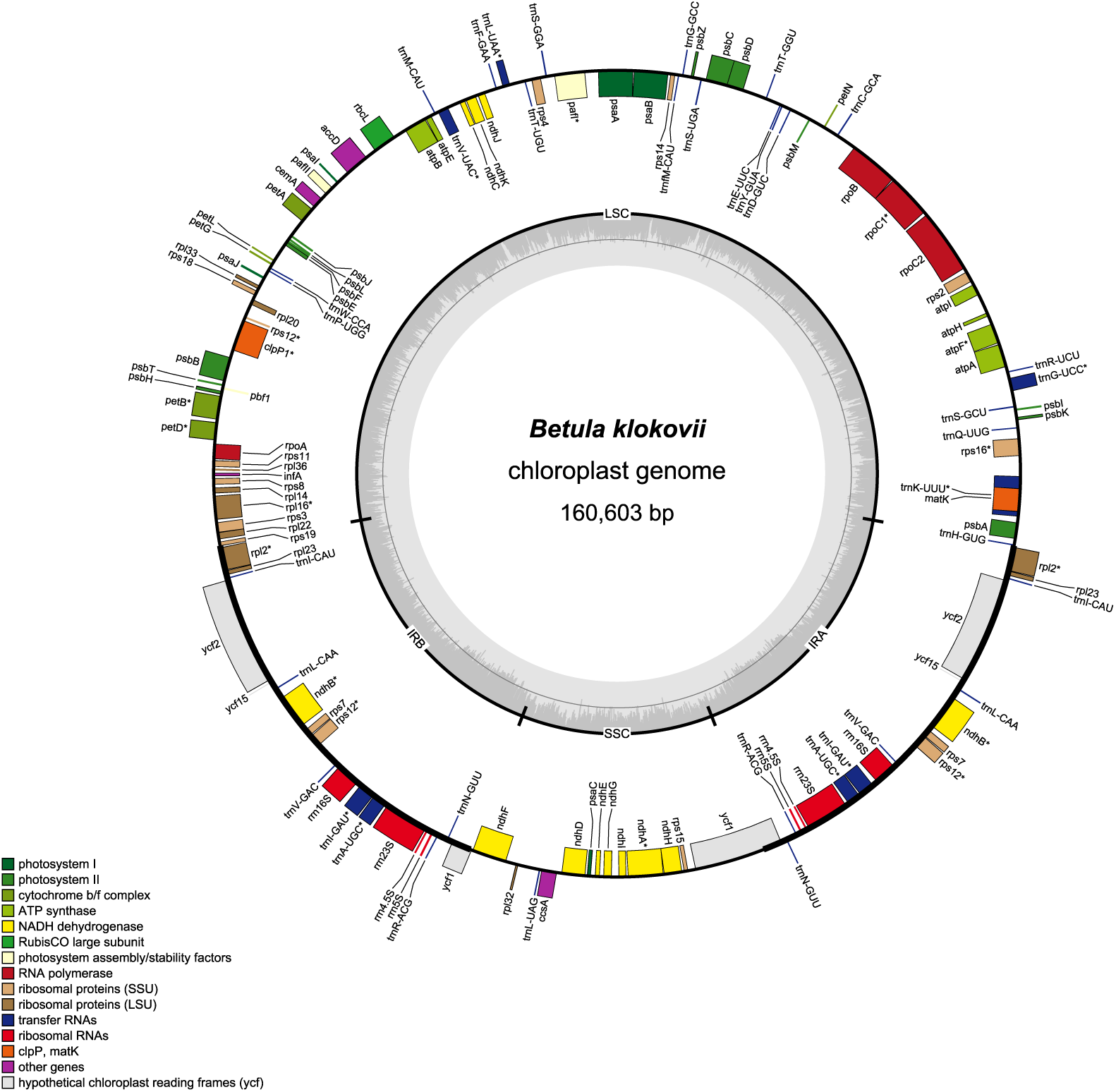
*Betula klokovii* plastome assembly (sample 03 (PP536053); annotation – GeSeq; visualization – OGDraw v1.3.1). The plot shows a circular organization of the plastome with two single-copy regions (large (LSC) and small (SSC), respectively) and two inverted repeats (IRA & IRB), and locations of the genes. Assemblies for all eight samples are in Supporting information, Fig. S13–S20.

### Plastome phylogeny

The topologies of Bayesian Inference (BI) phylogenetic trees based on plastome alignments with partitioning followed by model assignment according to AIC, AICc, and BIC were identical (Fig. 5, Supporting information, Fig. S22–S24, S38–S40). There is only one topological difference in comparison to the phylogenetic tree without partitioning which is not [highly] supported by posterior probability (Fig. 6, Supporting information, Fig. S21, S37).

**Fig. 6.**
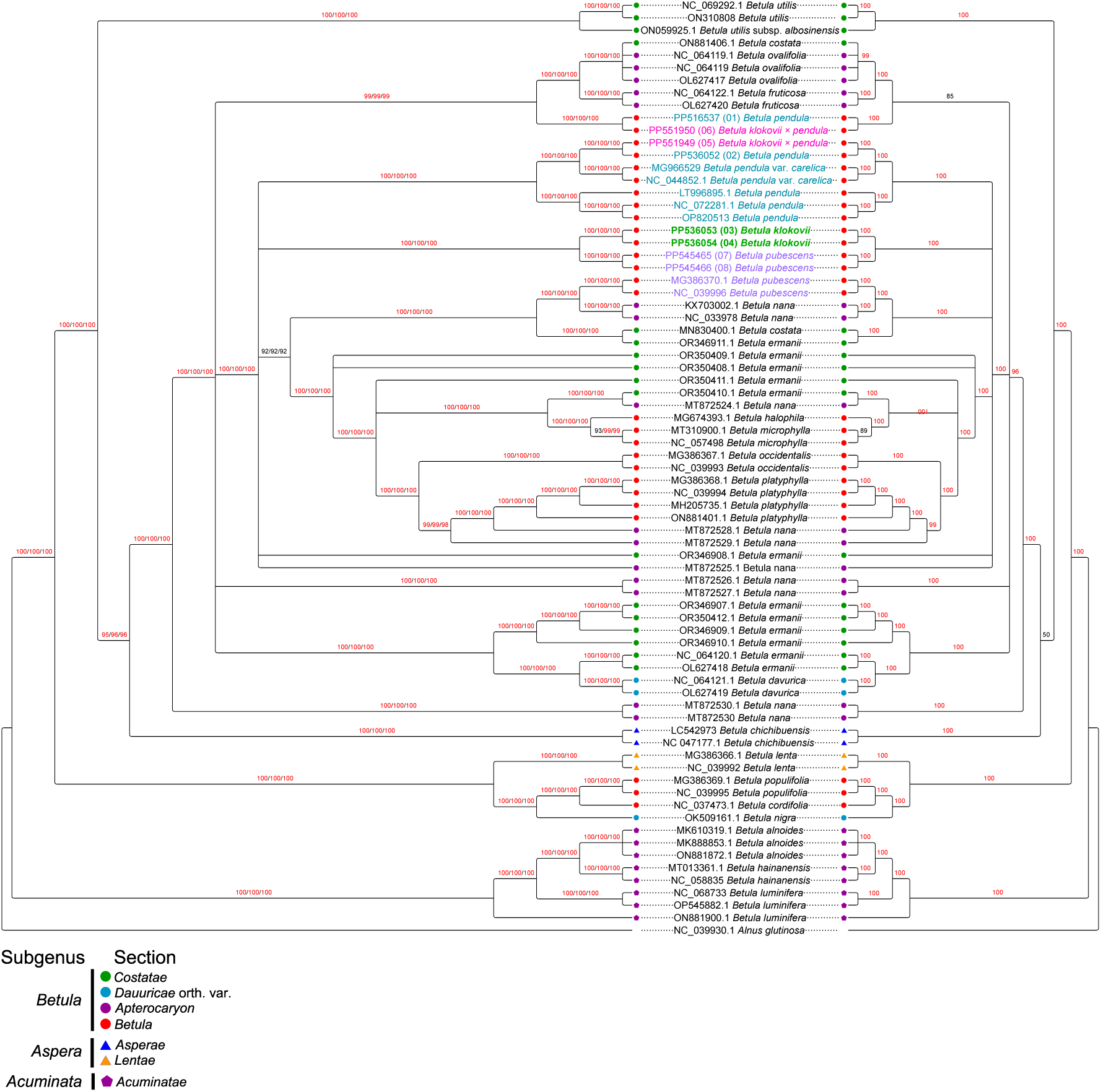
Combined plastome-based Bayesian Inference phylogeny reconstruction with (left, models assignment according to AIC, AICc, and BIC, respectively) and without partitioning (right, model GTR+I+G), 100,000,000 iterations with 10% burnin. MrBayes 3.2.6 high values of posterior probability (> 95%) are colored in red. *B. klokovii* is colored in green and written in bold, presumed *B. pendula* × *B. klokovii* hybrids are colored in magenta, *B. pubescens* is colored in violet, *B. pendula* and its intraspecific taxa are colored in cyan. Newly sequenced samples are indicated by numbers 01–08. Subgenera and sections are marked according to Wang & al., 2016.

The topologies of Maximum Likelihood (ML) phylogenetic trees were topologically slightly more diverse. Reconstructions based on partitioned alignment followed by model assignment according to AICc and BIC were topologically identical for RAxML (Supporting information, Fig. S27–S28, S43–S44) and both versions of IQTREE (Supporting information, Fig. S31–S32, S35–S36, S47–S48, S51–S52, S55–S56), while all those based either on a non-partitioned alignment (Supporting information, Fig. S21, S25, S29, S33, S37, S41, S45, S49, S53) or partitioned one according to AIC (Supporting information, Fig. S22, S26, S30, S34, S38, S42, S46, S50, S54) have minor topological differences.

The values of posterior probability for BI or bootstrap for ML are also slightly different between partitioned and non-partitioned data (Fig. 6; Supporting information, Fig. S21– S56) with a general tendency to be slightly higher for partitioned ones.

Removing duplicate/identical sequences from the alignment did not cause any substantial differences in the phylogeny reconstructions (Fig. S37–S56), and they followed the same pattern.

In general, phylogenetic reconstructions showed the majority of taxa were monophyletic, except *B. pednula, B. pubescens, B. nana, B. ermanii*, and *B. costata.* Also, there is a clear separation of subgenus *Acuminata* and section *Acuminatae* which were present as highly supported monophyletic basal clades in all cases. At the same time, there is no clear separation between subgenera *Aspera* and *Betula.* The same is also true for all the sections of these subgenera.

All phylogenetic reconstructions showed close relatedness of *B. klokovii* (PP536053 and PP536054) to Ukrainian samples of *B. pubescens* (PP545465 and PP545466). However, Klokov’s birch forms a separate highly supported monophyletic clade which is a sign of clear separation from downy birch. However, two other samples of *B. pubescens* MG386370.1 and NC_039996 from NCBI GenBank (which could be considered as one because the sequences are identical) group together with *B. nana* L., *B. ermanii* Cham., and *B. costata* Trautv. This could be explained by different factors: issues with the quality of plastome assemblies deposited into NCBI (in particular, MG386370.1 and NC_039996 assemblies have gaps and ambiguous sites), incorrect initial identification of the species/taxa, very wide consideration of *B. pubescens* (e.g., the entire *B. pubescens* aggr.), overlooked hybridization, or/and chloroplast capture, cryptic speciation, or the combination of different factors mentioned above.

Overall, the phylogeny based on whole plastomes is significantly better resolved in comparison to traditional markers such as internal transcribed spacers, SSRs, and individual plastid genes which is in agreement with previous studies (Wang & al., 2016; Yang & al., 2019a, 2022). However, the phylogeny remains not fully resolved. In particular, para-and polyphyly with different groupings are observed for different samples of *B. nana, B. costata, B. ermannii, B. pendula*, and *B. pubescens*.

Plastomes for *B. klokovii* were assembled for the first time. In comparison to all the other birch plastome sequences deposited in NCBI GenBank, they have a unique insert of 54bp “CACCTTTATTTTTGTGATTTAACATAGTTTAACATAATATTGATTTCTTACTTTT” between *psbB* and *psbD* genes (position 103390–103444 for PP536053 (sample 03) and position 13388–103442 for PP536054 (sample 04), respectively).

Phylogenetic networks demonstrate significant differences in number of reticulations depending on the method (Supporting information, Network S1–S12). However, all the networks show a similar pattern that is in agreement with other phylogenetic reconstructions. In particular, there is also a close relatedness with clear separation between *B. klokovii* and Ukrainian samples of *B. pubescens*.

## Discussion

Small isolated populations of endemic species/taxa could be a good model to study speciation processes. Small distribution areas, need for specific habitats, and low number of individuals make endemics much more vulnerable to climate change, human activities, and other impacting factors including irregular extreme events (Manes & al., 2021; Guo & al., 2023). According to the current assessment, over 17000 tree species are at risk due to the rapid global change (Boonman & al., 2024). At the same time, many of them are still lacking proper assessments and therefore considered data deficient (Boonman & al., 2024). The outcomes of detailed studies on rare endemics could also have practical applications in developing appropriate conservation measures, strategies, and policies.

In the current study, we attempted to clarify the phylogenetic relationships of critically endangered Ukrainian endemic birch *B. klokovii* with common species *B. pendula* and *B. pubescens* and with assumed hybrids *B. klokovii* × *B. pendula* based on LC-WGS. Besides, we aimed to test, whether herbarium material which was stored for a few years in normal conditions is suitable for such type of analysis.

The study reveals that birch herbarium specimens are suitable for a standard procedure of short-read whole genome sequencing. This is important since it enables sequencing of dried specimens from herbaria (Zeng & al., 2018; Raxworthy & Smith, 2021; Záveská Drábková, 2021). It also presents an approach for plastome assembly from unfiltered reads of LC-WGS. Although the current set of samples is limited, it provides new data on Ukrainian endemic birch species *B. klokovii* that could be useful for understanding its evolutionary history better and for developing better conservation or protection measures and policies for this species.

Low-coverage genome sequencing revealed some specific issues like high DNA fragmentation and imbalance between libraries and adaptors. This is most likely caused by using dried plant material (Raxworthy & Smith, 2021). Thus, in the case of scaling up the approach based on dried birch material, it would be reasonable to decrease the length of sequenced fragments from 2×300PE to 2×150PE. However, LC-WGS provided additional data and allowed to assemble complete plastid genomes from unfiltered reads using NOVOPlasty.

We were also able to recover genome-wide SNP data from LC-WGS. However, it was done only with mapping the reads to the *B. platyphylla* genome assembly (as reference) since this is the only chromosome-level genome assembly that is publicly available for birch. This poses certain limitations but is still reasonable in the context of phylogeny. UMAP graphs demonstrate a certain degree of isolation for *B. klokovii* from all the other studied birch taxa. At the same time, it reveals a very close placement of *B. pendula* and assumed hybrid *B. klokovii* × *B. pendula*. These were placed in different clusters only in the case of end-to-end alignment performed on reads without filtering for contamination. In our opinion, the best resolution from the biological point of view is provided by the graphs based on reads that were filtered for contamination together with “local” alignment option of bowtie2. These graphs are also in agreement with the outcome of the plastome-based phylogeny. The major difference is likely caused by the filtering for contamination although it removes not only contamination but also a part of plastid and mitochondrial reads. To get detailed better resolution there is a need to include more specimens and perhaps to perform a deeper sequencing.

Plastomes across Betulaceae (and *Betula* L. genus in particular) are highly conserved in genome size, gene order, structure, and have typical quadripartite structure as in most angiosperms (Yang & al., 2019a, 2022). Birch plastomes deposited in NCBI and included in our study vary in length from 158,647 (MG386370.1 and NC_39996 *B. pubescens*) to 161,349 bp (MG386368.1 and NC_039994.1 *B. platyphylla*). GC content varies from 35.9% (MT013361 and NC_058835 *B. hahaiensis*) to 36.4% (MG386370.1 and NC_39996 *B. pubescens*). Newly assembled plastomes vary in size from 160,535 (PP536052 (sample 02) *B. pendula*) to 160,625 bp (PP545465 (sample 07) *B. pubescens*), which is in the range of plastid genome sizes from NCBI. The GC content for all originally sequenced and assembled plastomes is 36.1% which is within the average range for birch plastomes. GC content could have an impact on both genome functioning and species ecology (Šmarda & Bureš, 2012; Šmarda & al., 2014). For example, in plastomes a correlation between CG content and accumulation of deleterious mutations is reported (Yu & al., 2021). At the same time, the GC content is reported to be relatively stable on the low taxonomic levels (family and below) (Šmarda & Bureš, 2012; Wu & al., 2024).

In comparison to downy birch plastome assemblies deposited in NCBI Gen Bank (MG386370.1 and NC_39996), newly sequenced and assembled ones for *B. pubescens* (PP545465 and PP545466) are 1,978–1,974 bp longer and less GC-rich. The potential cause for such a difference could be either an issue with older assemblies or, for example, initial misidentification, overlooked hybridization etc. It needs to be mentioned that some plastome assemblies deposited in NCBI GenBank (including reference ones) have certain issues (e.g., gaps, ambiguous nucleotides, etc.) that could have an impact on the phylogeny and make it harder to reconstruct. In addition, some variation in the size of plastomes among closely related species could be caused by variation in non-coding plastome regions (Karbstein & al., 2023; Long & al., 2024).

Although the current set of samples is limited, it provided insights into the taxonomic position of *B. klokovii* and also its presumable hybrids. Based on currently available data, *B. klokovii* is monophyletic and highly supported in all phylogenetic reconstructions. Therefore, it could be considered a separate species according to some species concepts, in particular the general linage concept (De Queiroz, 1998, 1999, 2007) and phylogenetic species concept (Donoghue, 1985).

*B. klokovii* is closely related to Ukrainian specimens of *B. pubescens* (PP545465 and PP545466) but at the same time is clearly separated from the latter. However, in our plastome-based phylogenetic reconstructions *B. pubescens* is not monophyletic since two samples from NCBI (MG386370.1 and NC_039996) cluster together with *B. nana, B. ermannii*, and *B. costata.* This could be due to different causes (see above): issues with the quality of plastome assemblies deposited into NCBI (in particular, MG386370.1 and NC_039996 assemblies have gaps and ambiguous sites), incorrect initial identification of the species/taxa, overlooked hybridization, or/and chloroplast capture, or the combination of these factors. In general, *B. pubescens* is a very complex and polymorphic group, and many more high-quality plastome assemblies are required to infer its structure. The same explanation also applies to several other cases of polytomy and uncertain placements (in particular for *B. nana, B. ermannii, and B. pendula*).

Based on currently available data we assume that *B. klokovii* is a separate taxon. The taxonomic rank of this taxon now can be defined as a separate species according to the morphological/phenetic and phylogenetic species concepts (Donoghue, 1985; De Queiroz, 1998, 1999, 2007). However, there is a need for a more precise rank identification that requires a larger study with more samples. At the same time, there is an ongoing debate on what should be considered a species in the case of *Betula* L. genus (Ashburner & McAllister, 2016; Wang & al., 2022). Prospective hybrids cluster together with *B. pendula* samples from the same geographic origin which indicates that they are either no hybrids or *B. pendula* was the maternal parent during hybridization. To clarify this aspect further there is a need for more samples and additional analyses that involve not only plastomes but nuclear genome data (e.g., phylogenetic networks reconstruction based on nuclear, plastid, and mitochondrial genomes).

Plastome phylogeny is in general better resolved in comparison with phylogenies based on a few genetic markers, which is in agreement with previous studies (Wang & al., 2016; Yang & al., 2019a, 2022). There is also a study arguing that reliable plastome-based phylogeny for birch is not possible due to a comparably low level of genetic differences among species (Shestibratov & al., 2021). However, the phylogenetic resolution of plastome-based trees also depends on the quality of plastome assemblies used for phylogenetic reconstructions and the frequency of interspecific hybridization. Also, there are known cases of conflicting phylogenies based on plastomes and nuclear genomes (Gitzendanner & al., 2018). Thus, phylogenetic trees based on plastomes should be complemented with trees constructed from nuclear or genome-wide markers.

The phylogenetic part of the present study includes more plastome assemblies from different taxa in comparison to previous ones (Wang & al., 2016; Yang & al., 2019a, 2022). It should be also noticed that most published studies on birch plastomes are focused on one or a few species (Hu & al., 2017; Wang & al., 2019; Yin & al., 2019, 2020; Hua & Zhu, 2020; Yoshida & al., 2020; Shestibratov & al., 2021; Zhang & al., 2023; Worth & al., 2023).

Plastome-based phylogeny has its benefits and limitations. The benefits are that chloroplast DNA is non-recombining which allows treating the entire plastome as a single locus, which reduces population size, and makes it possible to recover ancestral lineages (Palmé & al., 2003). In addition, haploid plastid genomes make it easier to perform a phylogenetic analysis involving samples with different ploidies (Gitzendanner & al., 2018), which is highly relevant for the genus *Betula* L.

The limitations for plastome-based phylogenetic reconstructions: in birch like in other angiosperms, plastids are haploid and maternally inherited (Tutin, 1993; Tsuda & Ide, 2010). Therefore, plastomes alone are not well-suited for detecting hybridization. At the same time, plastomes are more prone to stochastic events and there are known cases of chloroplast capture (Thomson & al., 2015a; Meucci & al., 2021; Yang & al., 2021; Touchette & al., 2024) that could have an impact on the phylogeny. Besides, low variability of plastid genomes could result in unresolved phylogenies (polytomies).

Plastome phylogeny reconstructions showed significant stability in topology (single or multiple models + different optimization criteria). Differences in the topology of ML trees were dependent on the method (in particular, there are different algorithms for bootstrap computation in RAxML and IQ-TREE) and whether the data were partitioned or not. However, the placement of *B. klokovii* was the same in all cases.

For more precise estimations of phylogeny and also to define the taxonomic rank of *B. klokovii*, additional research using plastomes and genome-wide markers and WGS is needed with more samples of *B. klokovii* and related species to be analyzed. At the same time, there is evidence that genus-wide genomic data and separation of subgenomes for polyploid species provide a significantly better resolved phylogeny for birch (Wang & al., 2021). Therefore, one of the best approaches would be to assemble reference-grade genomes (the details see Lawniczak & al., 2022; Mc Cartney & al., 2024) using long-read sequencing (e.g., ONT or PacBio HiFi) together with chromatin conformation capturing techniques (e.g., Hi-C, Omni-C, Pore-C) (Pucker & al., 2022; Karbstein & al., 2023) for several birch species including *B. klokovii.* This is especially important since there are no genome assemblies for polyploid birch species. In addition, all three currently available genome assemblies for birch do not match the modern criteria for reference-grade genome assemblies (Lawniczak & al., 2022; Mc Cartney & al., 2024). Whole genome sequences will provide detailed information on genome-wide differentiation, the role of polyploidization, and additional information about epigenetic modifications. Also, it will allow more precise reconstruction of phylogenetic relationships since it is known that the choice of the reference genome and filtering could have an impact on phylogeny (Thorburn & al., 2023; Rick & al., 2024). In addition, assembling reference genomes for critically endangered endemic birch species could have practical outcomes for conservation (Formenti & al., 2022; Theissinger & al., 2023; McCartney & al., 2024).

## Conclusion

Based on available data, *Betula klokovii* is likely a genetically distinct, separate taxon related to *Betula pubescens.* Highly supported separation from *B. pubescens* is present in all phylogenetic reconstructions and also on UMAP graphs based on genome-wide variant analysis.

SNP data recovered from LC-WGS and analyzed by non-linear dimension reduction approach UMAP show a certain degree of separation for *B. klokovii* from all the other studied birch taxa. The best taxonomic resolution was achieved with mapping filtered reads using local alignment.

Plastome-based phylogeny provides a better resolution of closely related birch taxa in comparison to traditional phylogenetic markers such as ITS.

Partitioning followed by an assignment of different models has minor or no impact on tree topology and support values in comparison to reconstructions obtained from non-partitioned data with a single model.

To resolve the status of *B. klokovii* and to estimate the level of its hybridization better, there is a need to sequence much more samples of this birch (ideally – the whole population) and also related species and prospective hybrids from close locations based on chloroplast and nuclear genomes.

Performing high-quality reference-grade whole-genome sequencing and genome assembly would be beneficial to address taxonomic problems with *B. klokovii* and related taxa, to better understand processes of speciation in birch and to develop more efficient conservation measures for critically endangered birch species.

## Acknowledgements

Andrii Shtogun, Dr. Mykola Shtogryn, Iryna Dovhaniuk, and other staff members of NNP “Kremenets Mountains” for help in collecting birch specimens

Prof. Dr. Sergei Mosyakin, Dr. Nataliia Shyan, and Dr. Igor Olshanskyi for taxonomic consultations and access to specimens (including type specimens) deposited in the National Herbarium of Ukraine (KW)

Prof. Dr. Elvira Hörandl for taxonomic consultations and suggestions. Dr. Víctor Chano for consultations/advice regarding plastid assemblies.

Oleksandra Dolynska for technical support and assistance during lab work.

Competence Centre for Genomic Analysis (CCGA) Kiel within the DFG competence network GWDG for providing computational resources for providing sequencing services for our pilot project; Dr. Janina Fuß and Dr. Sören Franzenburg for providing technical support and consultations regarding low-coverage sequencing.

This work used the Scientific Compute Cluster at GWDG, the joint data center of Max Planck Society for the Advancement of Science (MPG) and University of Göttingen.

## Data availability

Plastome assemblies generated within this study are deposited to NCBI GenBank (accessions PP516537, PP536052, PP536053, PP536054, PP545465, PP545466, PP551949, PP551950). Raw sequence reads are deposited to NCBI SRA BioProject PRJNA1088975.

Data on the performed analyses (alignments, partitioning, input files, bioinformatic scripts, etc.) is available as supporting information and placed on GitHub repository (https://github.com/AndriiTarieiev/Betula_klokovii_LC-WGS_data_analysis).

## Author contribution

AT – collected the material, took part in conceptualization, conducted lab work and bioinformatic analyses, wrote the first draft and edited the manuscript; KK – supported bioinformatic analyses and edited the manuscript; OG – conceptualization of the study, funding acquisition, edited the manuscript.

