## Supporting information - Figures S1-S56 for "Phylogenetic and taxonomic insights into *Betula*: low-coverage whole genome sequencing and plastome analysis with focus on the rare Ukrainian endemic species *Betula klokovii* Zaverucha"

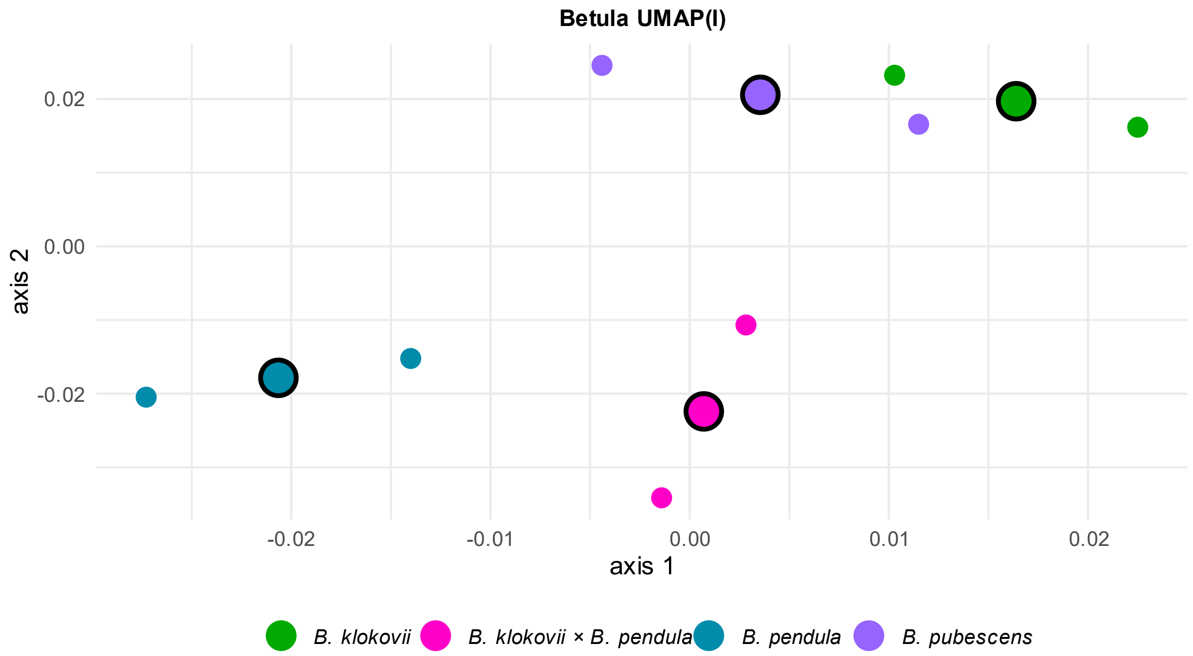


**Fig. S1** UMAP graph (3 neighbors, 2 components, 1000 epochs, 1000 bootstrap iterations) based on genome-wide SNPs recovered from unfiltered reads mapped on *B. platyphylla* genome assembly using end-to-end bowtie2 alignment (I)

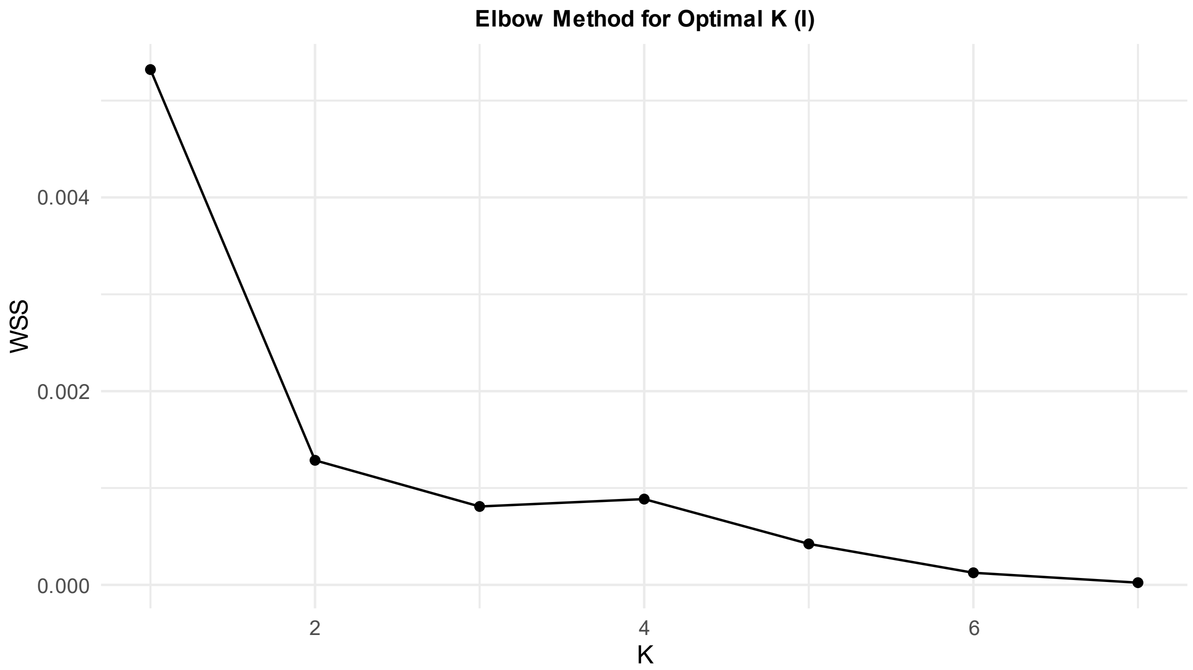


**Fig. S2** Elbow method for optimal K identification for UMAP (3 neighbors, 2 components, 1000 epochs, 1000 bootstrap iterations) based on genome-wide SNPs recovered from unfiltered reads mapped on *B. platyphylla* genome assembly using end-to-end bowtie2 alignment (I)

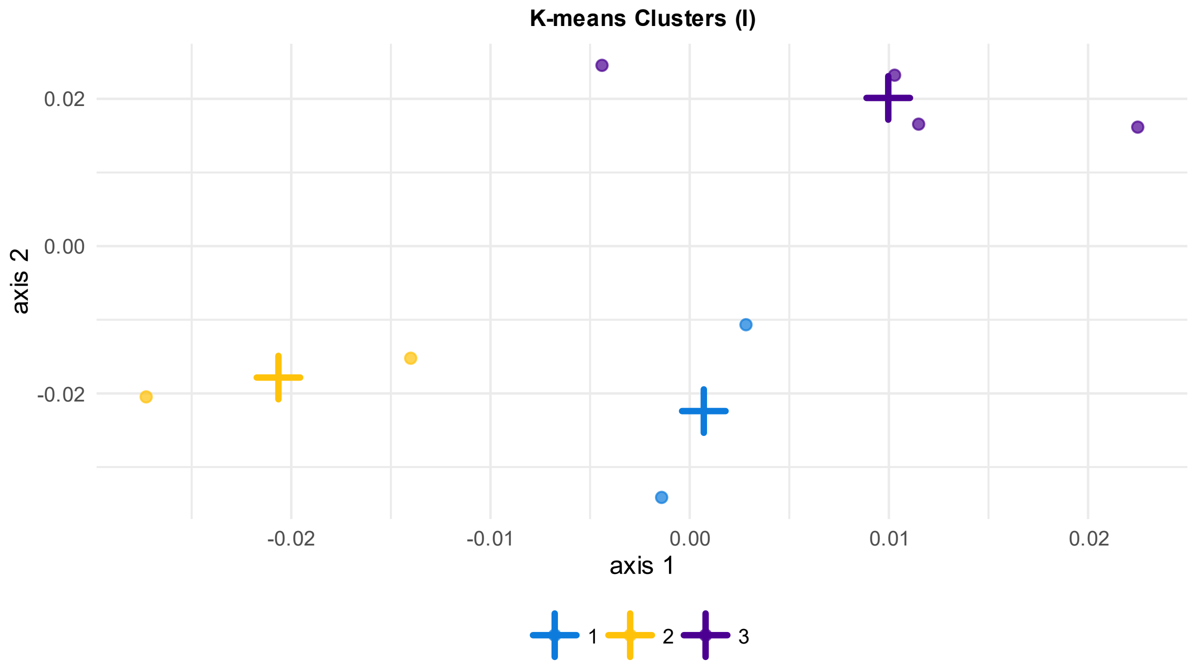


**Fig. S3** K-means clusters for UMAP (3 neighbors, 2 components, 1000 epochs, 1000 bootstrap iterations) based on genome-wide SNPs recovered from unfiltered reads mapped on *B. platyphylla* genome assembly using end-to-end bowtie2 alignment (I)

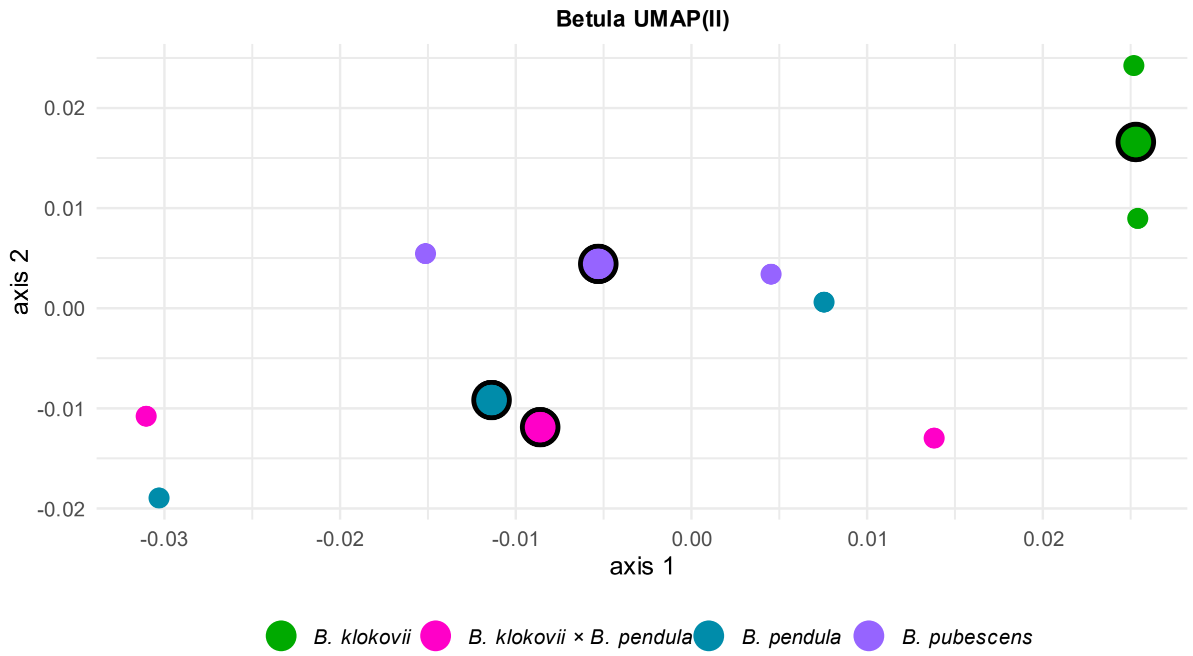


**Fig. S4** UMAP graph (3 neighbors, 2 components, 1000 epochs, 1000 bootstrap iterations) based on genome-wide SNPs recovered from unfiltered reads mapped on *B. platyphylla* genome assembly using local bowtie2 alignment (II)

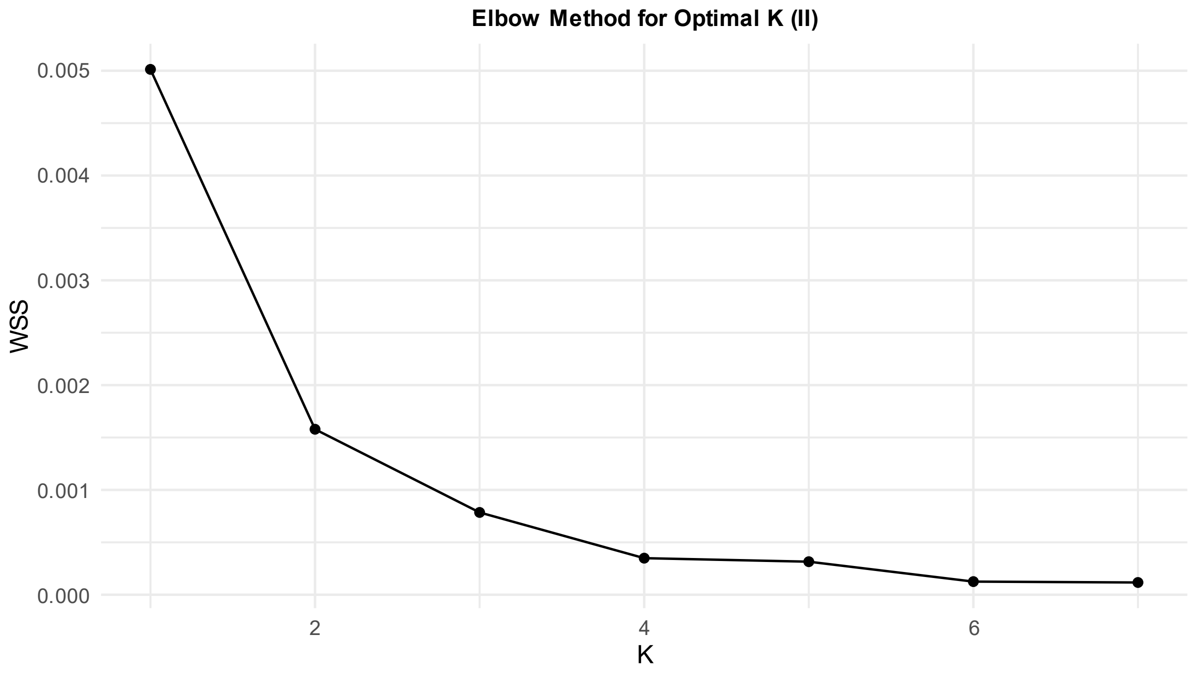


**Fig. S5** Elbow method for optimal K identification for UMAP (3 neighbors, 2 components, 1000 epochs, 1000 bootstrap iterations) based on genome-wide SNPs recovered from unfiltered reads mapped on *B. platyphylla* genome assembly using local bowtie2 alignment (II)

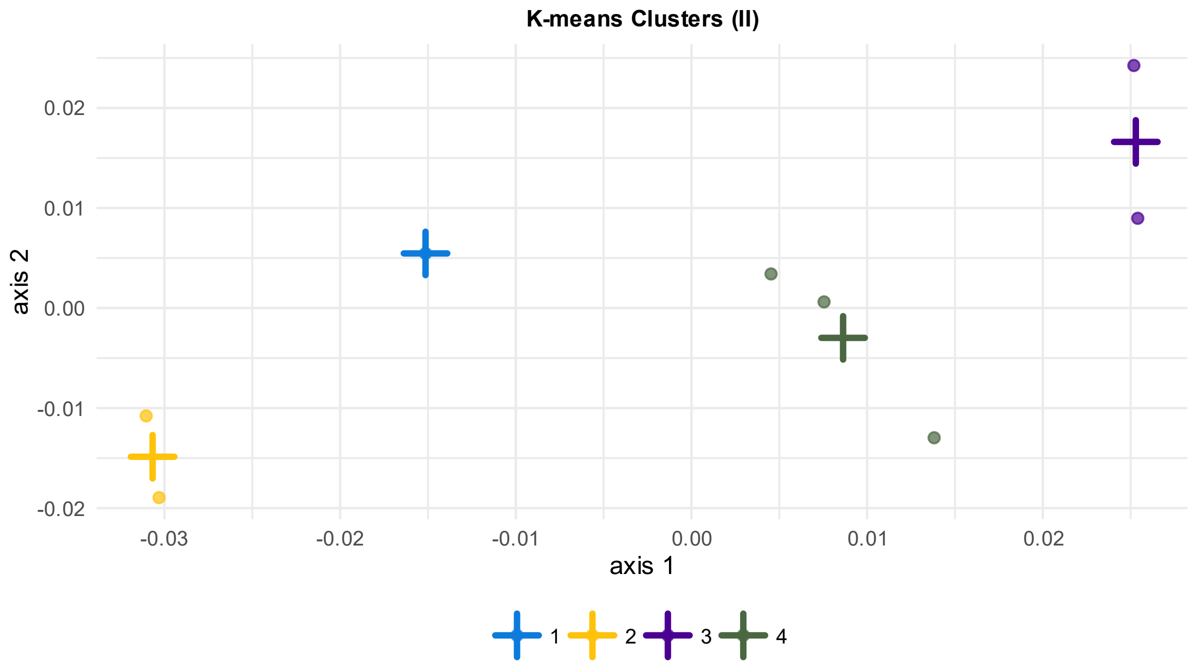


**Fig. S6** K-means clusters for UMAP (3 neighbors, 2 components, 1000 epochs, 1000 bootstrap iterations) based on genome-wide SNPs recovered from unfiltered reads mapped on *B. platyphylla* genome assembly using local bowtie2 alignment (II)

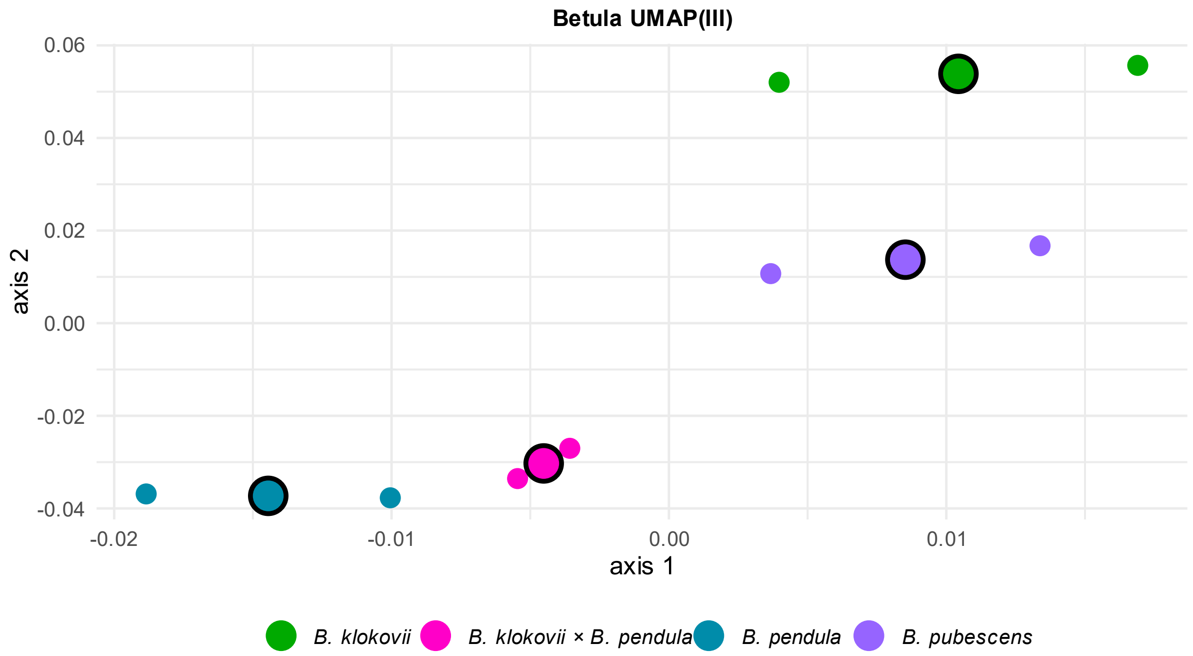


**Fig. S7** UMAP graph (3 neighbors, 2 components, 1000 epochs, 1000 bootstrap iterations) based on genome-wide SNPs recovered from paired and unpaired reads after filtering for contamination mapped on *B. platyphylla* genome assembly using local bowtie2 alignment (III)

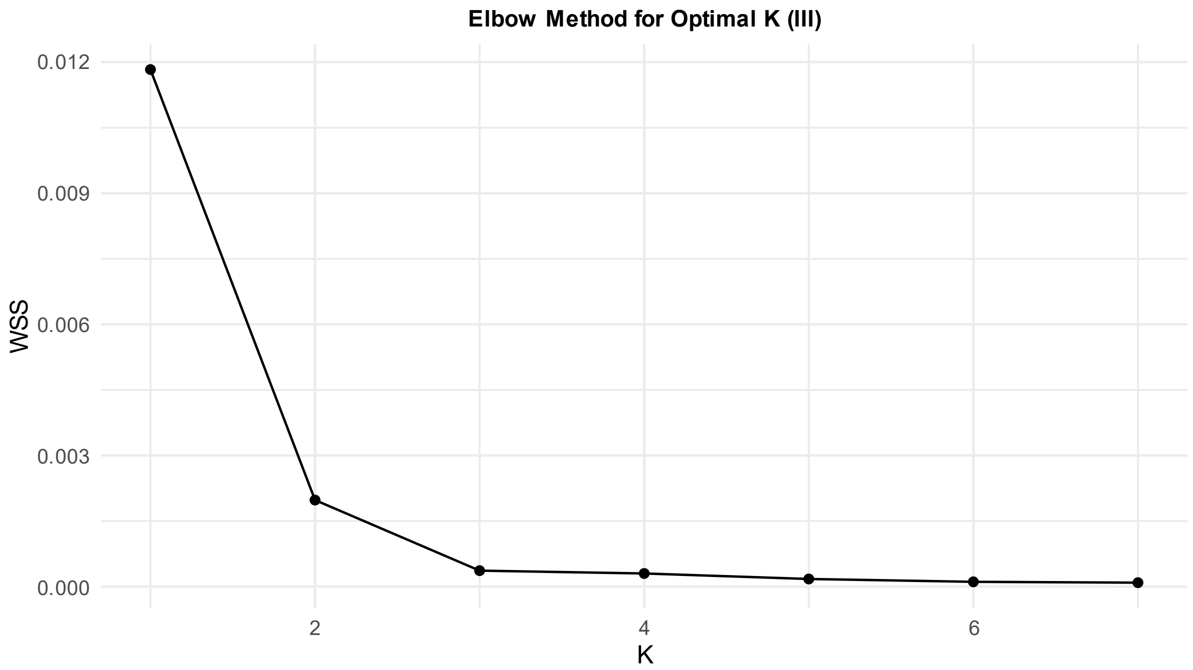


**Fig. S8** Elbow method for optimal K identification for UMAP (3 neighbors, 2 components, 1000 epochs, 1000 bootstrap iterations) based on genome-wide SNPs recovered from paired and unpaired reads after filtering for contamination mapped on *B. platyphylla* genome assembly using local bowtie2 alignment (III)

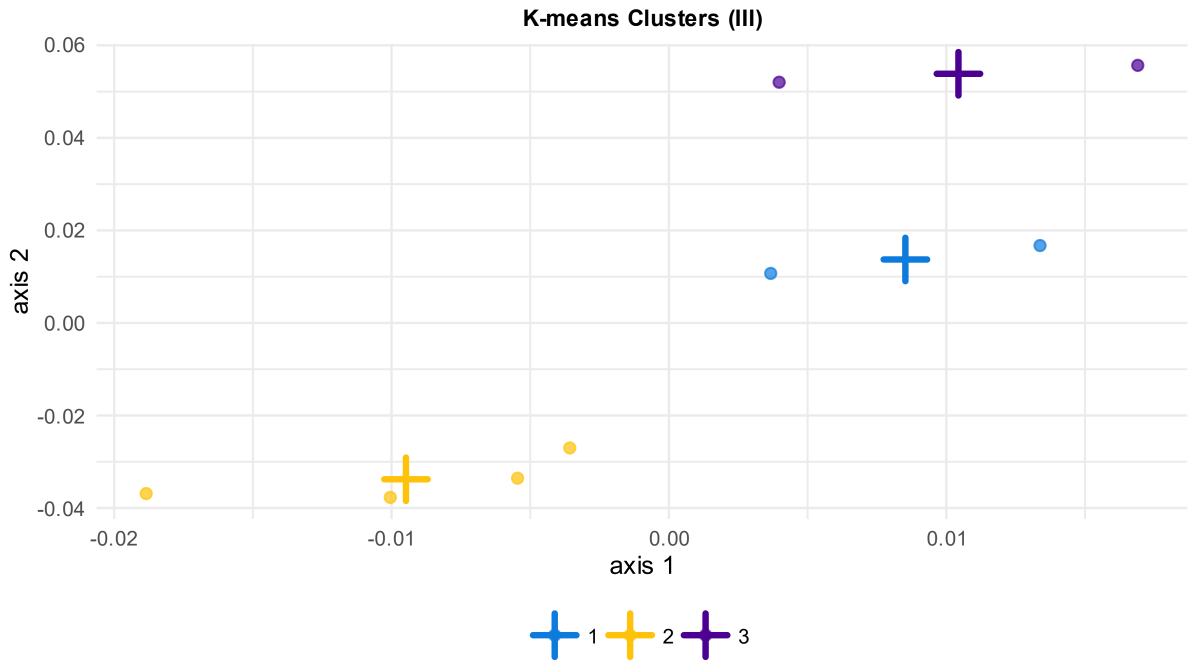


**Fig. S9** K-means clusters for UMAP (3 neighbors, 2 components, 1000 epochs, 1000 bootstrap iterations) based on genome-wide SNPs recovered from paired and unpaired reads after filtering for contamination mapped on *B. platyphylla* genome assembly using local bowtie2 alignment (III)

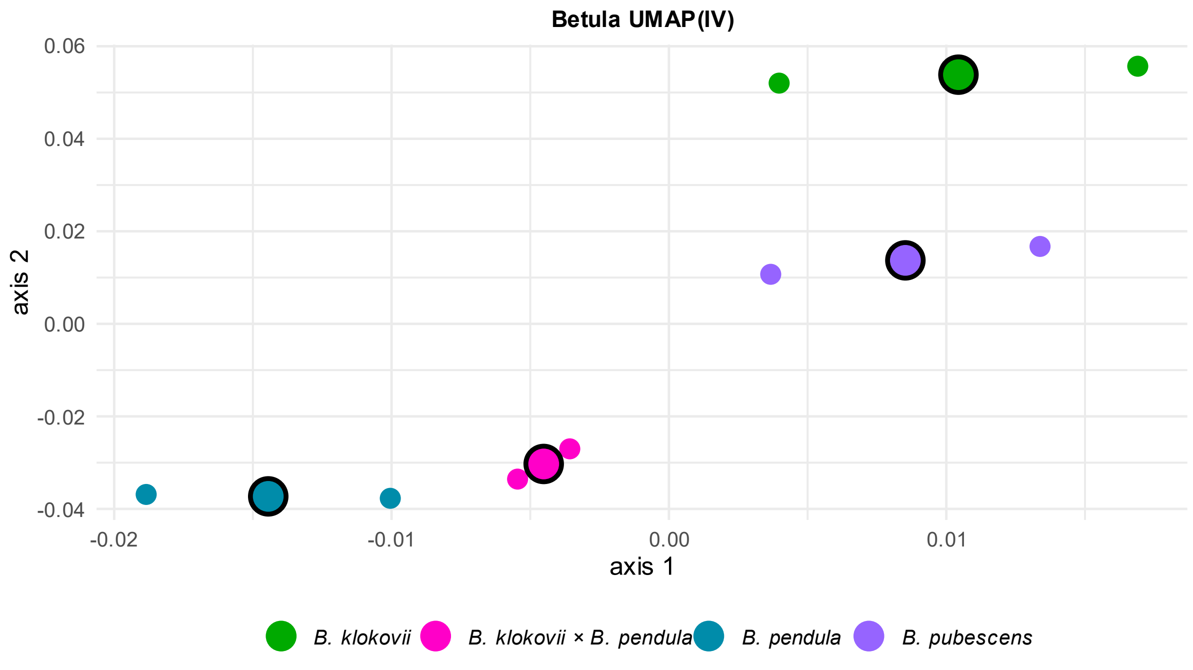


**Fig. S10** UMAP graph (3 neighbors, 2 components, 1000 epochs, 1000 bootstrap iterations) based on genome-wide SNPs recovered from paired and unpaired reads after filtering for contamination and excluding unaligned ones mapped on *B. platyphylla* genome assembly using local bowtie2 alignment (IV)

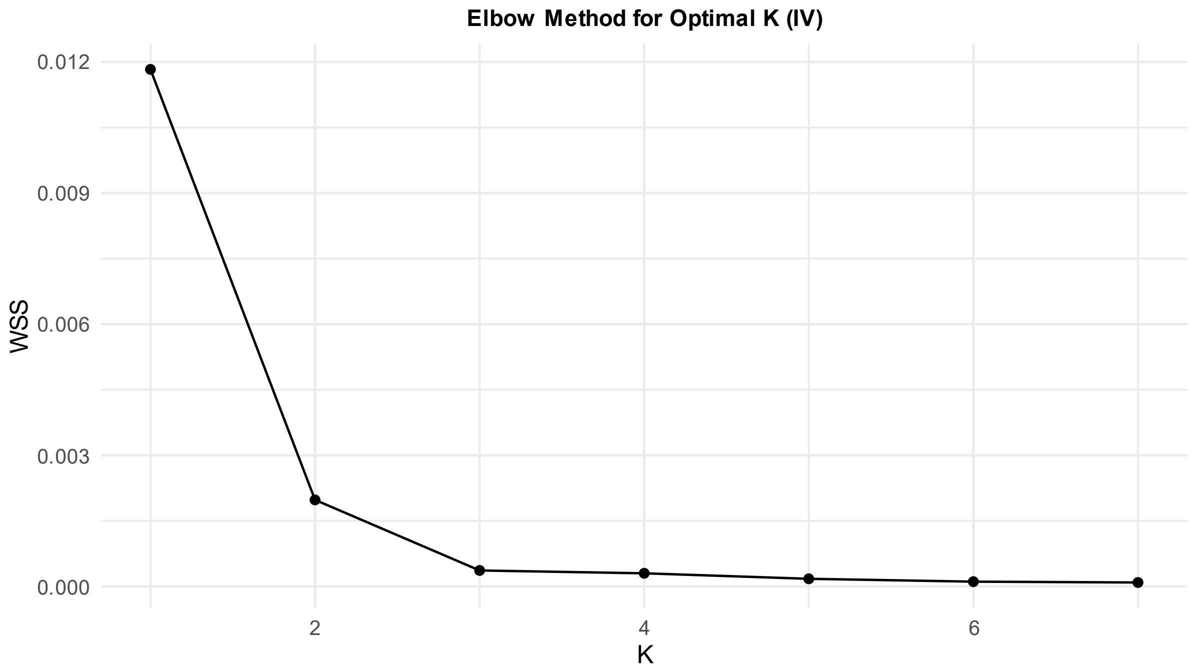


**Fig. S11** Elbow method for optimal K identification for UMAP (3 neighbors, 2 components, 1000 epochs, 1000 bootstrap iterations) based on genome-wide SNPs recovered from paired and unpaired reads after filtering for contamination and excluding unaligned ones mapped on *B. platyphylla* genome assembly using local bowtie2 alignment (IV)

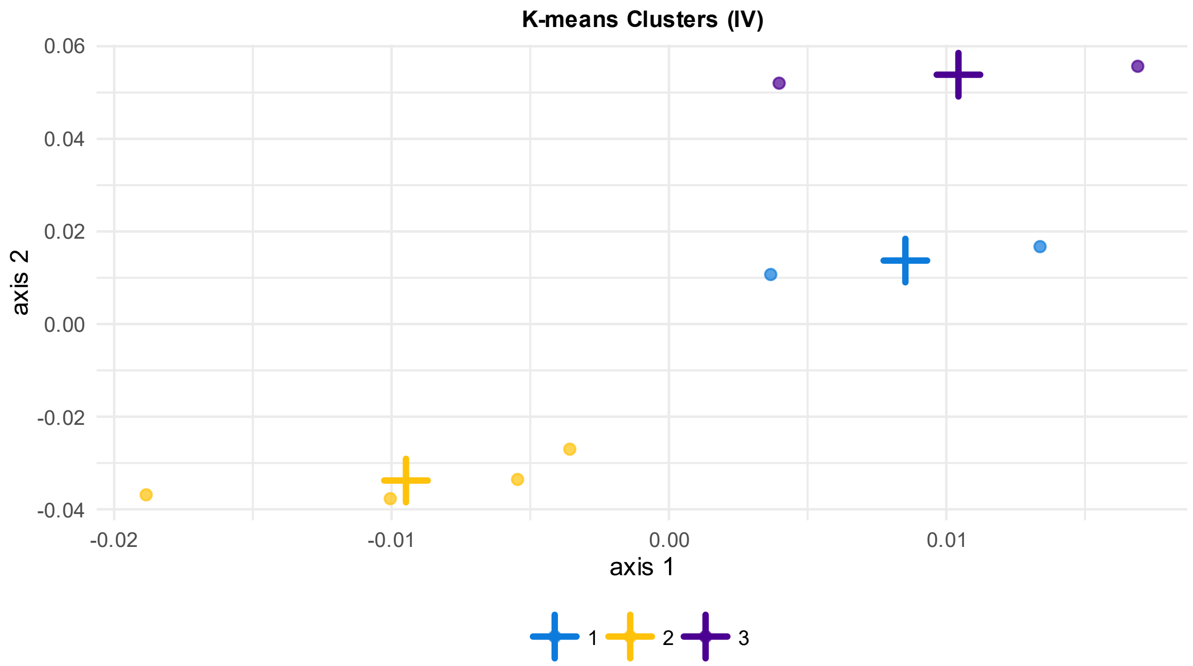


**Fig. S12** K-means clusters for UMAP (3 neighbors, 2 components, 1000 epochs, 1000 bootstrap iterations) based on genome-wide SNPs recovered from paired and unpaired reads after filtering for contamination and excluding unaligned ones mapped on *B. platyphylla* genome assembly using local bowtie2 alignment (IV)

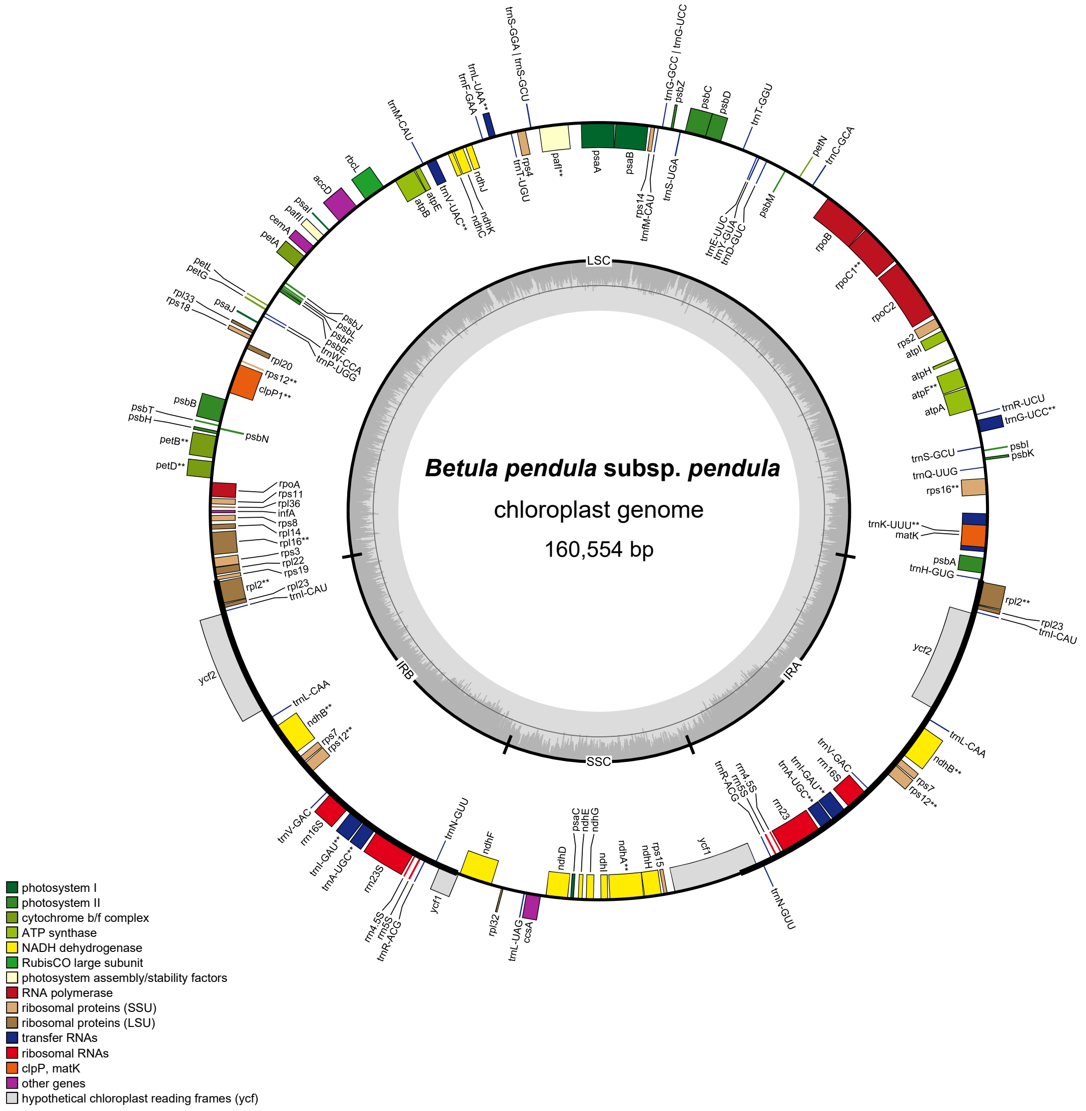


**Fig. S13** *Betula pendula* plastome assembly (sample 01 (PP516537); annotation – GeSeq; visualization – OGDraw v1.3.1).

The plot shows a circular organization of the plastome with two single-copy regions (large (LSC) and small (SSC), respectively) and two inverted repeats (IRA & IRB), and locations of the genes.

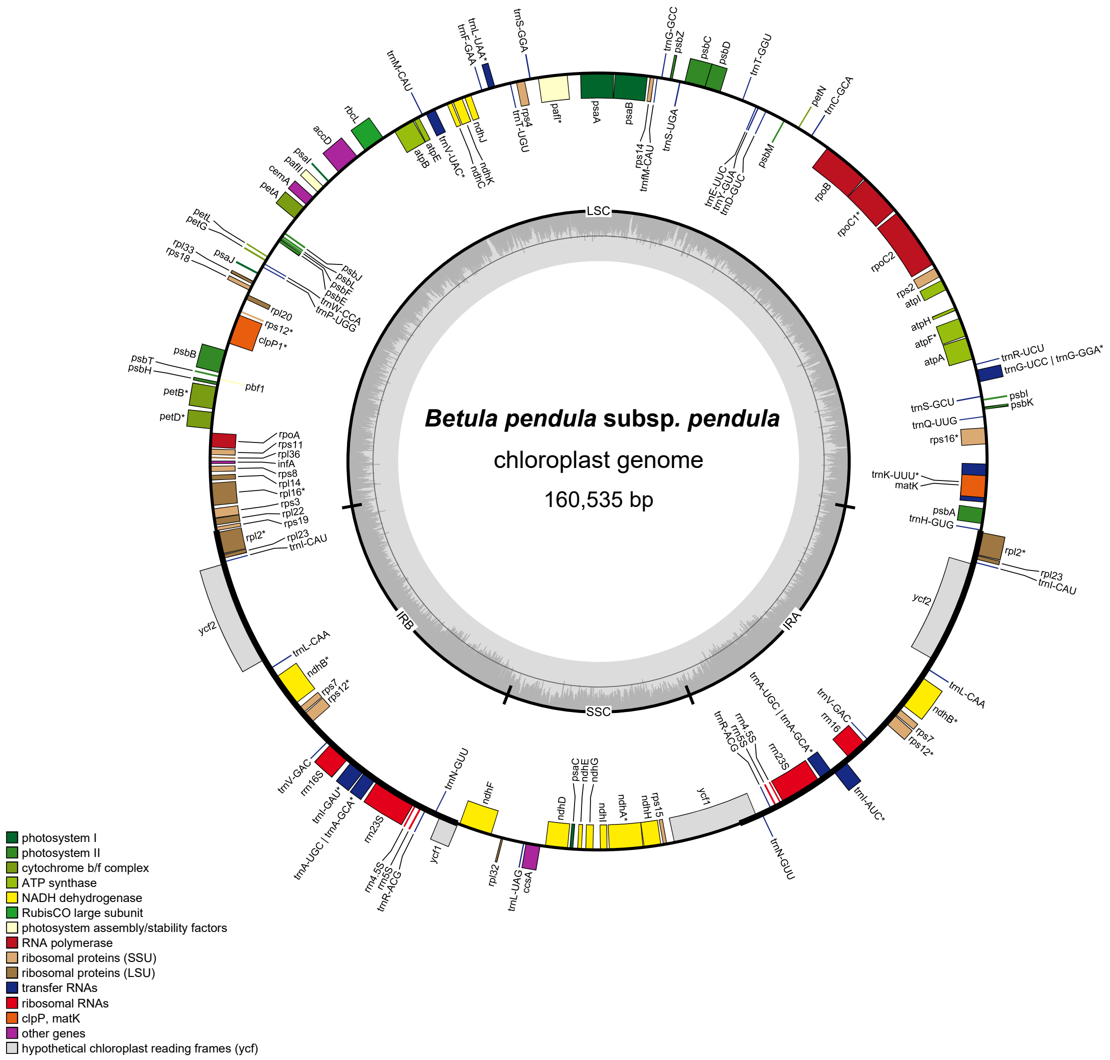


**Fig. S14** *Betula pendula* plastome assembly (sample 02 (PP536052); annotation – GeSeq; visualization – OGDraw v1.3.1).

The plot shows a circular organization of the plastome with two single-copy regions (large (LSC) and small (SSC), respectively) and two inverted repeats (IRA & IRB), and locations of the genes.

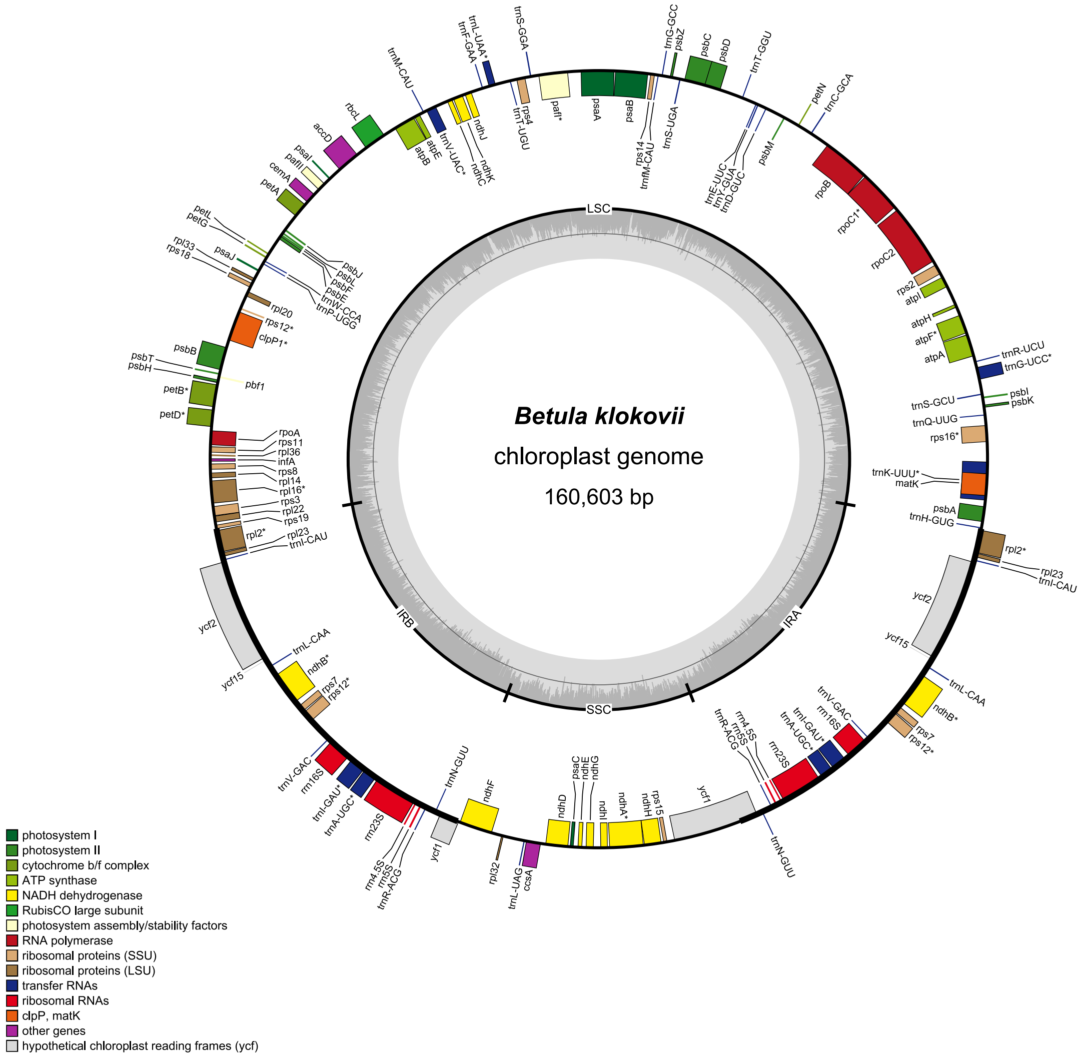


**Fig. S15** *Betula klokovii* plastome assembly (sample 03 (PP536053); annotation – GeSeq; visualization – OGDraw v1.3.1).

The plot shows a circular organization of the plastome with two single-copy regions (large (LSC) and small (SSC), respectively) and two inverted repeats (IRA & IRB), and locations of the genes.

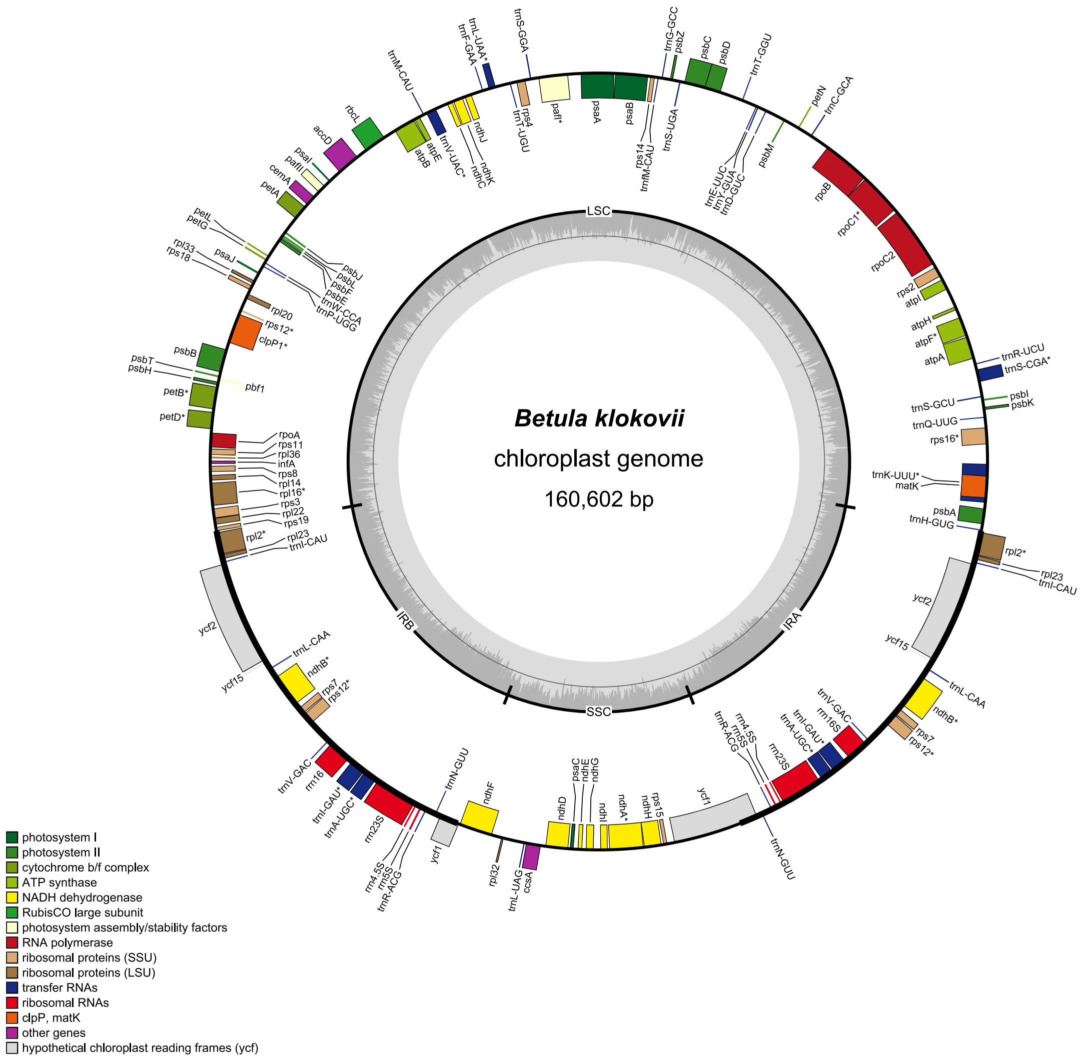


**Fig. S16** *Betula klokovii* plastome assembly (sample 04 (PP536054); annotation – GeSeq; visualization – OGDraw v1.3.1).

The plot shows a circular organization of the plastome with two single-copy regions (large (LSC) and small (SSC), respectively) and two inverted repeats (IRA & IRB), and locations of the genes.

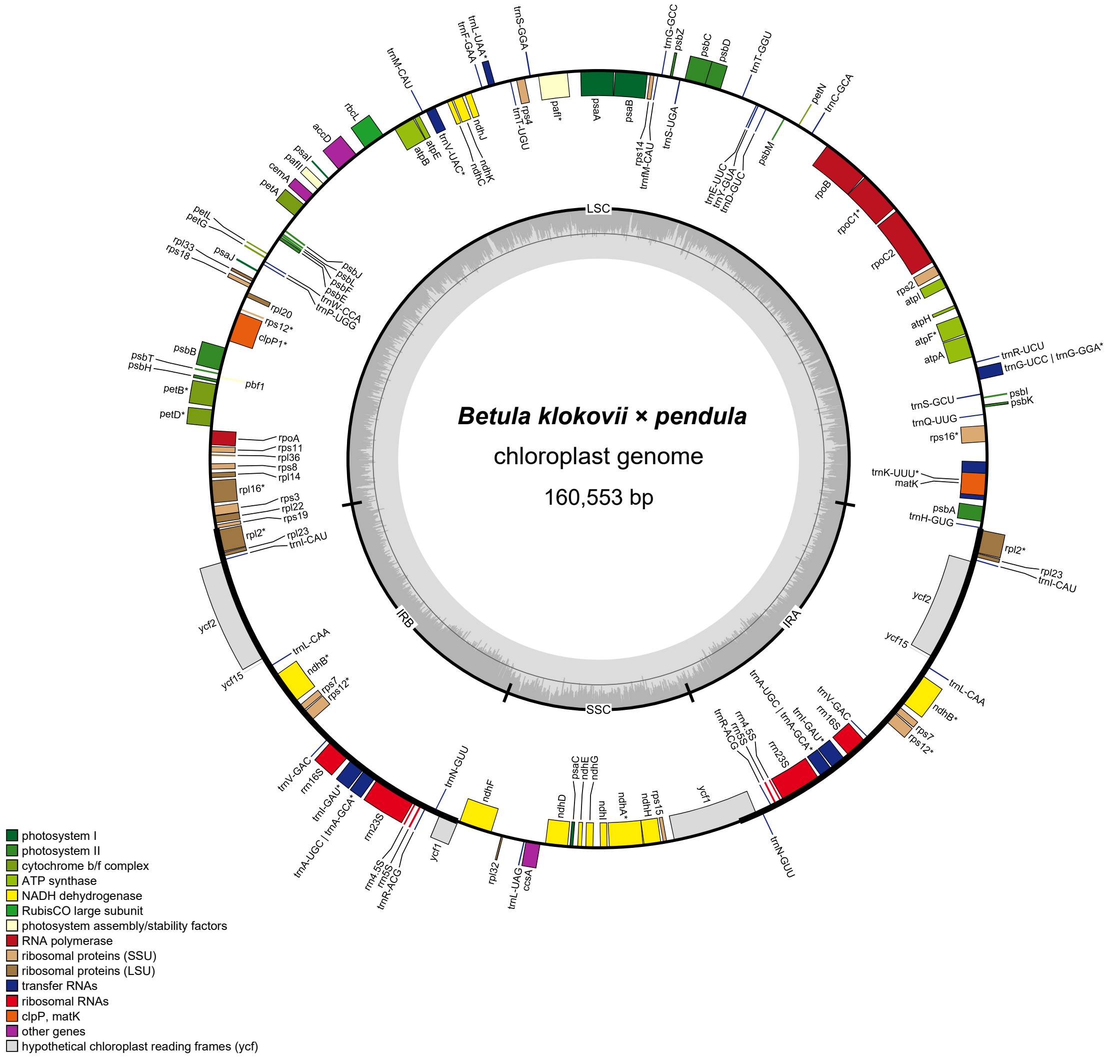


**Fig. S17** *Betula klokovii* × *Betula pendula* plastome assembly (sample 05 (PP551949); annotation – GeSeq; visualization – OGDraw v1.3.1).

The plot shows a circular organization of the plastome with two single-copy regions (large (LSC) and small (SSC), respectively) and two inverted repeats (IRA & IRB), and locations of the genes.

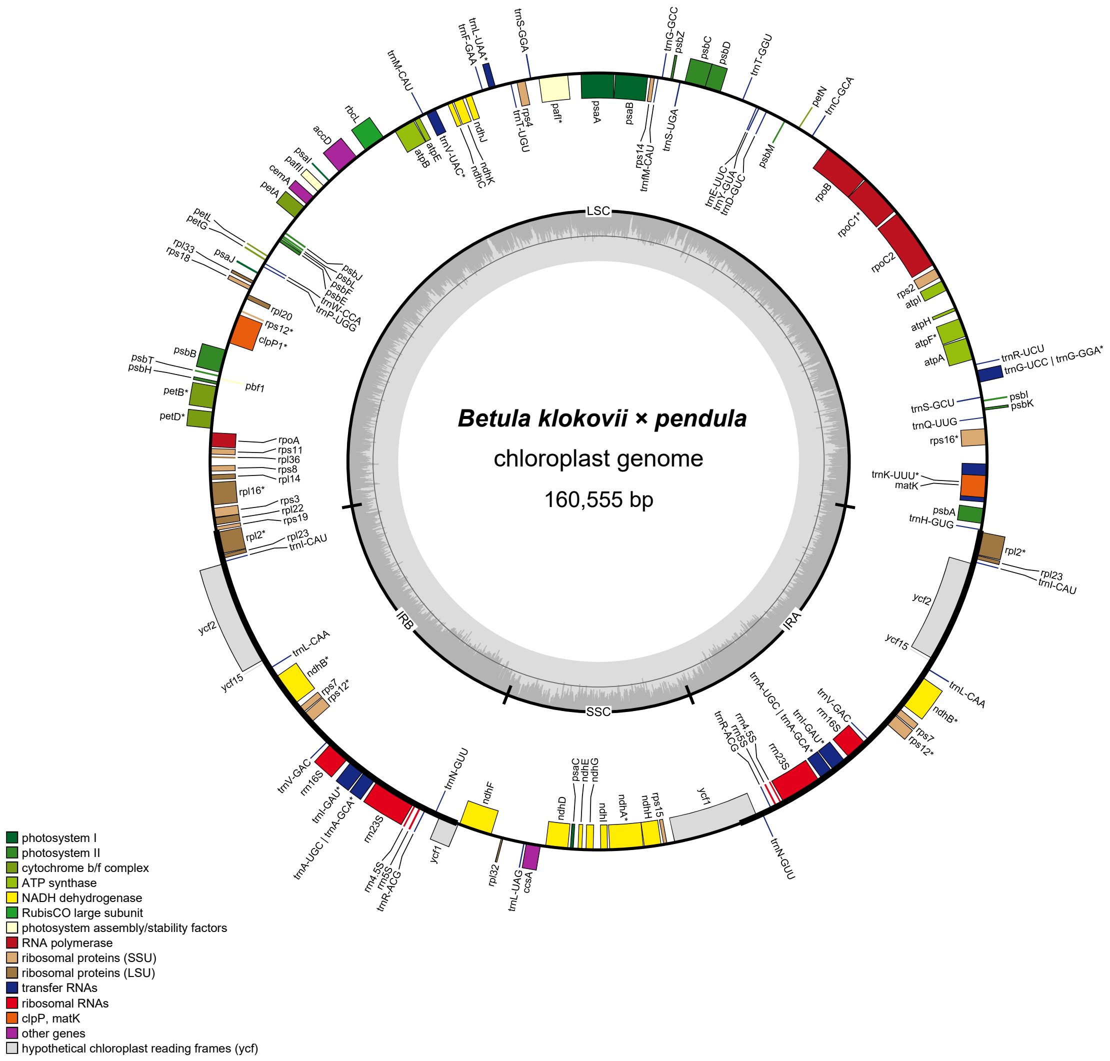


**Fig. S18** *Betula klokovii* × *Betula pendula* plastome assembly (sample 06 (PP551950); annotation – GeSeq; visualization – OGDraw v1.3.1).

The plot shows a circular organization of the plastome with two single-copy regions (large (LSC) and small (SSC), respectively) and two inverted repeats (IRA & IRB), and locations of the genes.

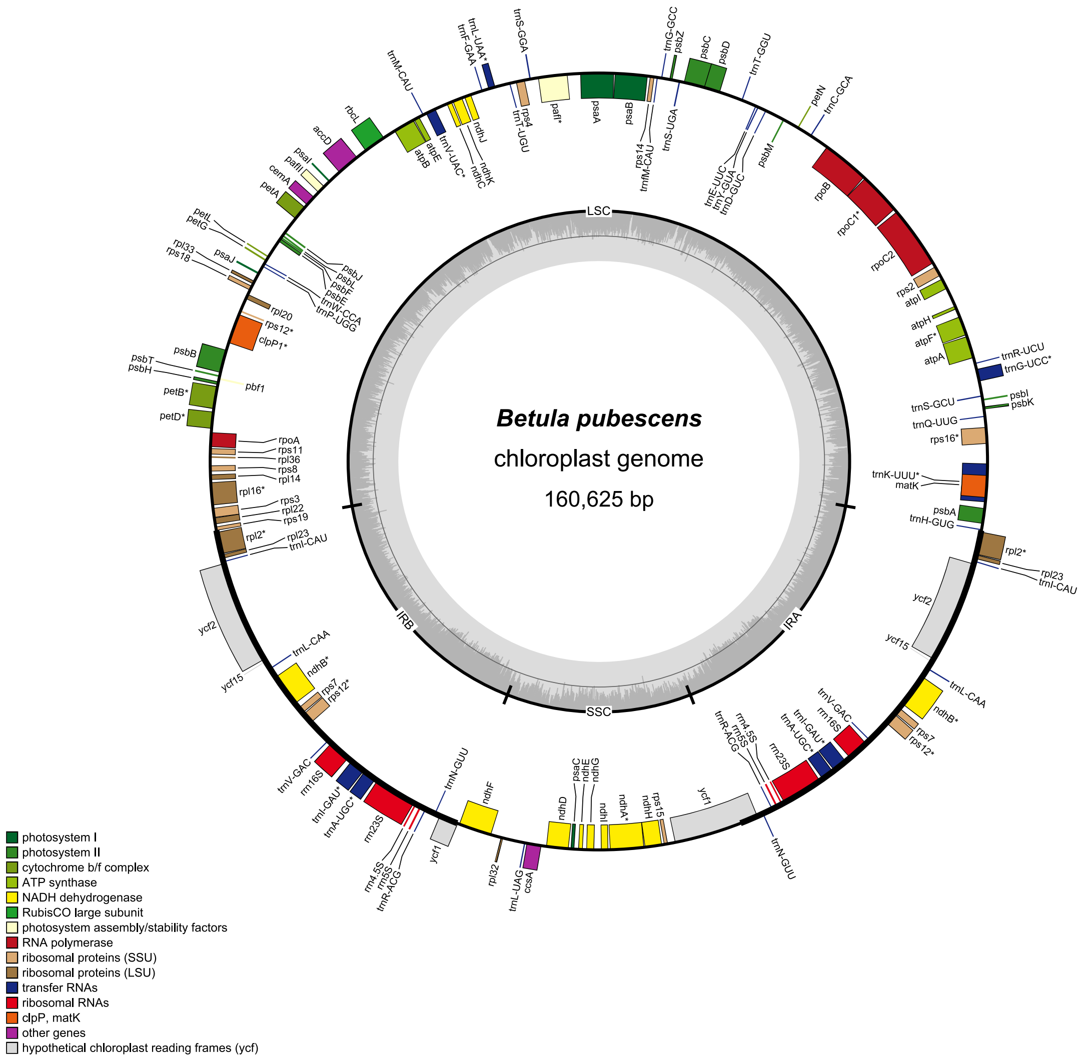


**Fig. S19** *Betula pubescens* plastome assembly (sample 07 (PP545465); annotation – GeSeq; visualization – OGDraw v1.3.1).

The plot shows a circular organization of the plastome with two single-copy regions (large (LSC) and small (SSC), respectively) and two inverted repeats (IRA & IRB), and locations of the genes.

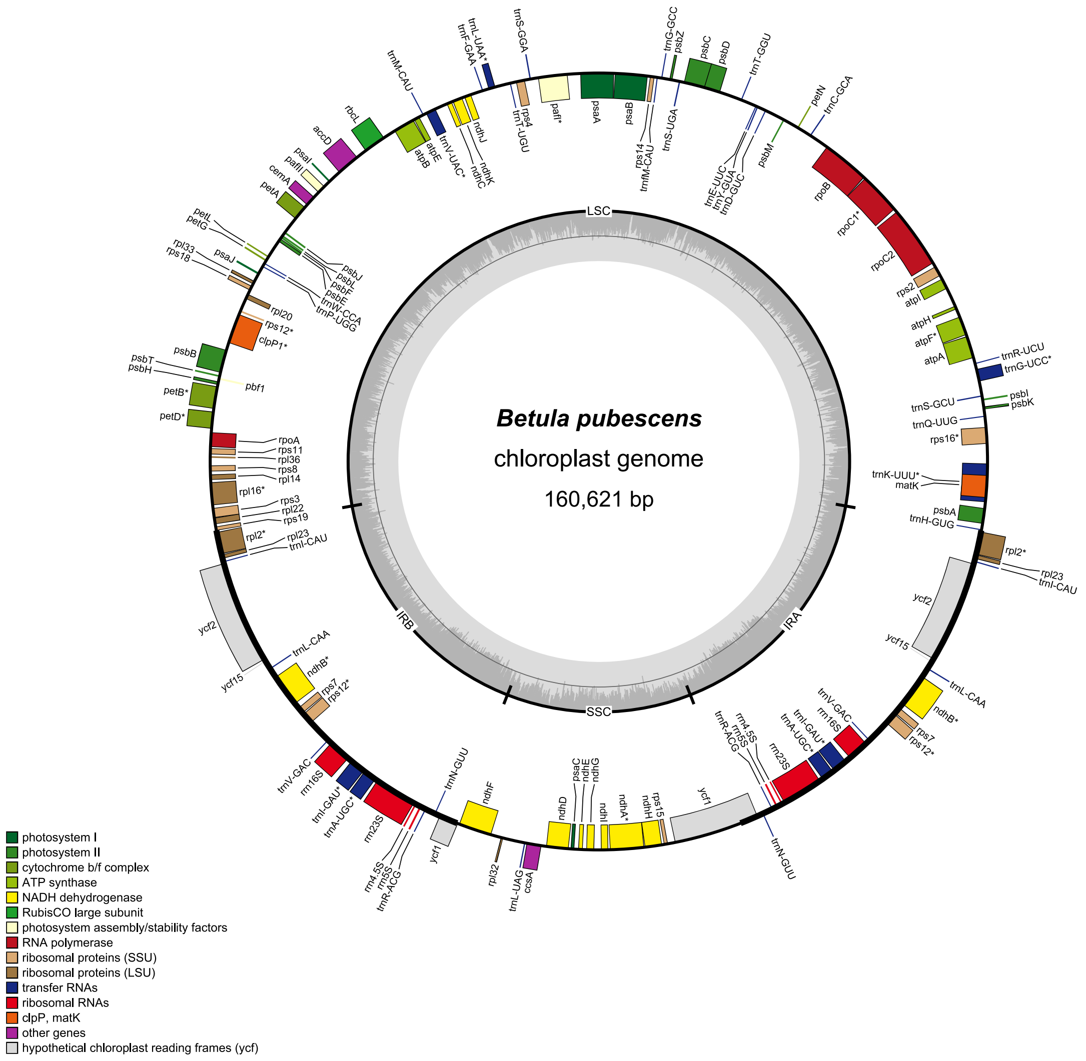


**Fig. S20** *Betula pubescens* plastome assembly (sample 08 (PP545466); annotation – GeSeq; visualization – OGDraw v1.3.1).

The plot shows a circular organization of the plastome with two single-copy regions (large (LSC) and small (SSC), respectively) and two inverted repeats (IRA & IRB), and locations of the genes.

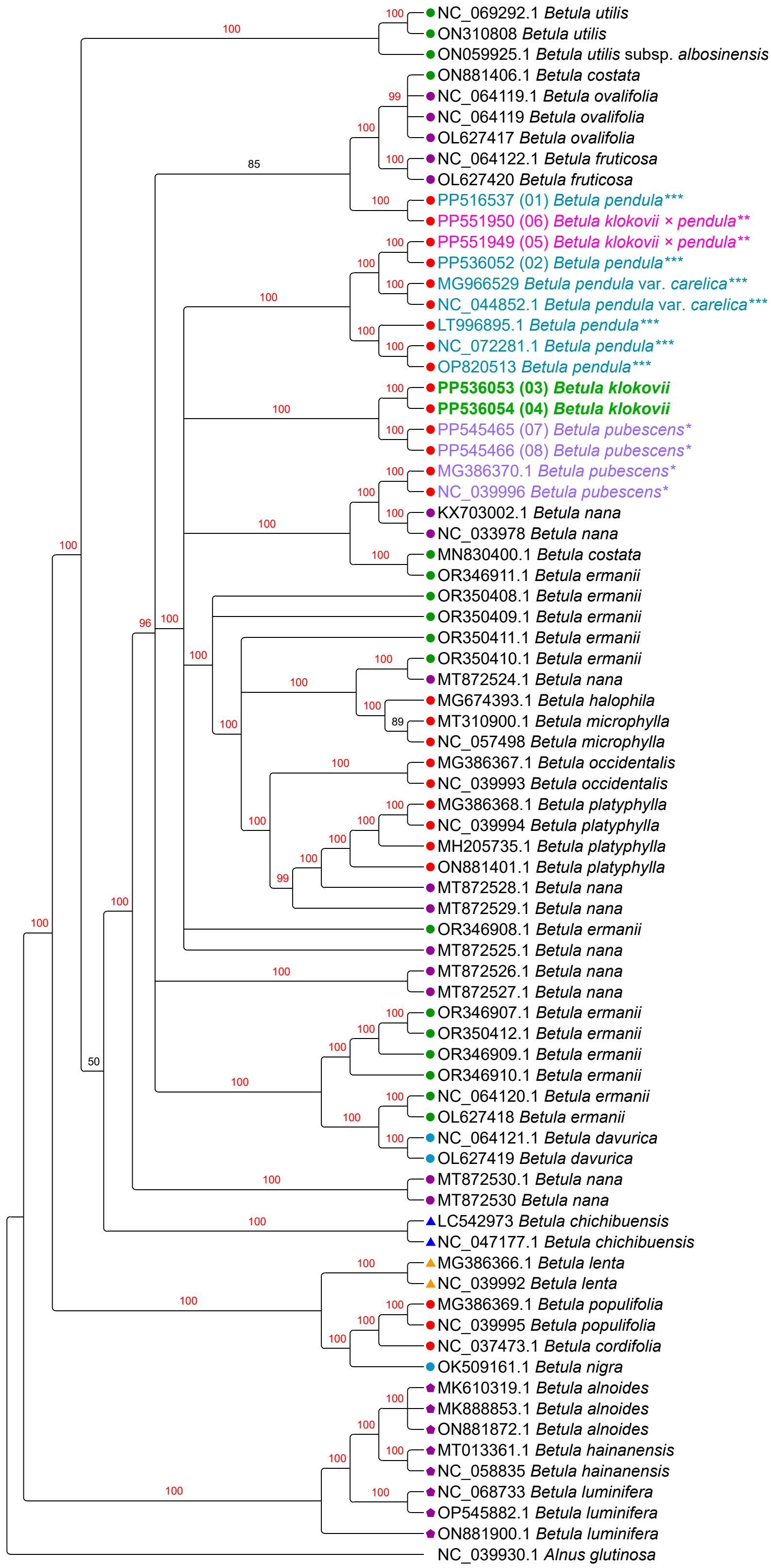


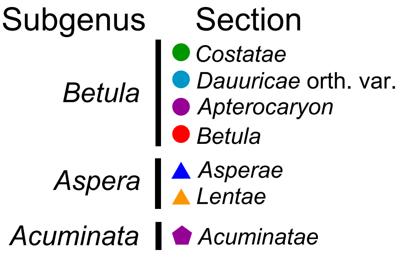


**Fig. S21** Plastome phylogeny based on all available plastome assemblies without partitioning, Bayesian Inference, MrBayes, GTR+I+G, 100 000 000 iterations with 10% burnin. Posterior probability values of supported clades are marked red. B. klokovii is colored in green and written in bold, presumed *B. pendula* × *B. klokovii* hybrids marked by double asterisk (**) and colored in magenta, *B. pubescens* is marked by asterisk (*) and colored in violet, *B. pendula* and its intraspecific taxa is marked by triple asterisk (***) and colored in cyan. Newly sequenced samples are indicated by numbers 01–08. Subgenera and sections are marked according to Wang & al., 2016.

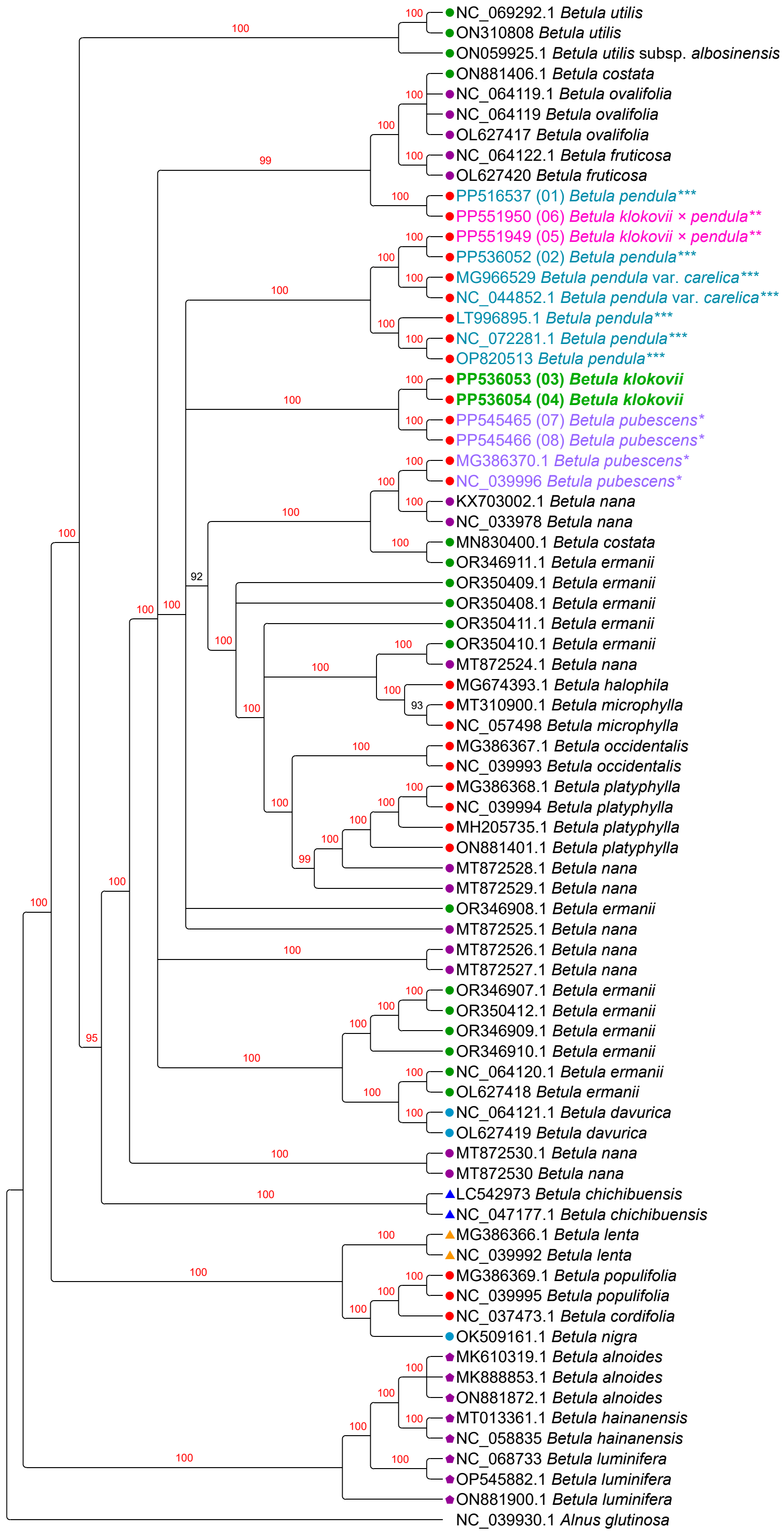


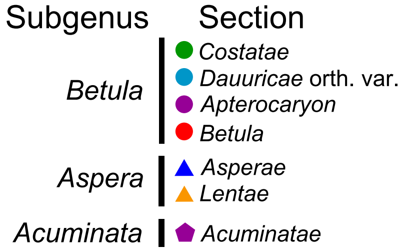


**Fig. S22** Plastome phylogeny based on all available plastome assemblies with partitioning, models assignment according to Akaike information criterion (AIC), Bayesian Inference, MrBayes, 100 000 000 iterations with 10% burnin. Posterior probability values of supported clades are marked red. *B. klokovii* is colored in green and written in bold, presumed *B. pendula* × *B. klokovii* hybrids marked by double asterisk (**) and colored in magenta, *B. pubescens* is marked by asterisk (*) and colored in violet, *B. pendula* and its intraspecific taxa is marked by triple asterisk (***) and colored in cyan. Newly sequenced samples are indicated by numbers 01–08. Subgenera and sections are marked according to Wang & al., 2016.

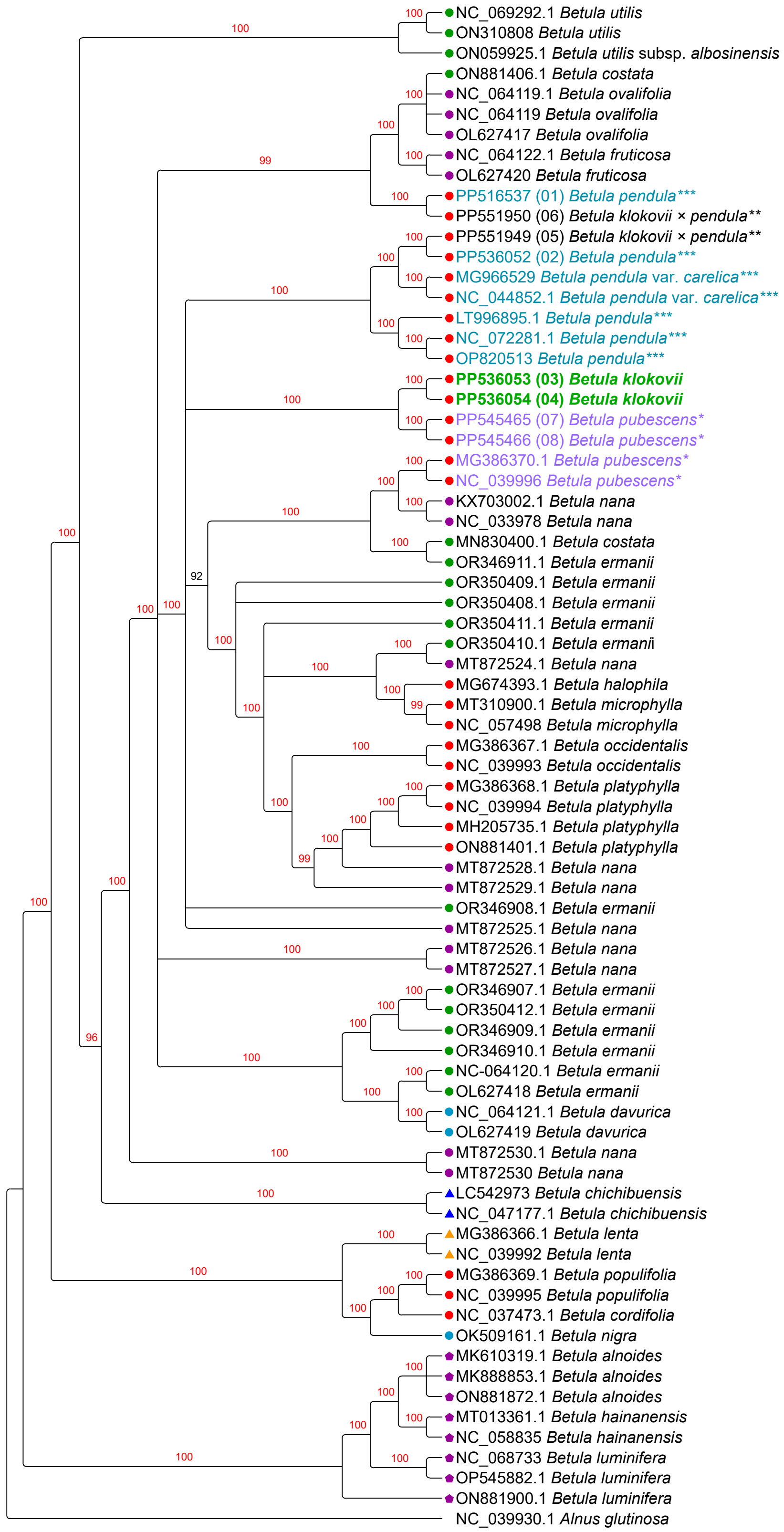


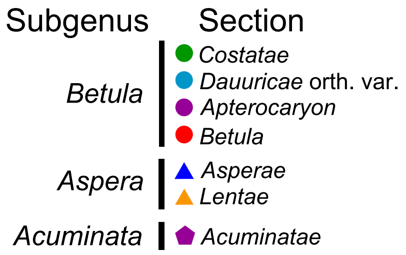


**Fig. S23** Plastome phylogeny based on all available plastome assemblies with partitioning, models assignment according to Akaike information criterion with small sample correction (AICc), Bayesian Inference, MrBayes, 100 000 000 iterations with 10% burnin. Posterior probability values of supported clades are marked red. *B. klokovii* is colored in green and written in bold, presumed *B. pendula* × *B. klokovii* hybrids marked by double asterisk (**) and colored in magenta, *B. pubescens* is marked by asterisk (*) and colored in violet, *B. pendula* and its intraspecific taxa is marked by triple asterisk (***) and colored in cyan. Newly sequenced samples are indicated by numbers 01–08. Subgenera and sections are marked according to Wang & al., 2016.

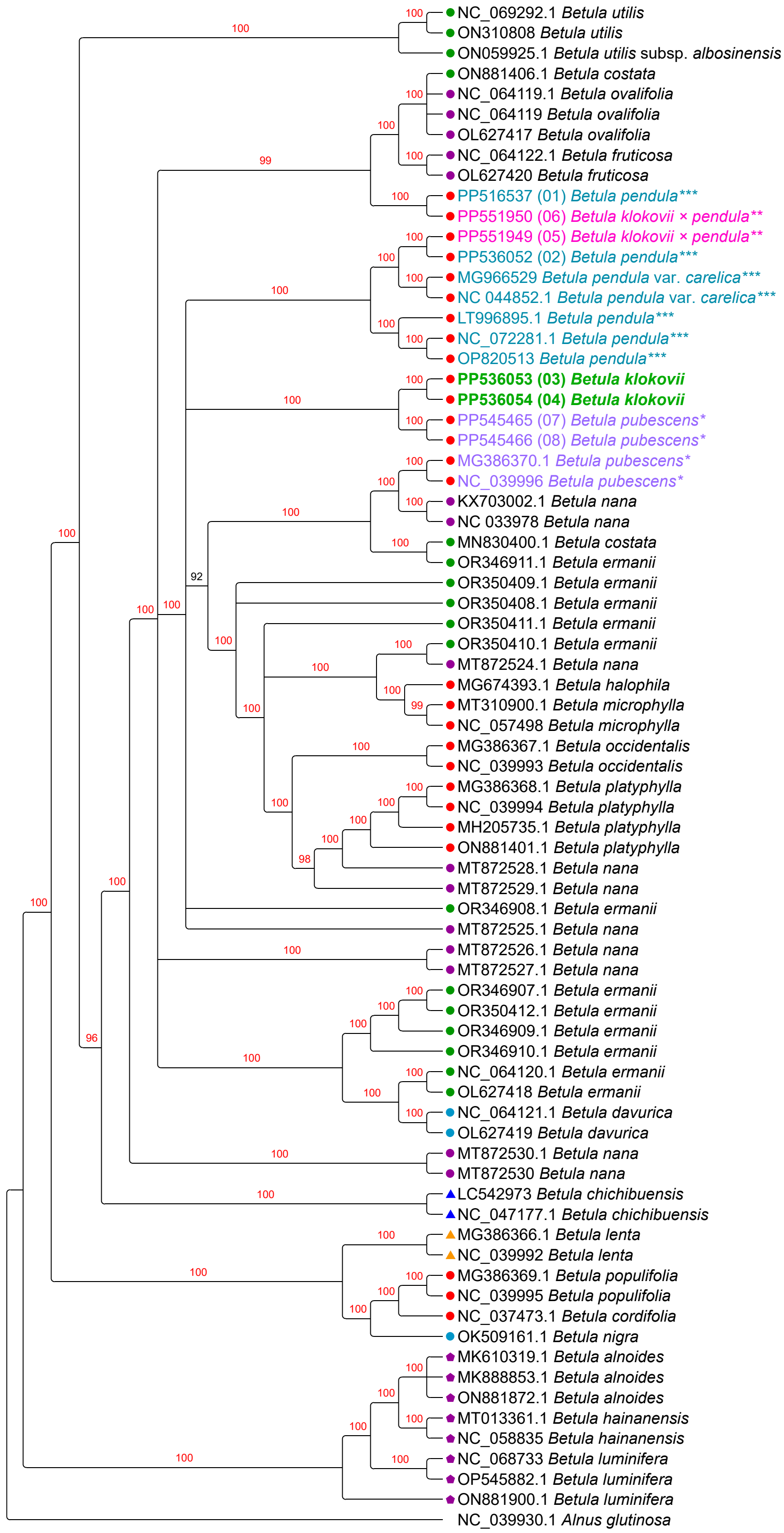


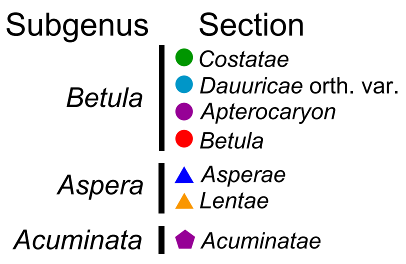


**Fig. S24** Plastome phylogeny based on all available plastome assemblies with partitioning, models assignment according to Bayesian Information Criterion (BIC), Bayesian Inference, MrBayes, 100 000 000 iterations with 10% burnin. Posterior probability values of supported clades are marked red. *B. klokovii* is colored in green and written in bold, presumed *B. pendula* × *B. klokovii* hybrids marked by double asterisk (**) and colored in magenta, *B. pubescens* is marked by asterisk (*) and colored in violet, *B. pendula* and its intraspecific taxa is marked by triple asterisk (***) and colored in cyan. Newly sequenced samples are indicated by numbers 01–08. Subgenera and sections are marked according to Wang & al., 2016.

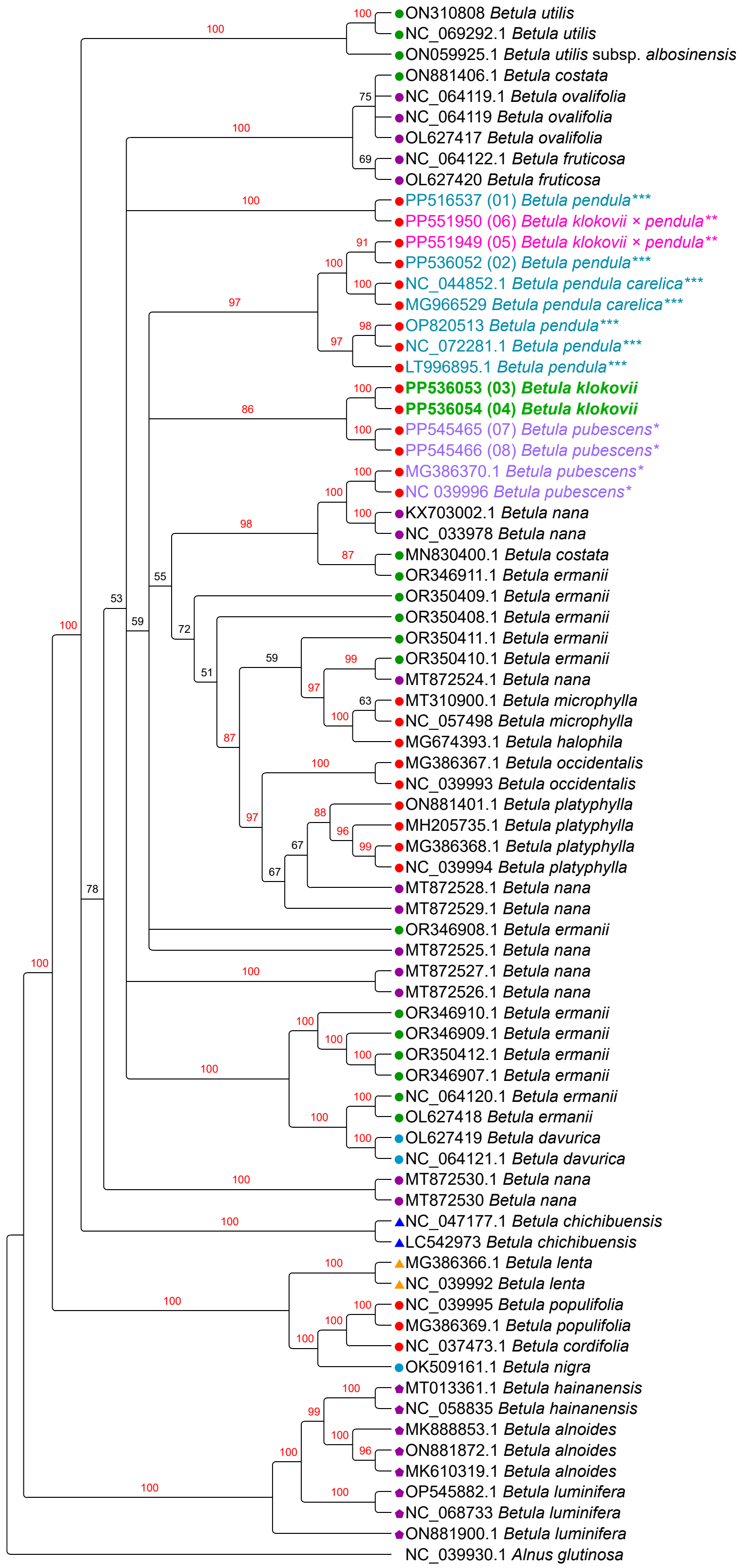

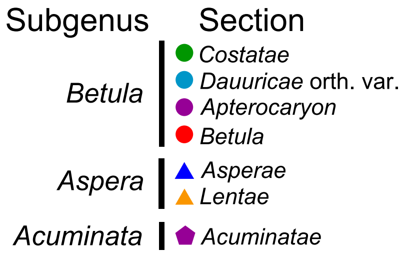


**Fig. S25** Plastome phylogeny based on all available plastome assemblies without partitioning, Maximum Likelihood, RAxML, TVM+I+G, 10 000 iterations bootstrap. Bootstrap values for highly supported clades are marked red. *B. klokovii* is colored in green and written in bold, presumed *B. pendula* × *B. klokovii* hybrids marked by double asterisk (**) and colored in magenta, *B. pubescens* is marked by asterisk (*) and colored in violet, *B. pendula* and its intraspecific taxa is marked by triple asterisk (***) and colored in cyan. Newly sequenced samples are indicated by numbers 01–08. Subgenera and sections are marked according to Wang & al., 2016.

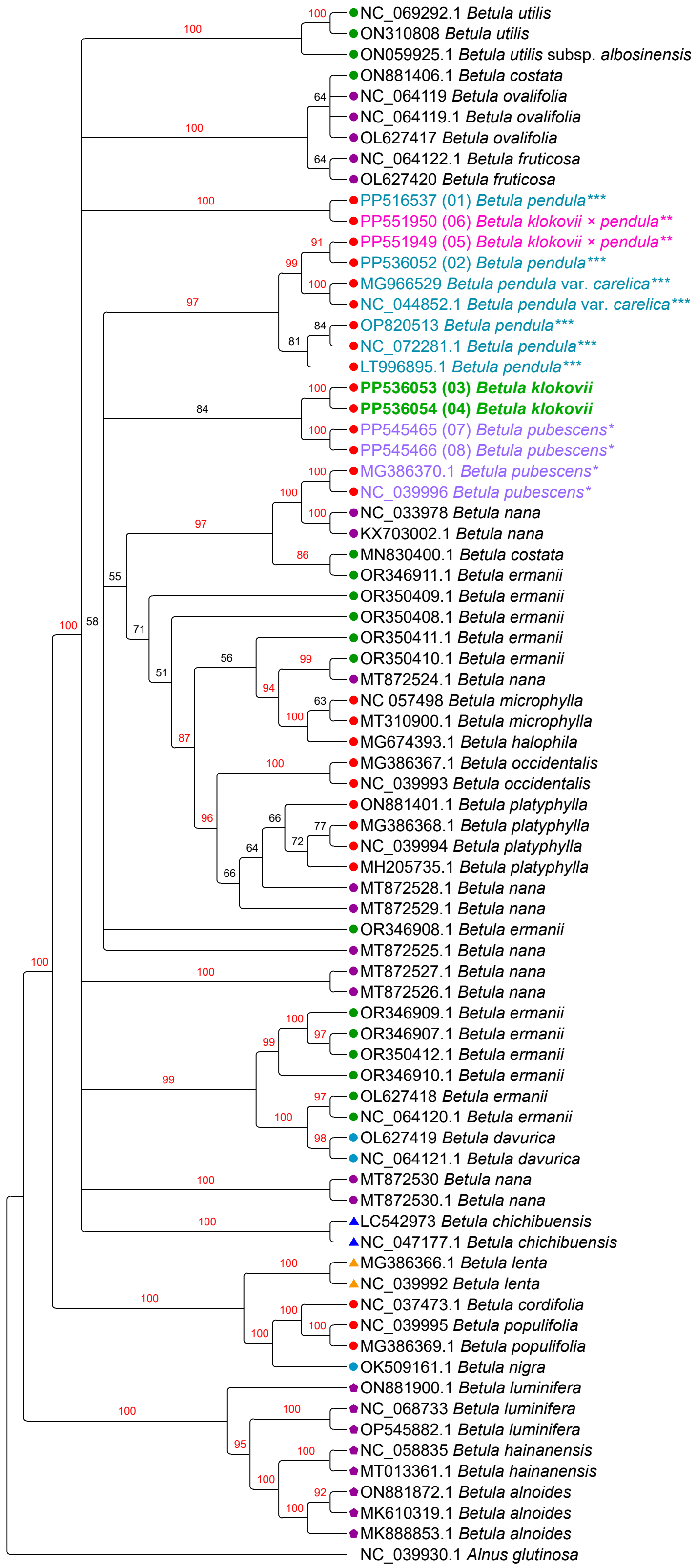


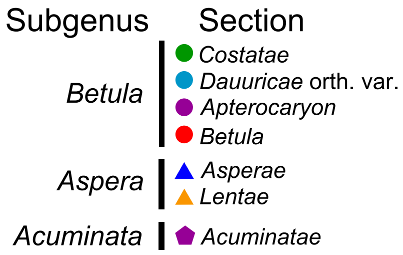


**Fig. S26** Plastome phylogeny based on all available plastome assemblies with partitioning, models assignment according to Akaike information criterion (AIC), Maximum Likelihood, RAxML, 10 000 iterations bootstrap. Bootstrap values for highly supported clades are marked red. *B. klokovii* is colored in green and written in bold, presumed *B. pendula* × *B. klokovii* hybrids marked by double asterisk (**) and colored in magenta, *B. pubescens* is marked by asterisk (*) and colored in violet, *B. pendula* and its intraspecific taxa is marked by triple asterisk (***) and colored in cyan. Newly sequenced samples are indicated by numbers 01–08. Subgenera and sections are marked according to Wang & al., 2016.

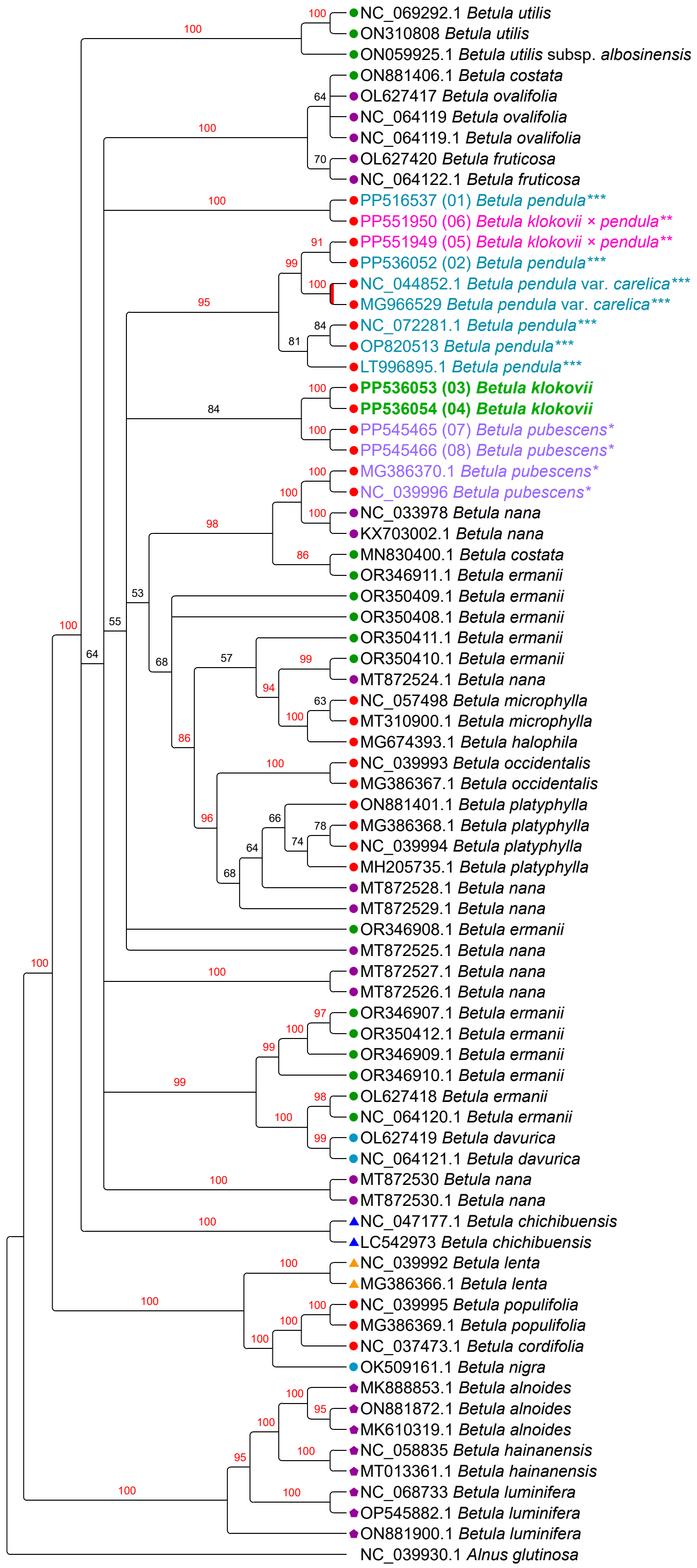


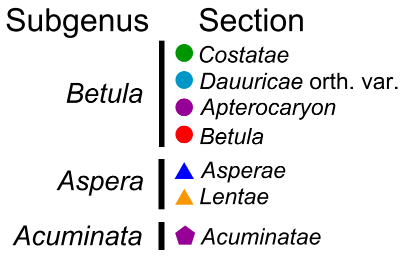


**Fig. S27** Plastome phylogeny reconstruction based on all available plastome assemblies with partitioning, models assignment according to Akaike information criterion with small sample correction (AICc), Maximum Likelihood, RAxML, 10 000 iterations bootstrap. Bootstrap values for highly supported clades are marked red. *B. klokovii* is colored in green and written in bold, presumed *B. pendula* × *B. klokovii* hybrids marked by double asterisk (**) and colored in magenta, *B. pubescens* is marked by asterisk (*) and colored in violet, *B. pendula* and its intraspecific taxa is marked by triple asterisk (***) and colored in cyan. Newly sequenced samples are indicated by numbers 01–08. Subgenera and sections are marked according to Wang & al., 2016.

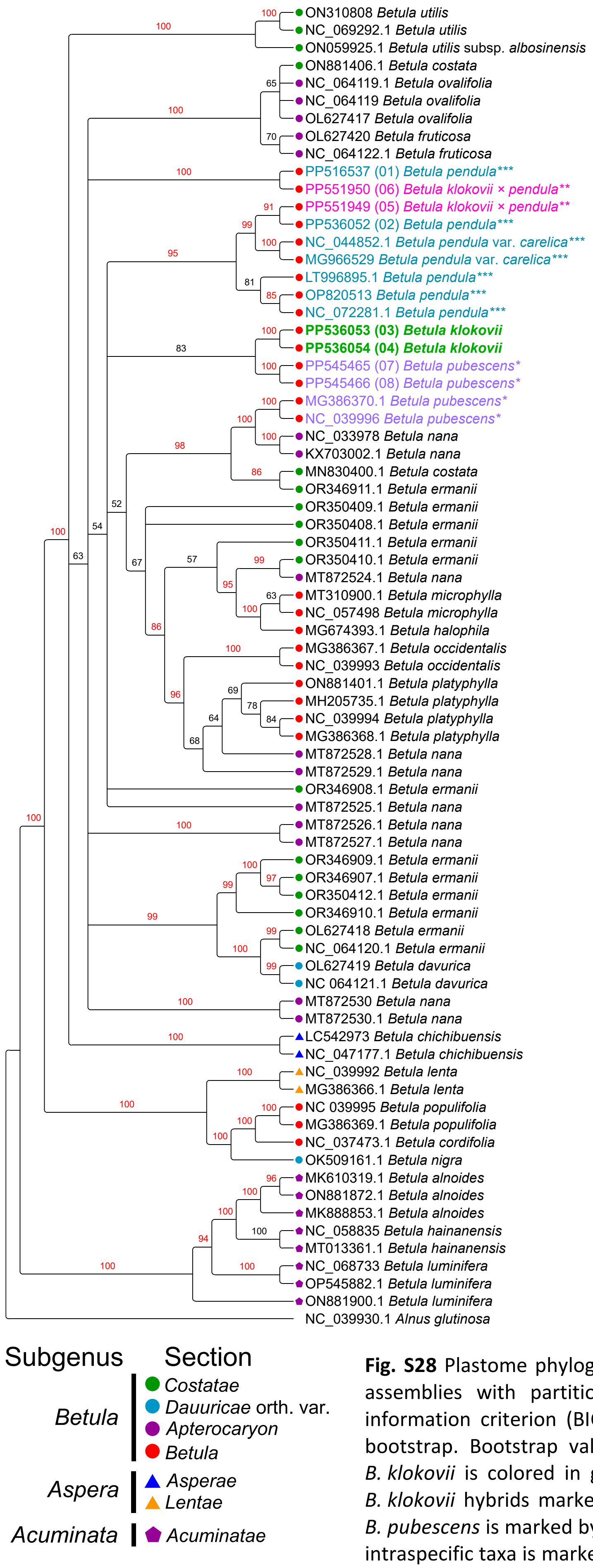


**Fig. S28** Plastome phylogeny reconstruction based on all available plastome assemblies with partitioning, models assignment according to Bayesian information criterion (BIC), Maximum Likelihood, RAxML, 10 000 iterations bootstrap. Bootstrap values for highly supported clades are marked red. *B. klokovii* is colored in green and written in bold, presumed *B. pendula* × *B. klokovii* hybrids marked by double asterisk (**) and colored in magenta, *B. pubescens* is marked by asterisk (*) and colored in violet, *B. pendula* and its intraspecific taxa is marked by triple asterisk (***) and colored in cyan. Newly sequenced samples are indicated by numbers 01–08. Subgenera and sections are marked according to Wang & al., 2016.

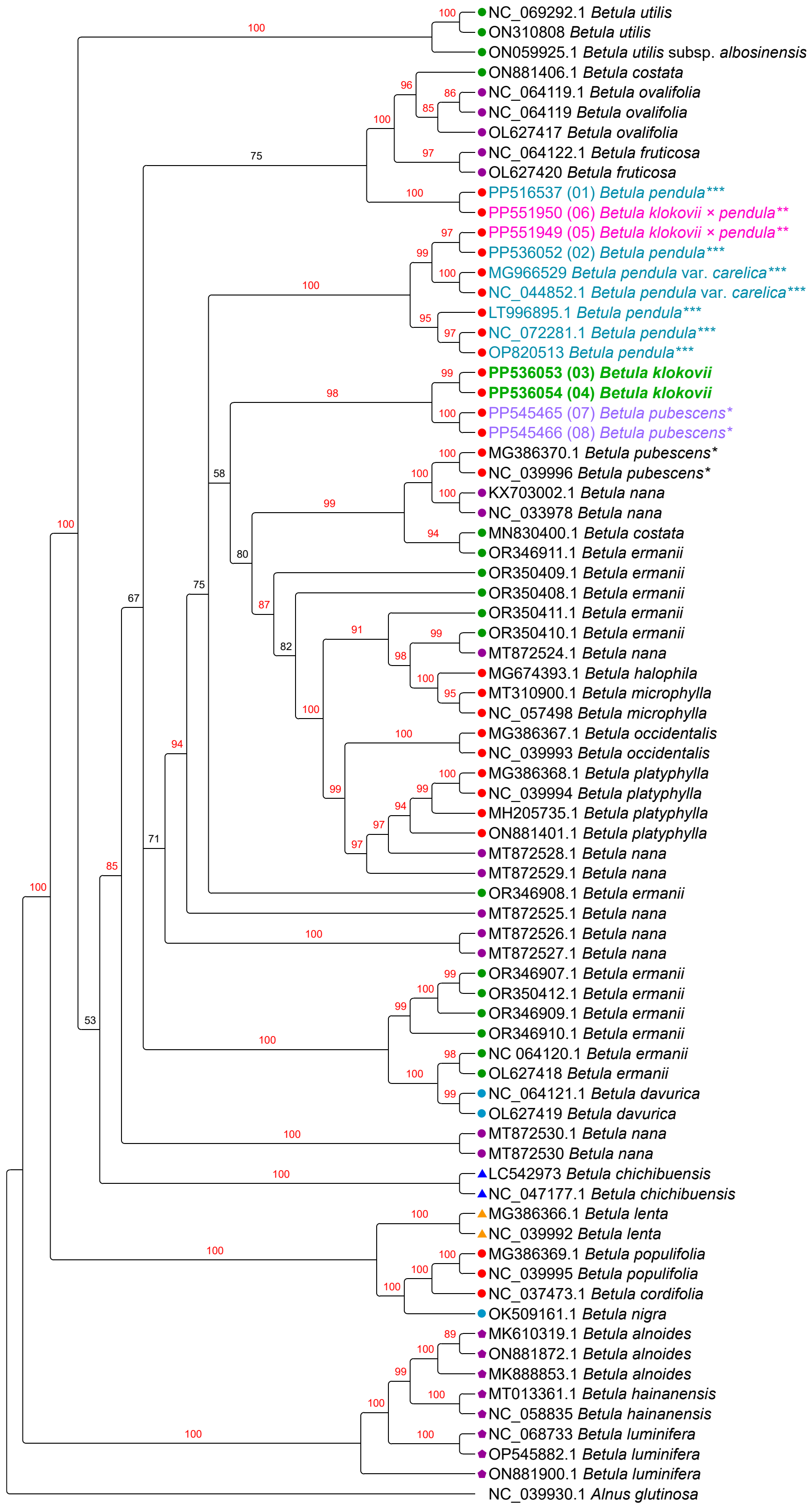


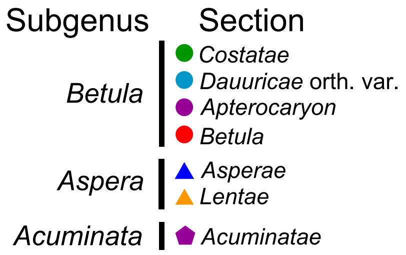


**Fig. S29** Plastome phylogeny based on all available plastome assemblies without partitioning, Maximum Likelihood, IQ-TREE1, TVM+I+G, 10 000 iterations ultrafast bootstrap. Bootstrap values for highly supported clades are marked red. *B. klokovii* is colored in green and written in bold, presumed *B. pendula* × *B. klokovii* hybrids marked by double asterisk (**) and colored in magenta, *B. pubescens* is marked by asterisk (*) and colored in violet, *B. pendula* and its intraspecific taxa is marked by triple asterisk (***) and colored in cyan. Newly sequenced samples are indicated by numbers 01–08. Subgenera and sections are marked according to Wang & al., 2016.

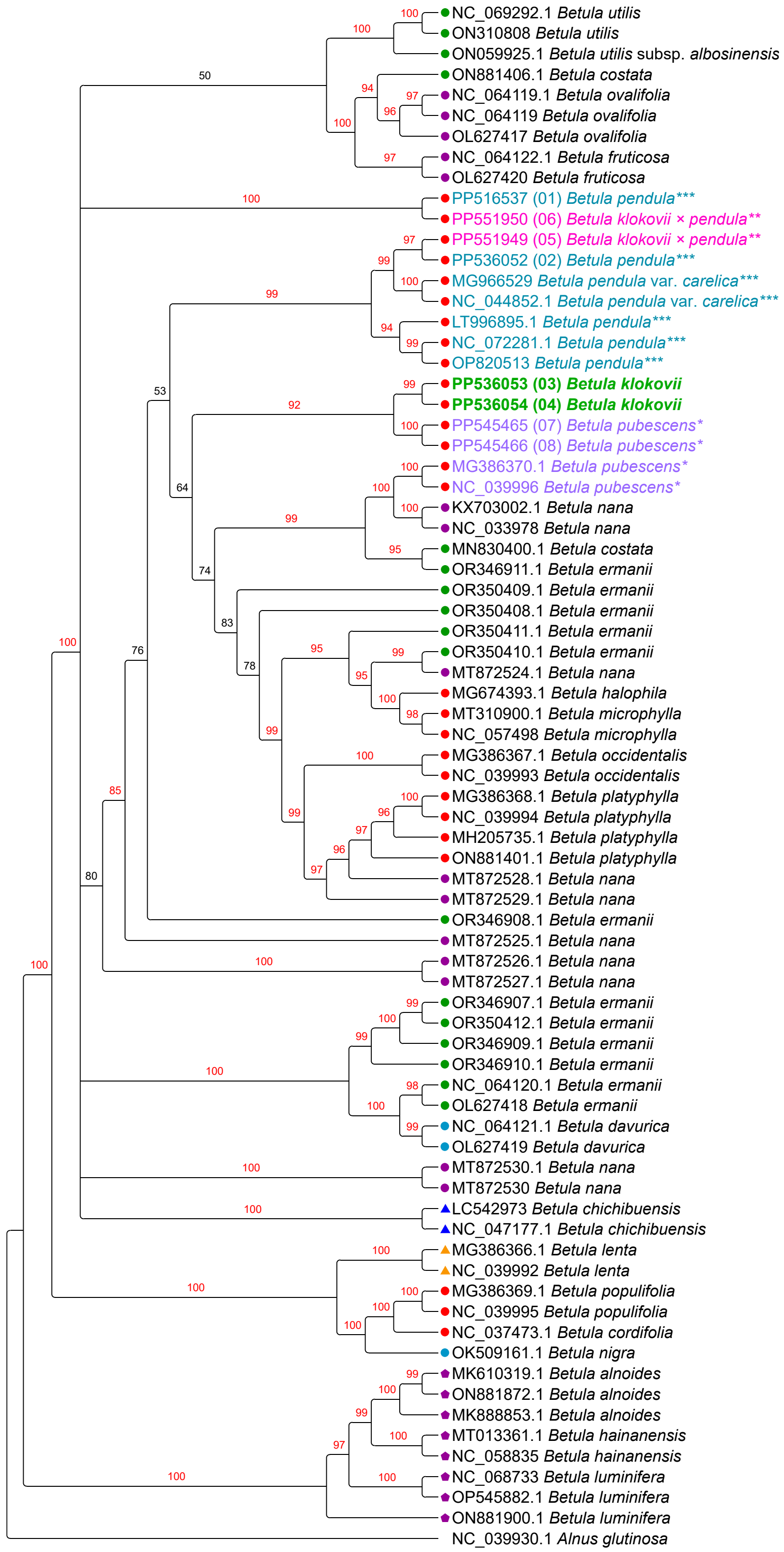

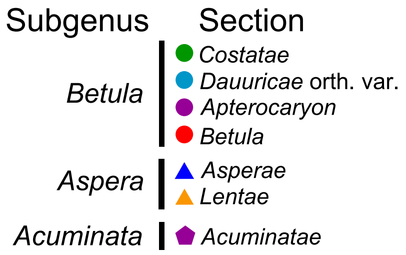


**Fig. S30** Plastome phylogeny based on all available plastome assemblies with partitioning, models assignment according to Akaike information criterion (AIC) Maximum Likelihood, IQ-TREE1, 10 000 iterations ultrafast bootstrap. Bootstrap values for highly supported clades are marked red. *B. klokovii* is colored in green and written in bold, presumed *B. pendula* × *B. klokovii* hybrids marked by double asterisk (**) and colored in magenta, *B. pubescens* is marked by asterisk (*) and colored in violet, *B. pendula* and its intraspecific taxa is marked by triple asterisk (***) and colored in cyan. Newly sequenced samples are indicated by numbers 01–08. Subgenera and sections are marked according to Wang & al., 2016.

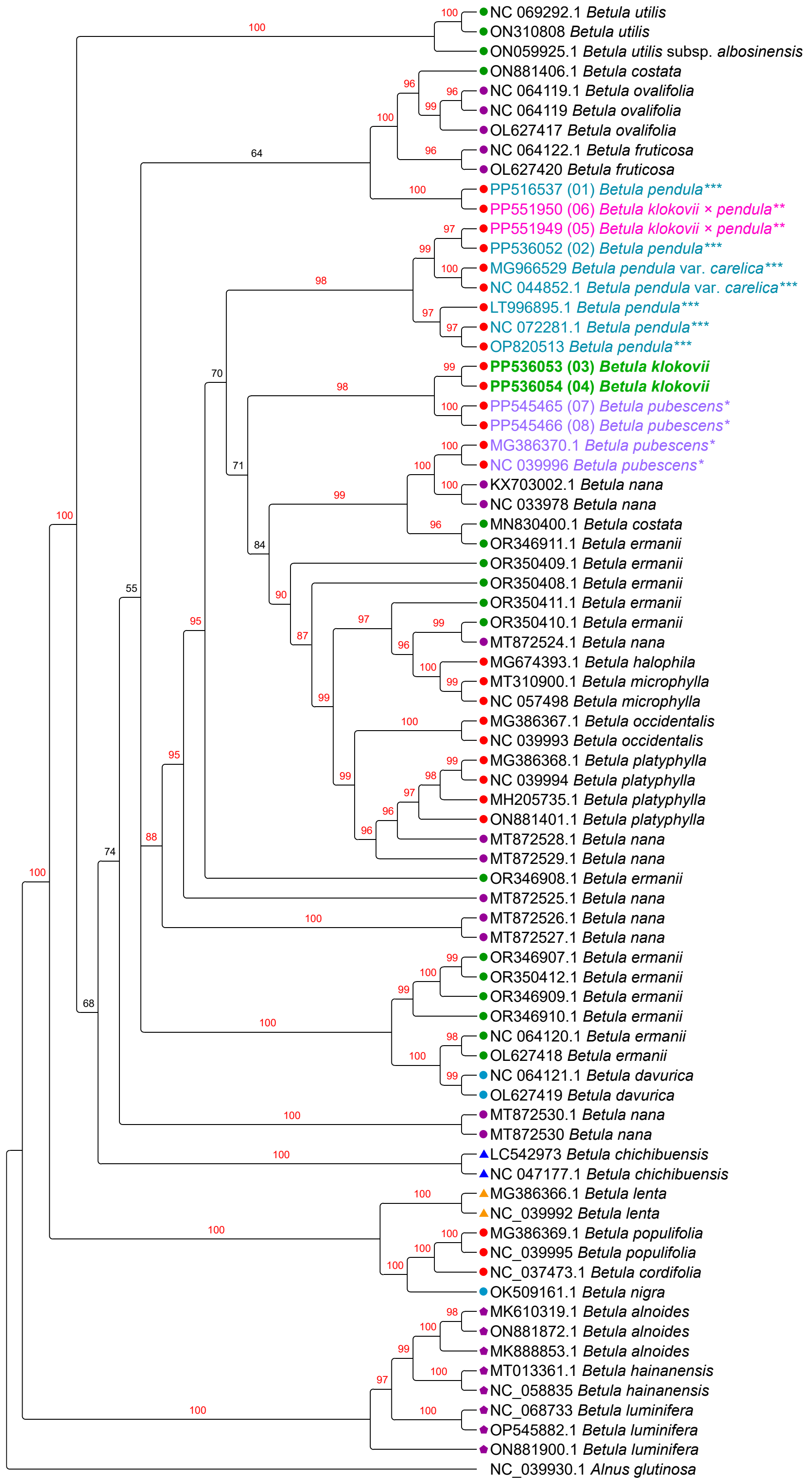


| Subgenus | Section |
|---|---|
| *Betula* | ● *Costatae* |
| | ● *Dauuricae* orth. var. |
| | ● *Apterocaryon* |
| | ● *Betula* |
| *Aspera* | ▲ *Asperae* |
| | ▲ *Lentae* |
| *Acuminata* | ⬟ *Acuminatae* |

**Fig. S31** Plastome phylogeny based on all available plastome assemblies with partitioning, models assignment according to Akaike information criterion with small sample correction (AICc), Maximum Likelihood, IQ-TREE1, 10 000 iterations ultrafast bootstrap. Bootstrap values for highly supported clades are marked red. *B. klokovii* is colored in green and written in bold, presumed *B. pendula* × *B. klokovii* hybrids marked by double asterisk (**) and colored in magenta, *B. pubescens* is marked by asterisk (*) and colored in violet, *B. pendula* and its intraspecific taxa is marked by triple asterisk (***) and colored in cyan. Newly sequenced samples are indicated by numbers 01–08. Subgenera and sections are marked according to Wang & al., 2016.

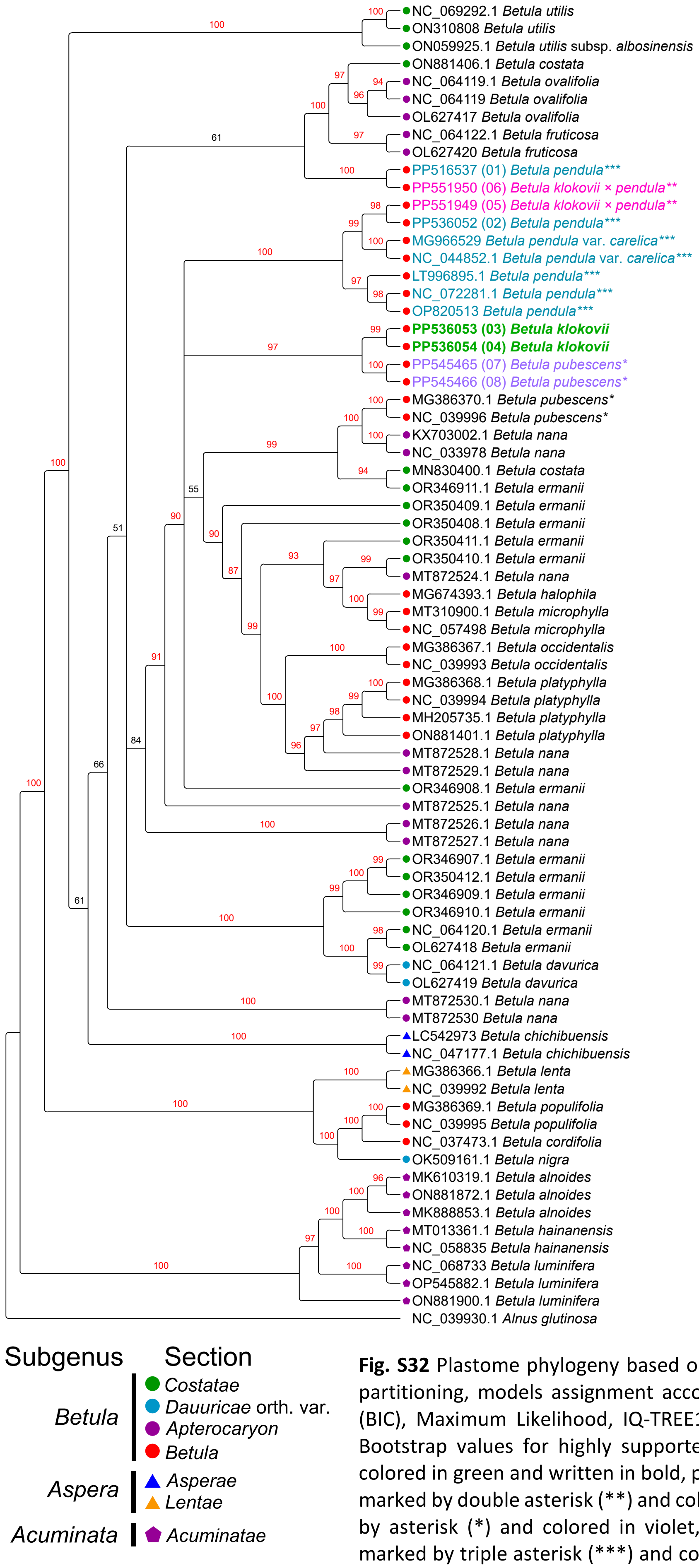


**Fig. S32** Plastome phylogeny based on all available plastome assemblies with partitioning, models assignment according to Bayesian information criterion (BIC), Maximum Likelihood, IQ-TREE1, 10 000 iterations ultrafast bootstrap. Bootstrap values for highly supported clades are marked red. *B. klokovii* is colored in green and written in bold, presumed *B. pendula* × *B. klokovii* hybrids marked by double asterisk (**) and colored in magenta, *B. pubescens* is marked by asterisk (*) and colored in violet, *B. pendula* and its intraspecific taxa is marked by triple asterisk (***) and colored in cyan. Newly sequenced samples are indicated by numbers 01–08. Subgenera and sections are marked according to Wang & al., 2016.

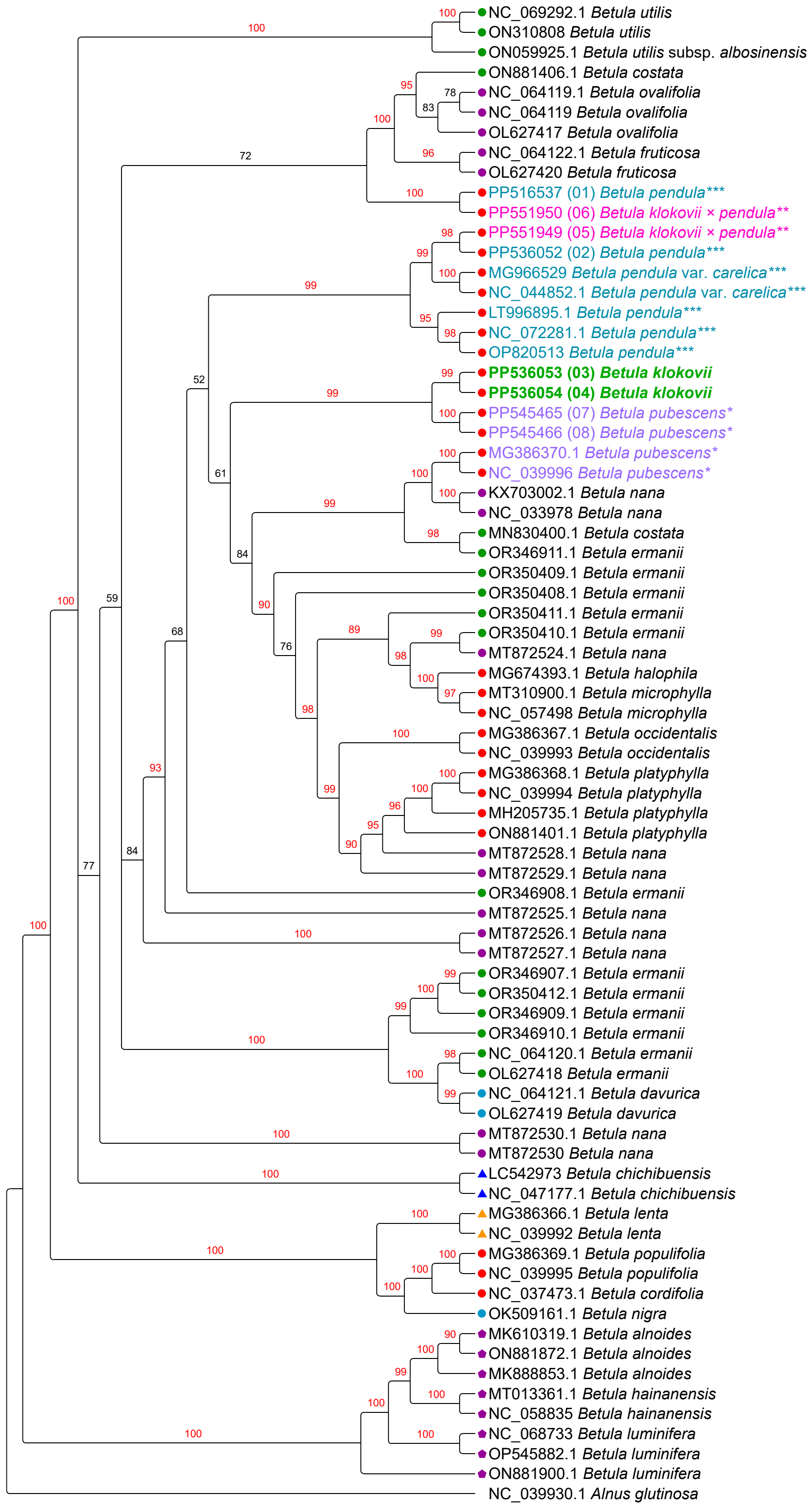


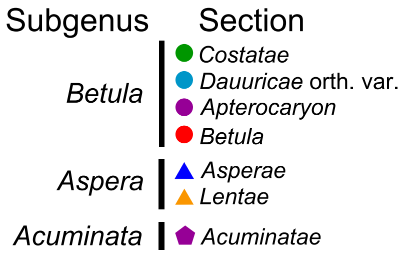


**Fig. S33** Plastome phylogeny based on all available plastome assemblies without partitioning, Maximum Likelihood, IQ-TREE2, TVM+I+G, 10 000 iterations ultrafast bootstrap. Bootstrap values for highly supported clades are marked red. *B. klokovii* is colored in green and written in bold, presumed *B. pendula* × *B. klokovii* hybrids marked by double asterisk (**) and colored in magenta, *B. pubescens* is marked by asterisk (*) and colored in violet, *B. pendula* and its intraspecific taxa is marked by triple asterisk (***) and colored in cyan. Newly sequenced samples are indicated by numbers 01–08. Subgenera and sections are marked according to Wang & al., 2016.

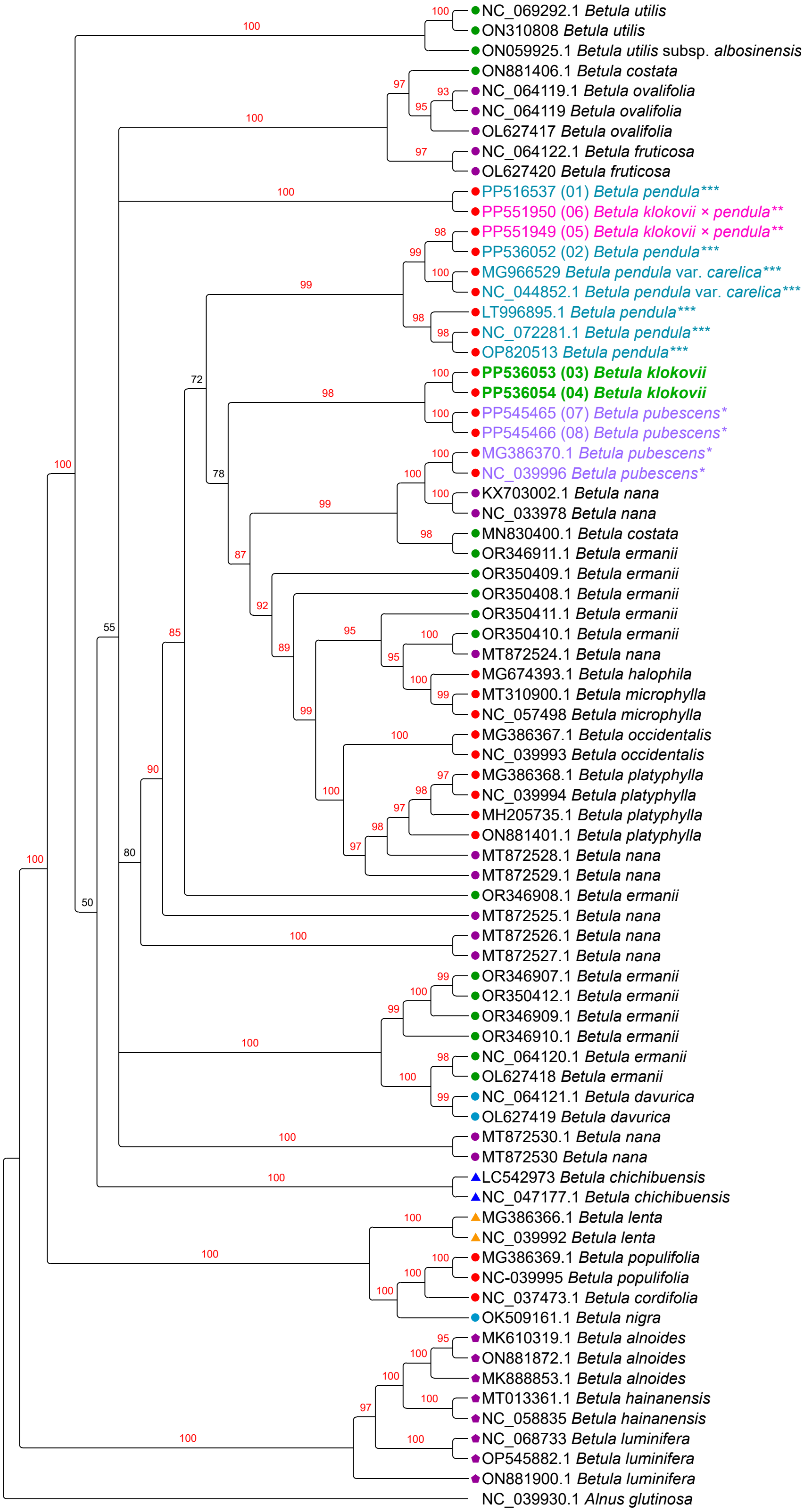


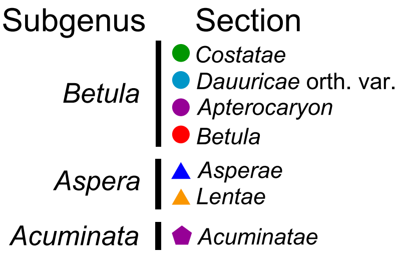


**Fig. S34** Plastome phylogeny based on all available plastome assemblies with partitioning, models assignment according to Akaike information criterion (AIC), Maximum Likelihood, IQ-TREE2, 10 000 iterations ultrafast bootstrap. Bootstrap values for highly supported clades are marked red. *B. klokovii* is colored in green and written in bold, presumed *B. pendula* × *B. klokovii* hybrids marked by double asterisk (**) and colored in magenta, *B. pubescens* is marked by asterisk (*) and colored in violet, *B. pendula* and its intraspecific taxa is marked by triple asterisk (***) and colored in cyan. Newly sequenced samples are indicated by numbers 01–08. Subgenera and sections are marked according to Wang & al., 2016.

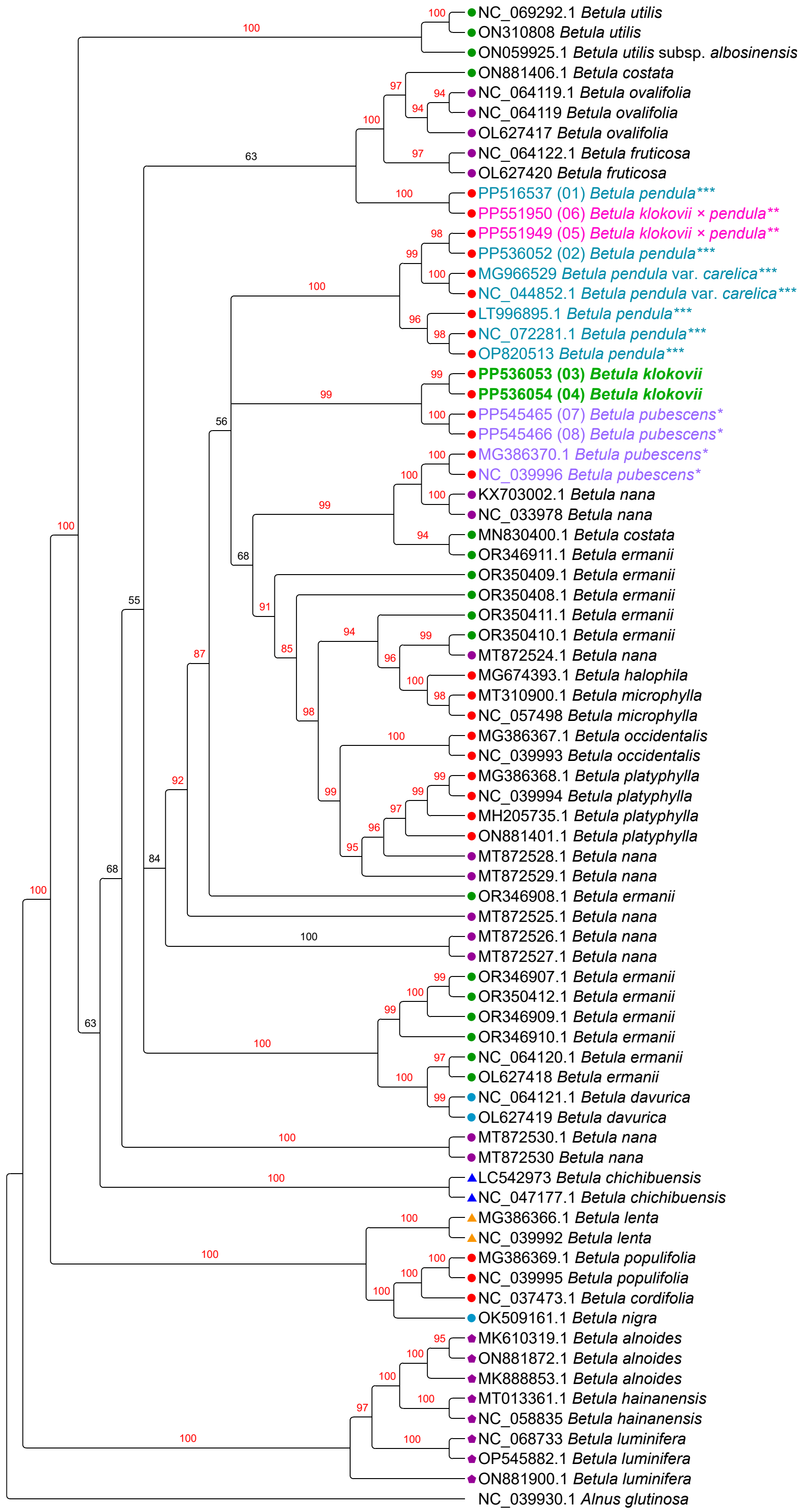


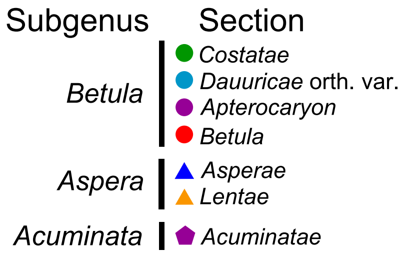


**Fig. S35** Plastome phylogeny based on all available plastome assemblies with partitioning, models assignment according to Akaike information criterion with small sample correction (AICc), Maximum Likelihood, IQ-TREE2, 10 000 iterations ultrafast bootstrap. Bootstrap values for highly supported clades are marked red. *B. klokovii* is colored in green and written in bold, presumed *B. pendula* × *B. klokovii* hybrids marked by double asterisk (**) and colored in magenta, *B. pubescens* is marked by asterisk (*) and colored in violet, *B. pendula* and its intraspecific taxa is marked by triple asterisk (***) and colored in cyan. Newly sequenced samples are indicated by numbers 01–08. Subgenera and sections are marked according to Wang & al., 2016.

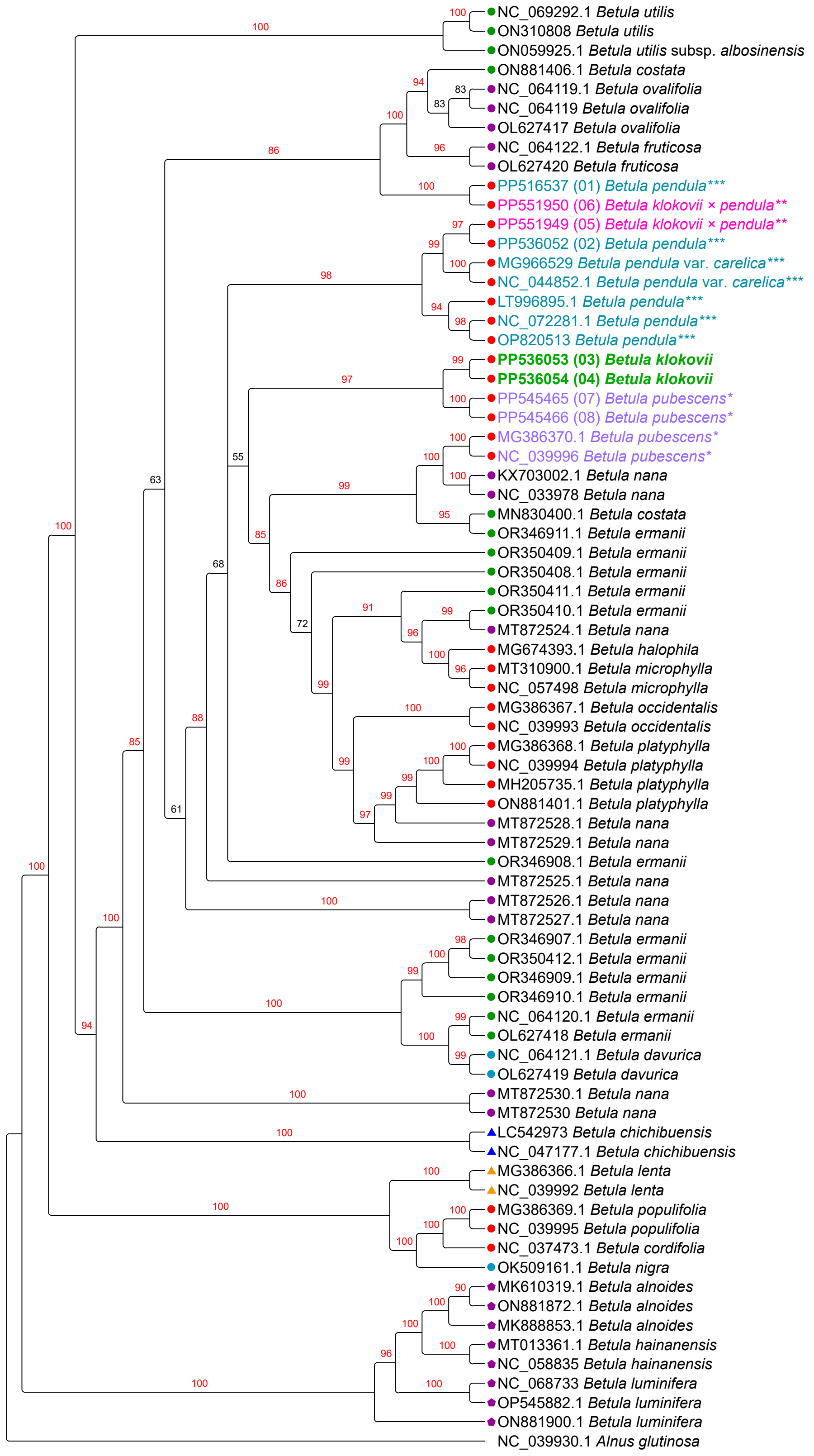


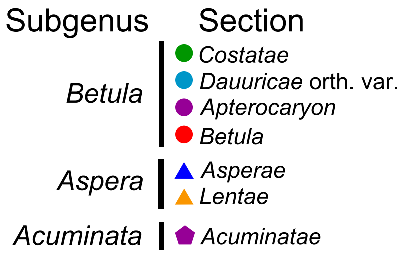


**Fig. S36** Plastome phylogeny based on all available plastome assemblies with partitioning, models assignment according to Bayesian information criterion (BIC), Maximum Likelihood, IQ-TREE2, 10 000 iterations ultrafast bootstrap. Bootstrap values for highly supported clades are marked red. *B. klokovii* is colored in green and written in bold, presumed *B. pendula* × *B. klokovii* hybrids marked by double asterisk (**) and colored in magenta, *B. pubescens* is marked by asterisk (*) and colored in violet, *B. pendula* and its intraspecific taxa is marked by triple asterisk (***) and colored in cyan. Newly sequenced samples are indicated by numbers 01–08. Subgenera and sections are marked according to Wang & al., 2016.

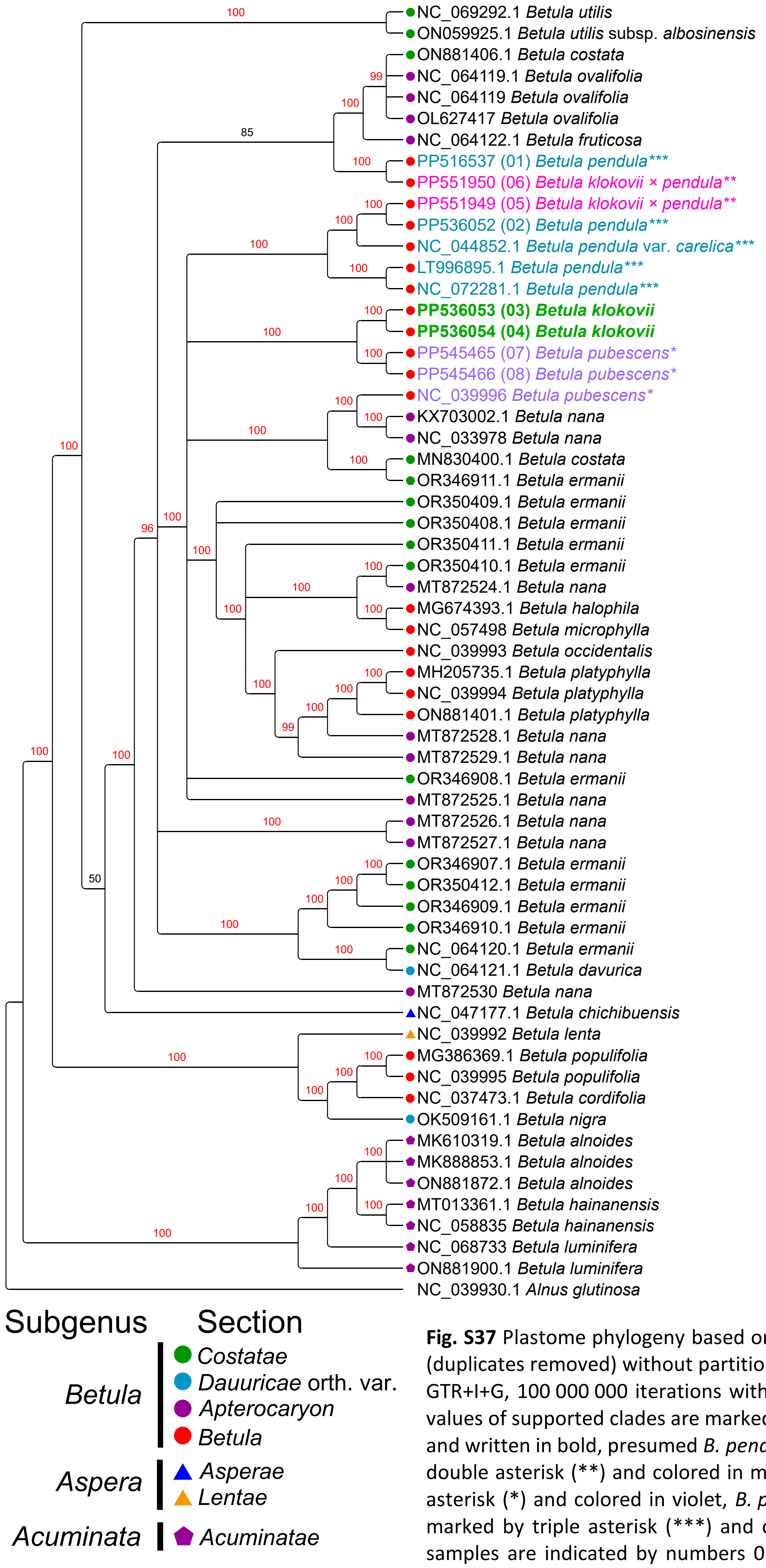


**Fig. S37** Plastome phylogeny based on unique plastome assemblies only (duplicates removed) without partitioning, Bayesian Inference, MrBayes, GTR+I+G, 100 000 000 iterations with 10% burnin. Posterior probability values of supported clades are marked red. *B. klokovii* is colored in green and written in bold, presumed *B. pendula* × *B. klokovii* hybrids marked by double asterisk (**) and colored in magenta, *B. pubescens* is marked by asterisk (*) and colored in violet, *B. pendula* and its intraspecific taxa is marked by triple asterisk (***) and colored in cyan. Newly sequenced samples are indicated by numbers 01–08. Subgenera and sections are marked according to Wang & al., 2016.

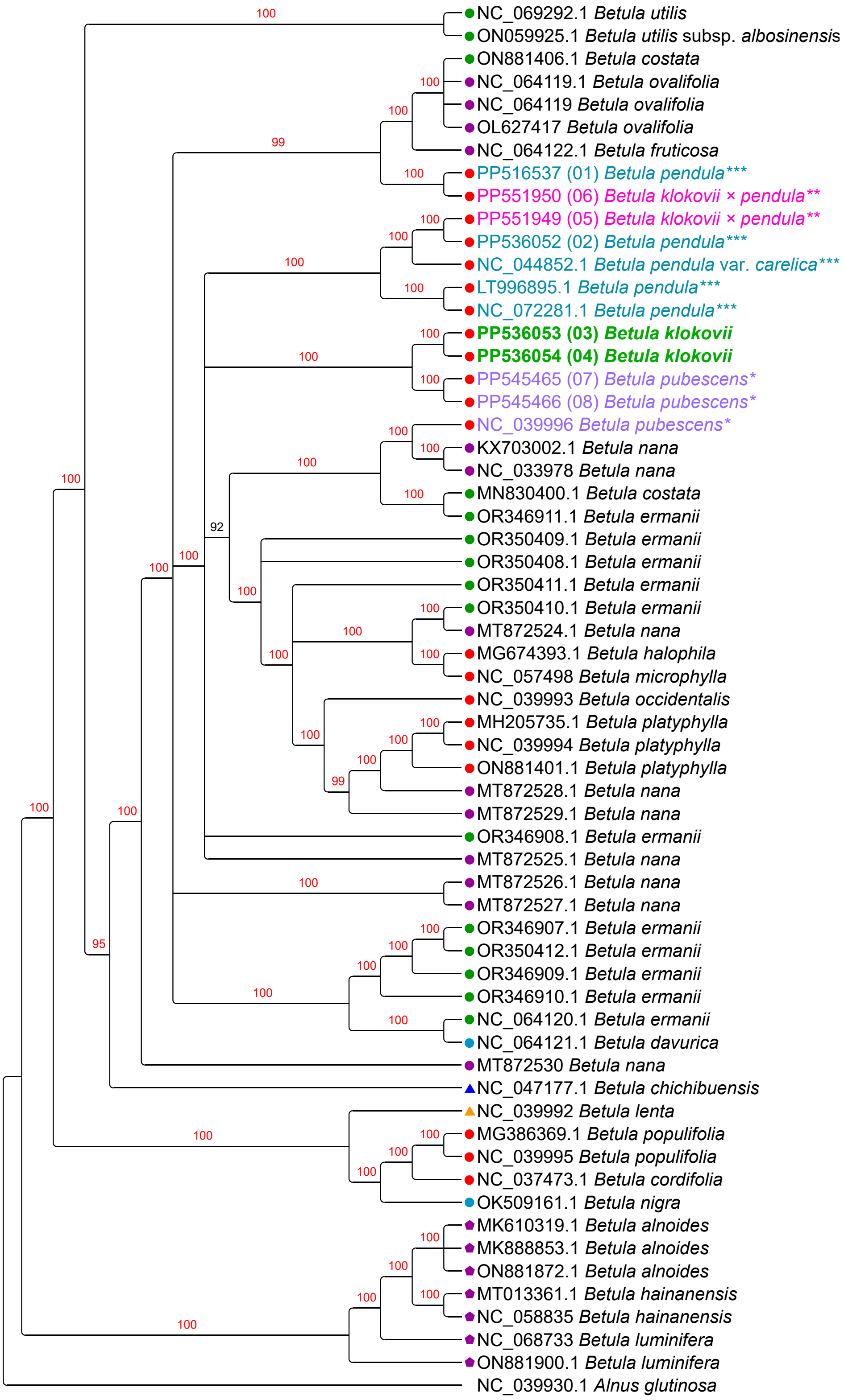


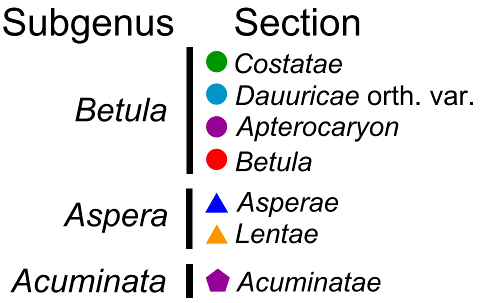


**Fig. S38** Plastome phylogeny based on unique plastome assemblies only (duplicates removed) with partitioning, models assignment according to Akaike information criterion (AIC), Bayesian Inference, MrBayes, 100 000 000 iterations with 10% burnin. Posterior probability values of supported clades are marked red. *B. klokovii* is colored in green and written in bold, presumed *B. pendula* × *B. klokovii* hybrids marked by double asterisk (**) and colored in magenta, *B. pubescens* is marked by asterisk (*) and colored in violet, *B. pendula* and its intraspecific taxa is marked by triple asterisk (***) and colored in cyan. Newly sequenced samples are indicated by numbers 01–08. Subgenera and sections are marked according to Wang & al., 2016.

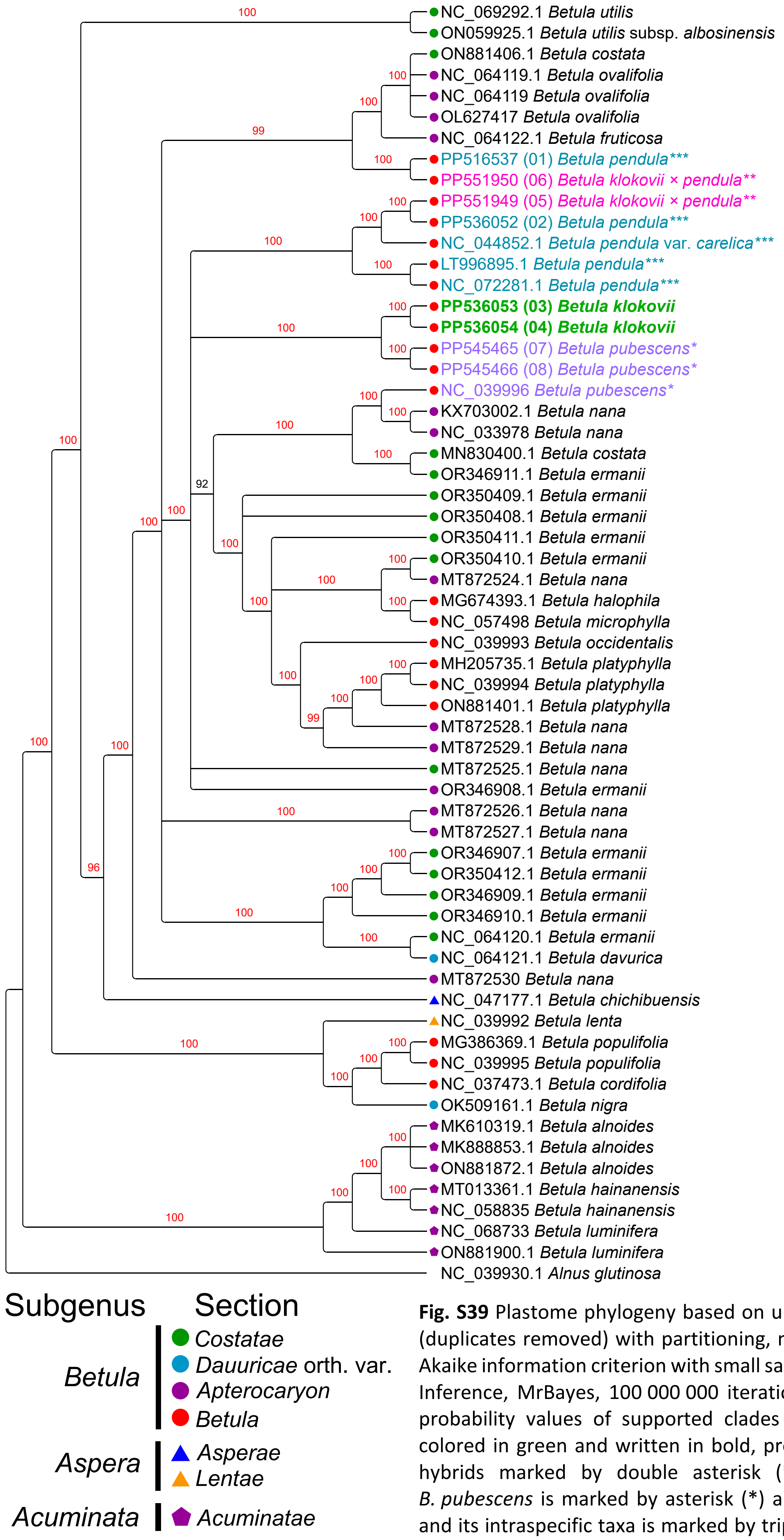


**Fig. S39** Plastome phylogeny based on unique plastome assemblies only (duplicates removed) with partitioning, models assignment according to Akaike information criterion with small sample correction (AICc), Bayesian Inference, MrBayes, 100 000 000 iterations with 10% burnin. Posterior probability values of supported clades are marked red. *B. klokovii* is colored in green and written in bold, presumed *B. pendula* × *B. klokovii* hybrids marked by double asterisk (**) and colored in magenta, *B. pubescens* is marked by asterisk (*) and colored in violet, *B. pendula* and its intraspecific taxa is marked by triple asterisk (***) and colored in cyan. Newly sequenced samples are indicated by numbers 01–08. Subgenera and sections are marked according to Wang et al., 2016.

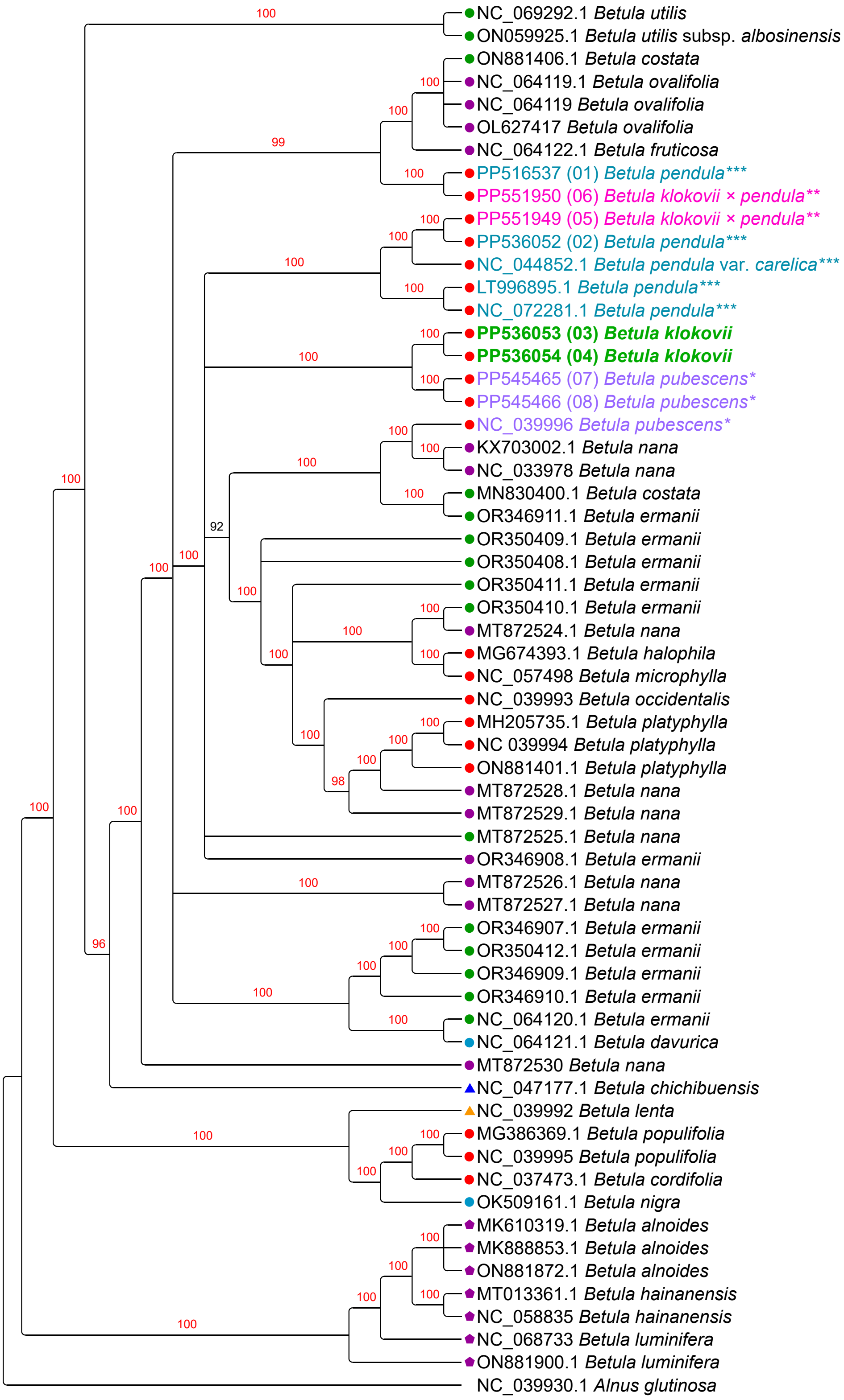


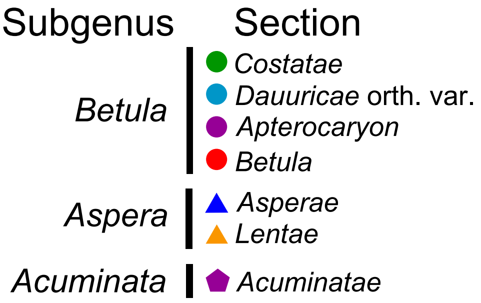


**Fig. S40** Plastome phylogeny reconstruction based on unique plastome assemblies only (duplicates removed) with partitioning, models assignment according to Bayesian Information Criterion (BIC), Bayesian Inference, MrBayes, 100 000 000 iterations with 10% burnin. Posterior probability values of supported clades are marked red. *B. klokovii* is colored in green and written in bold, presumed *B. pendula* × *B. klokovii* hybrids marked by double asterisk (**) and colored in magenta, *B. pubescens* is marked by asterisk (*) and colored in violet, *B. pendula* and its intraspecific taxa is marked by triple asterisk (***) and colored in cyan. Newly sequenced samples are indicated by numbers 01–08. Subgenera and sections are marked according to Wang et al., 2016.

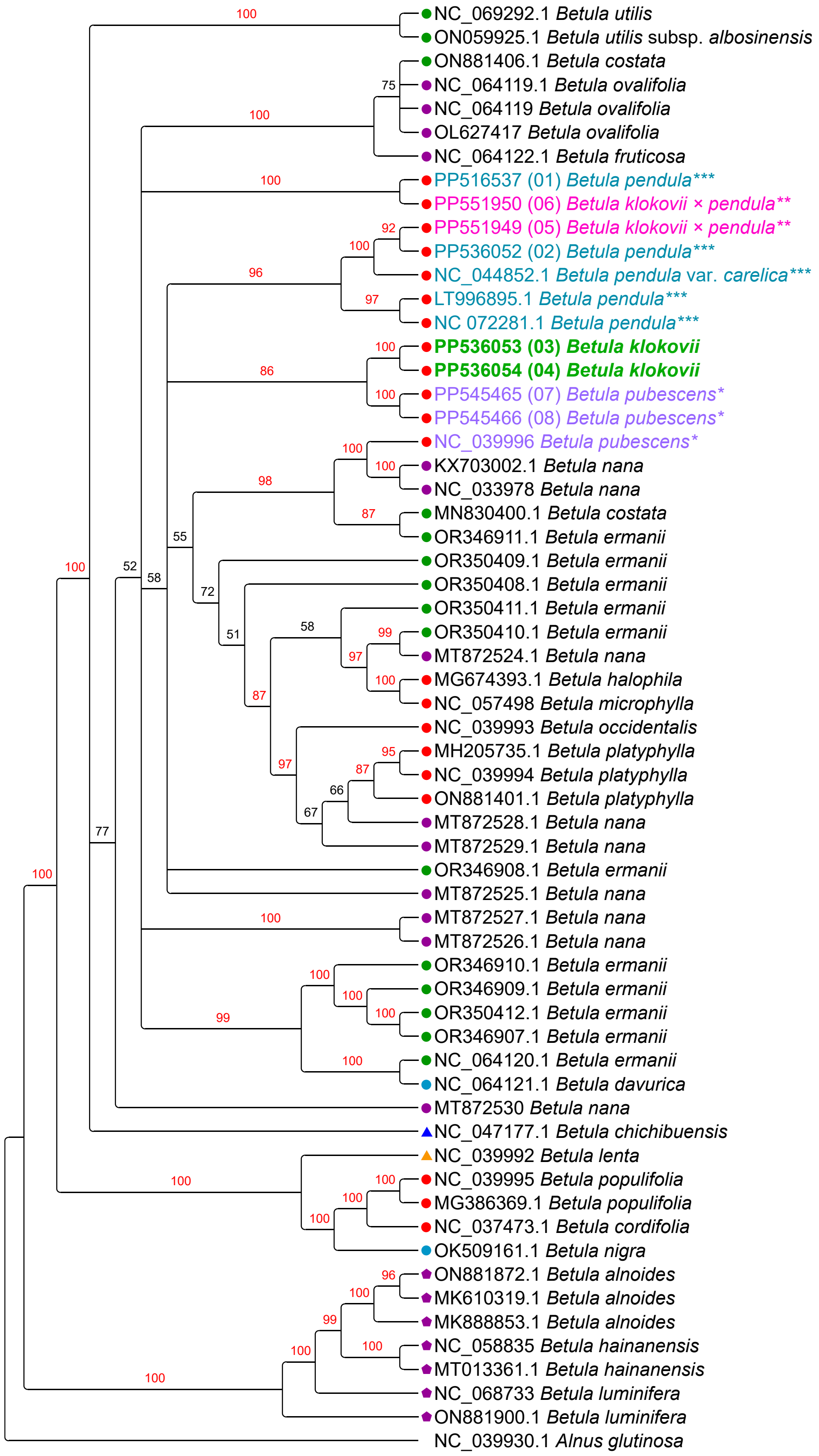


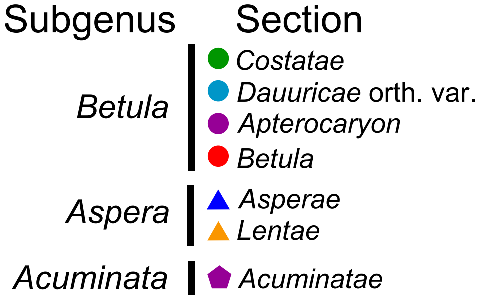


**Fig. S41** Plastome phylogeny based on unique plastome assemblies only (duplicates removed) without partitioning, Maximum Likelihood, RAxML, TVM+I+G, 10 000 iterations bootstrap. Bootstrap values for highly supported clades are marked red. *B. klokovii* is colored in green and written in bold, presumed *B. pendula* × *B. klokovii* hybrids marked by double asterisk (**) and colored in magenta, *B. pubescens* is marked by asterisk (*) and colored in violet, *B. pendula* and its intraspecific taxa is marked by triple asterisk (***) and colored in cyan. Newly sequenced samples are indicated by numbers 01–08. Subgenera and sections are marked according to Wang & al., 2016.

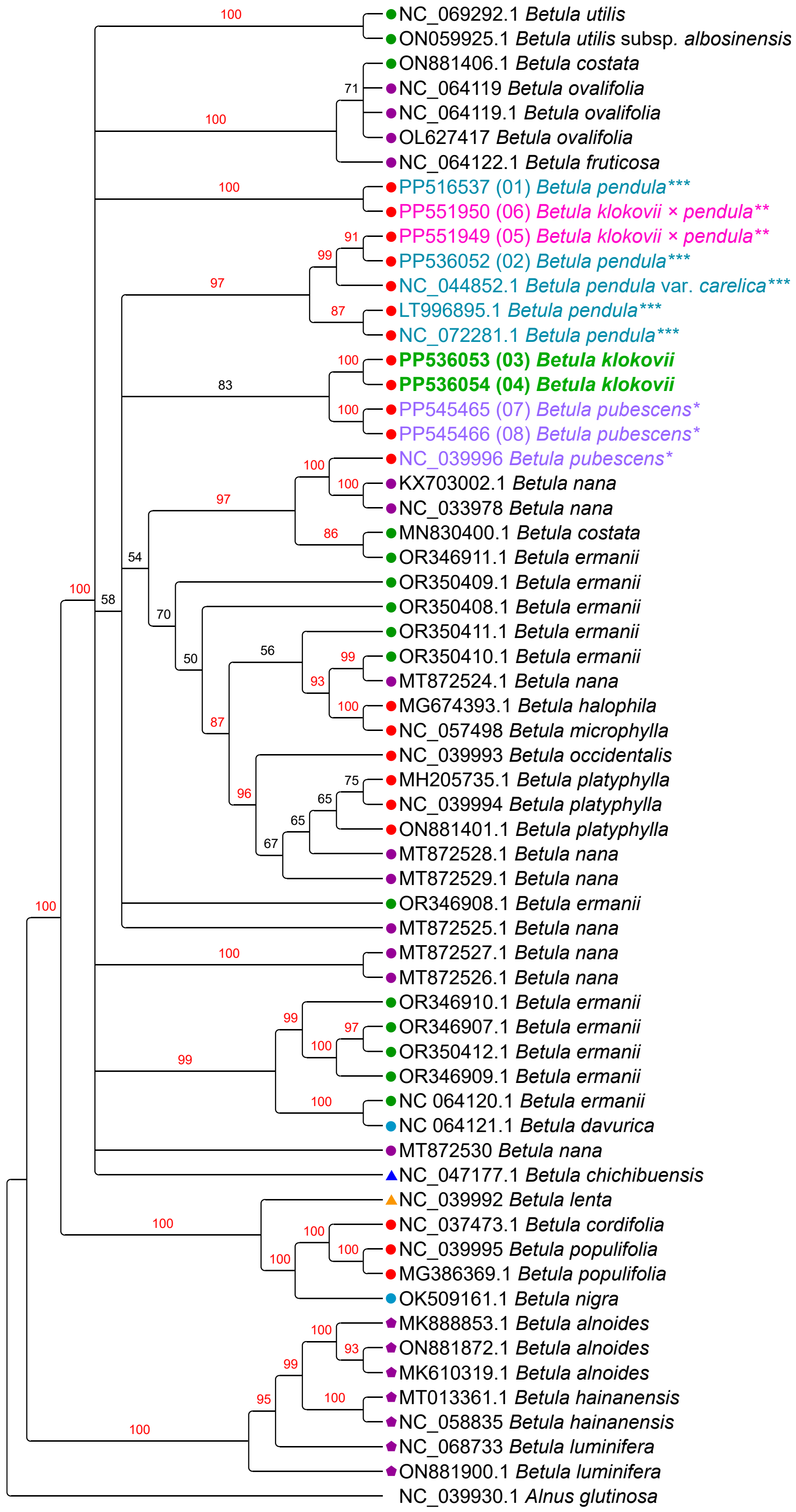


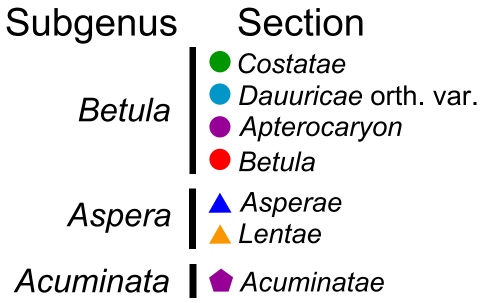


**Fig. S42** Plastome phylogeny based on unique plastome assemblies only (duplicates removed) with partitioning, models assignment according to Akaike information criterion (AIC), Maximum Likelihood, RAxML, 10 000 iterations bootstrap. Bootstrap values for highly supported clades are marked red. *B. klokovii* is colored in green and written in bold, presumed *B. pendula* × *B. klokovii* hybrids marked by double asterisk (**) and colored in magenta, *B. pubescens* is marked by asterisk (*) and colored in violet, *B. pendula* and its intraspecific taxa is marked by triple asterisk (***) and colored in cyan. Newly sequenced samples are indicated by numbers 01–08. Subgenera and sections are marked according to Wang & al., 2016.

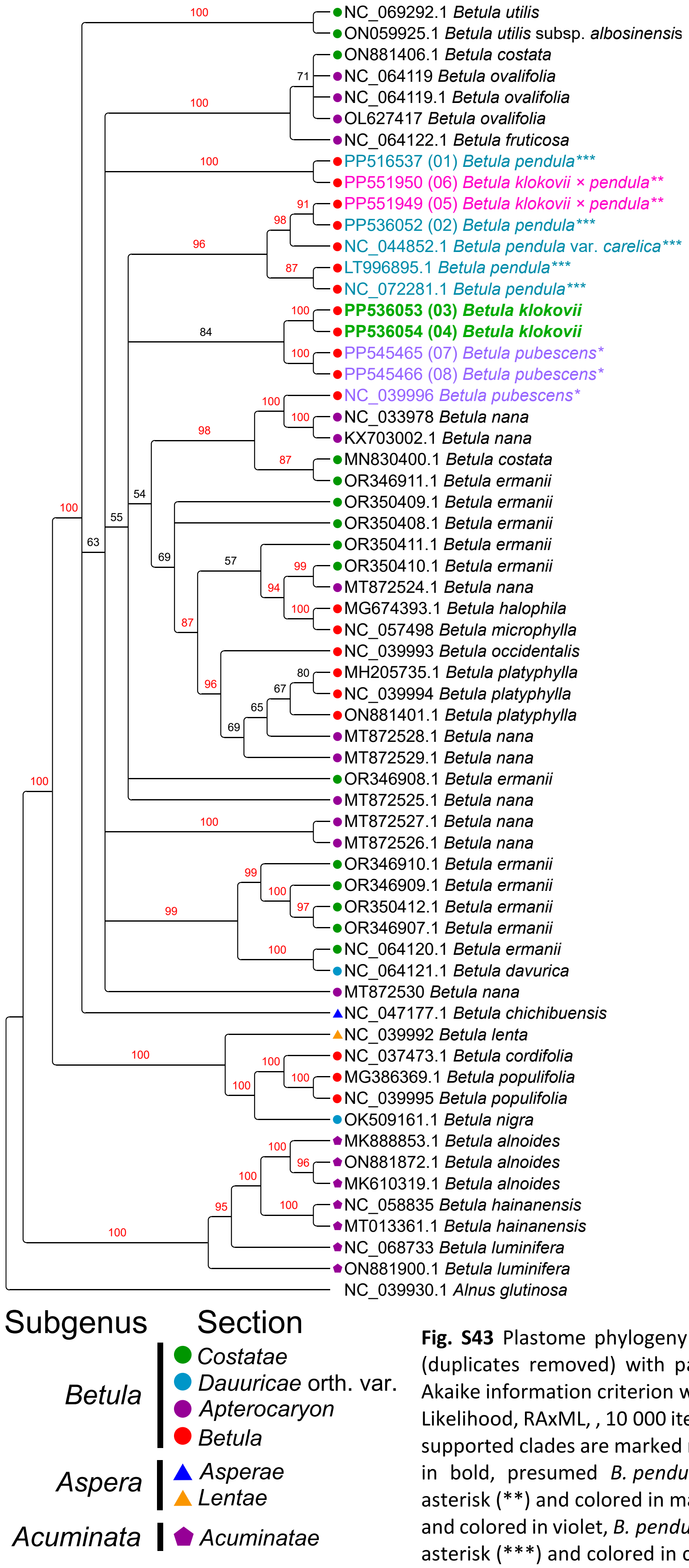


**Fig. S43** Plastome phylogeny based on unique plastome assemblies only (duplicates removed) with partitioning, models assignment according to Akaike information criterion with small sample correction (AICc), Maximum Likelihood, RAxML, , 10 000 iterations bootstrap. Bootstrap values for highly supported clades are marked red. *B. klokovii* is colored in green and written in bold, presumed *B. pendula* × *B. klokovii* hybrids marked by double asterisk (**) and colored in magenta, *B. pubescens* is marked by asterisk (*) and colored in violet, *B. pendula* and its intraspecific taxa is marked by triple asterisk (***) and colored in cyan. Newly sequenced samples are indicated by numbers 01–08. Subgenera and sections are marked according to Wang & al., 2016.

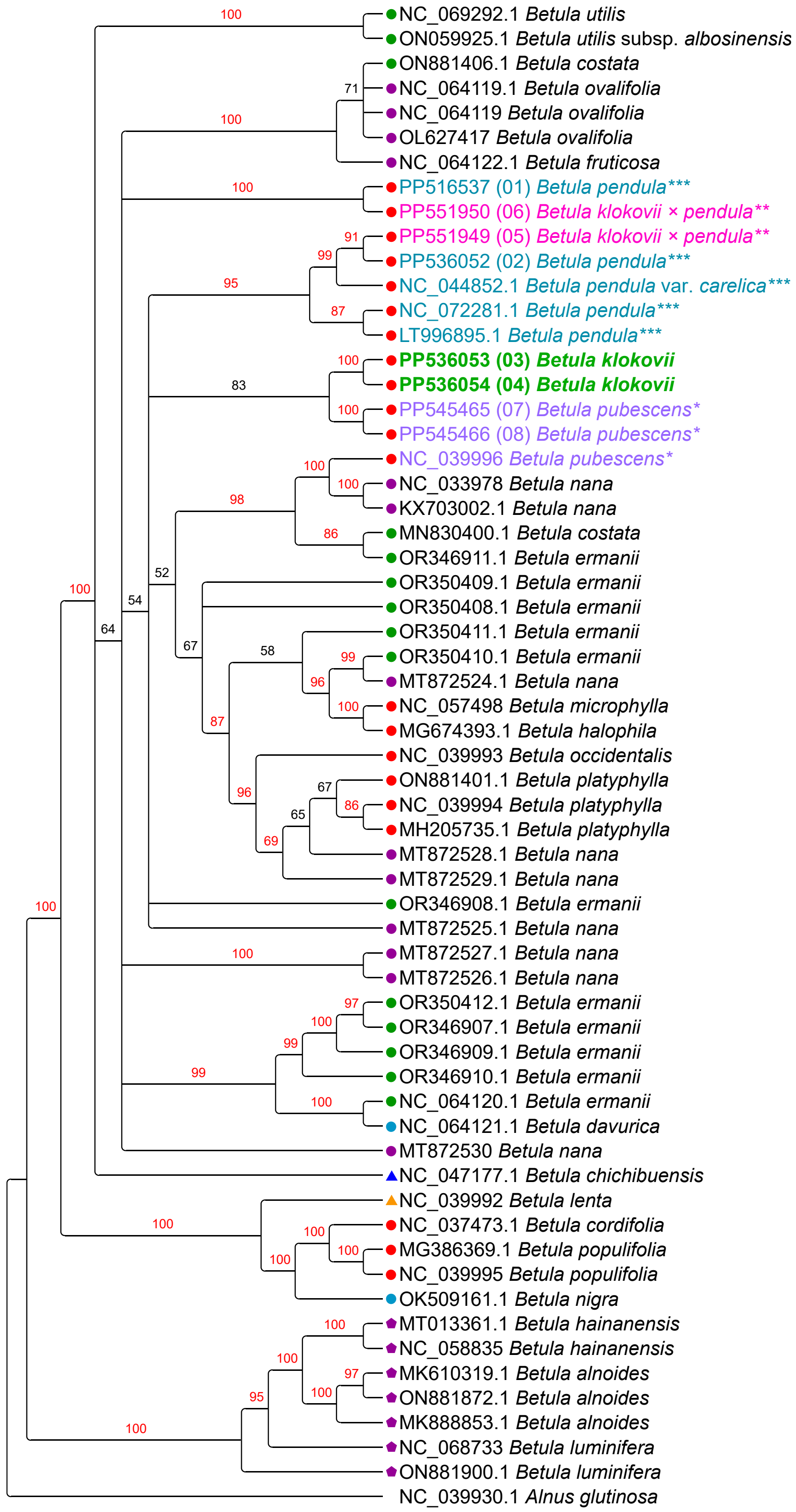


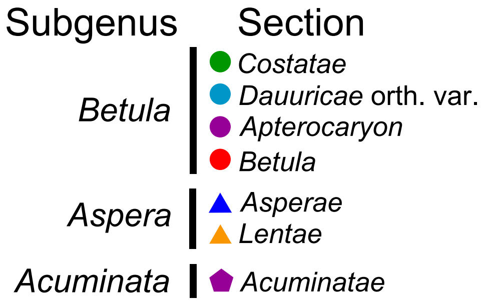


**Fig. S44** Plastome phylogeny based on unique plastome assemblies only (duplicates removed) with partitioning, models assignment according to Bayesian information criterion (BIC), Maximum Likelihood, RAxML, 10 000 iterations bootstrap. Bootstrap values for highly supported clades are marked red. *B. klokovii* is colored in green and written in bold, presumed *B. pendula* × *B. klokovii* hybrids marked by double asterisk (**) and colored in magenta, *B. pubescens* is marked by asterisk (*) and colored in violet, *B. pendula* and its intraspecific taxa is marked by triple asterisk (***) and colored in cyan. Newly sequenced samples are indicated by numbers 01–08. Subgenera and sections are marked according to Wang & al., 2016.

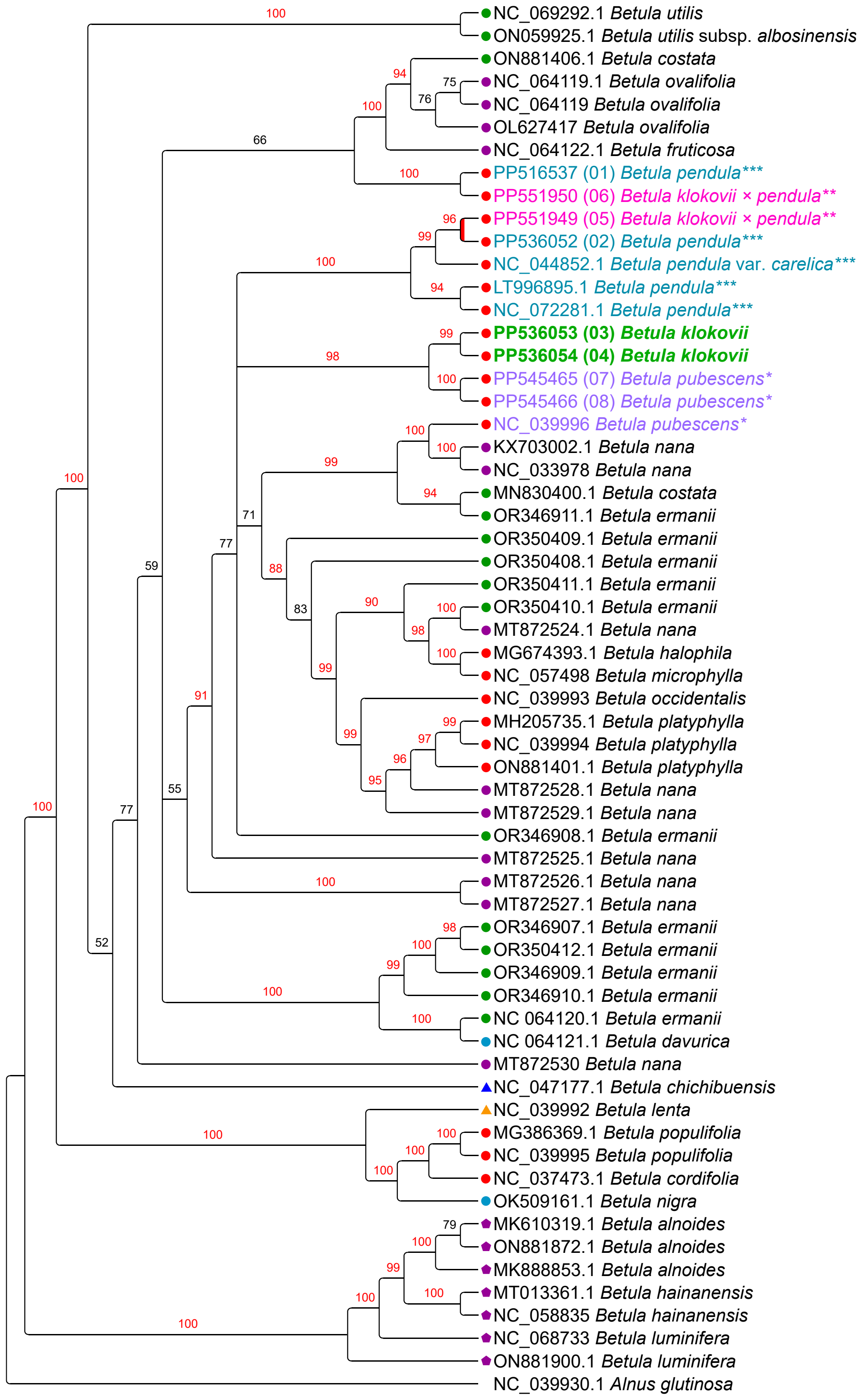

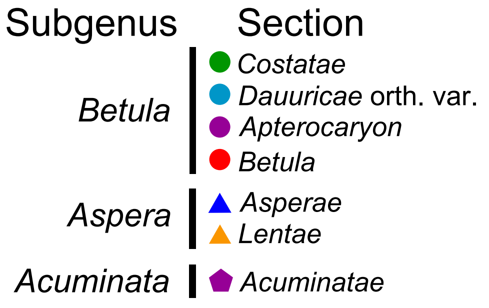


**Fig. S45** Plastome phylogeny based on unique plastome assemblies only (duplicates removed) without partitioning, Maximum Likelihood, IQ-TREE1, TVM+I+G, 10 000 iterations ultrafast bootstrap. Bootstrap values for highly supported clades are marked red. *B. klokovii* is colored in green and written in bold, presumed *B. pendula* × *B. klokovii* hybrids marked by double asterisk (**) and colored in magenta, *B. pubescens* is marked by asterisk (*) and colored in violet, *B. pendula* and its intraspecific taxa is marked by triple asterisk (***) and colored in cyan. Newly sequenced samples are indicated by numbers 01–08. Subgenera and sections are marked according to Wang & al., 2016.

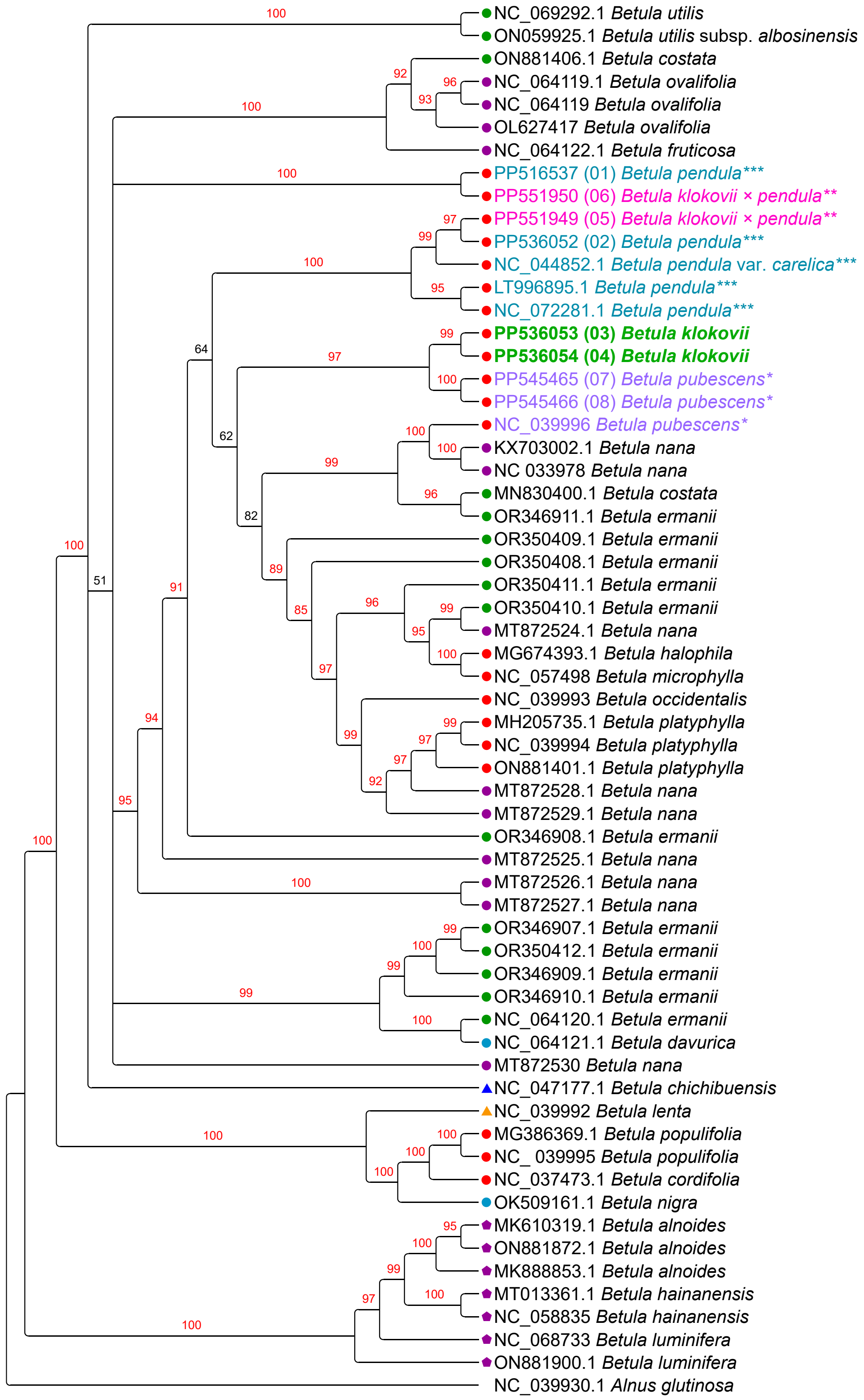


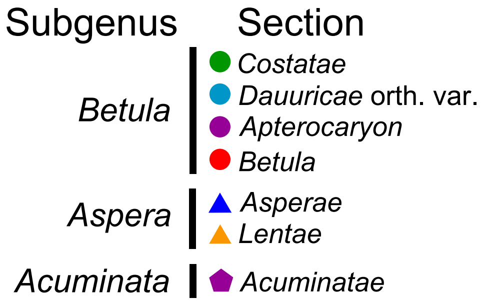


**Fig. S46** Plastome phylogeny based on unique plastome assemblies only (duplicates removed) with partitioning, models assignment according to Akaike information criterion (AIC) Maximum Likelihood, IQ-TREE1, 10 000 iterations ultrafast bootstrap. Bootstrap values for highly supported clades are marked red. *B. klokovii* is colored in green and written in bold, presumed *B. pendula* × *B. klokovii* hybrids marked by double asterisk (**) and colored in magenta, *B. pubescens* is marked by asterisk (*) and colored in violet, *B. pendula* and its intraspecific taxa is marked by triple asterisk (***) and colored in cyan. Newly sequenced samples are indicated by numbers 01–08. Subgenera and sections are marked according to Wang & al., 2016.

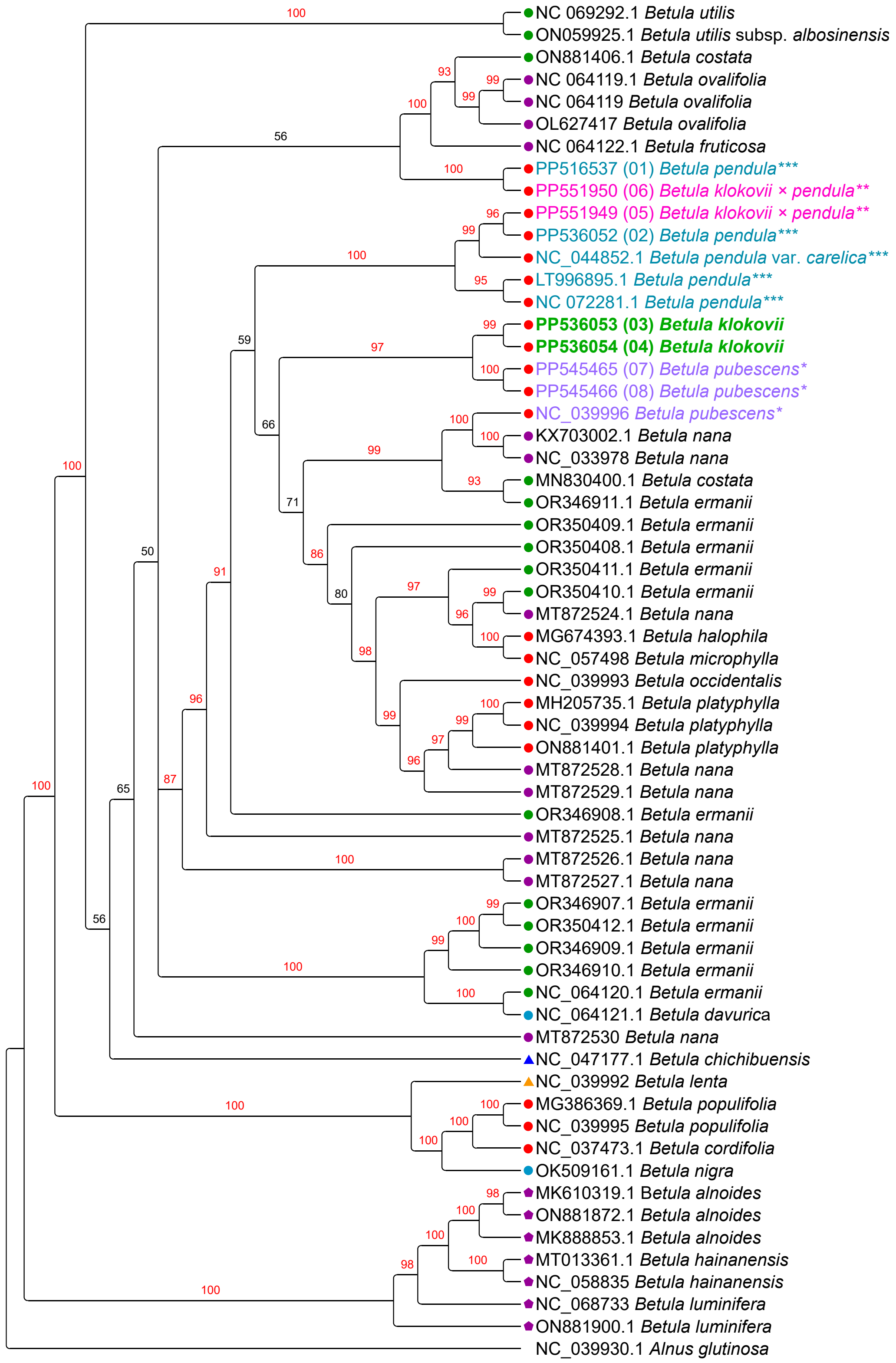


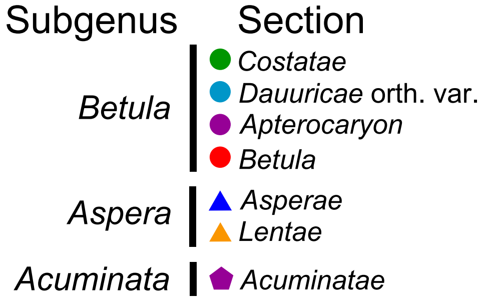


**Fig. S47** Plastome phylogeny based on unique plastome assemblies only (duplicates removed) with partitioning, models assignment according to Akaike information criterion with small sample correction (AICc), Maximum Likelihood, IQ-TREE1, 10 000 iterations ultrafast bootstrap. Bootstrap values for highly supported clades are marked red. *B. klokovii* is colored in green and written in bold, presumed *B. pendula* × *B. klokovii* hybrids marked by double asterisk (**) and colored in magenta, *B. pubescens* is marked by asterisk (*) and colored in violet, *B. pendula* and its intraspecific taxa is marked by triple asterisk (***) and colored in cyan. Newly sequenced samples are indicated by numbers 01–08. Subgenera and sections are marked according to Wang & al., 2016.

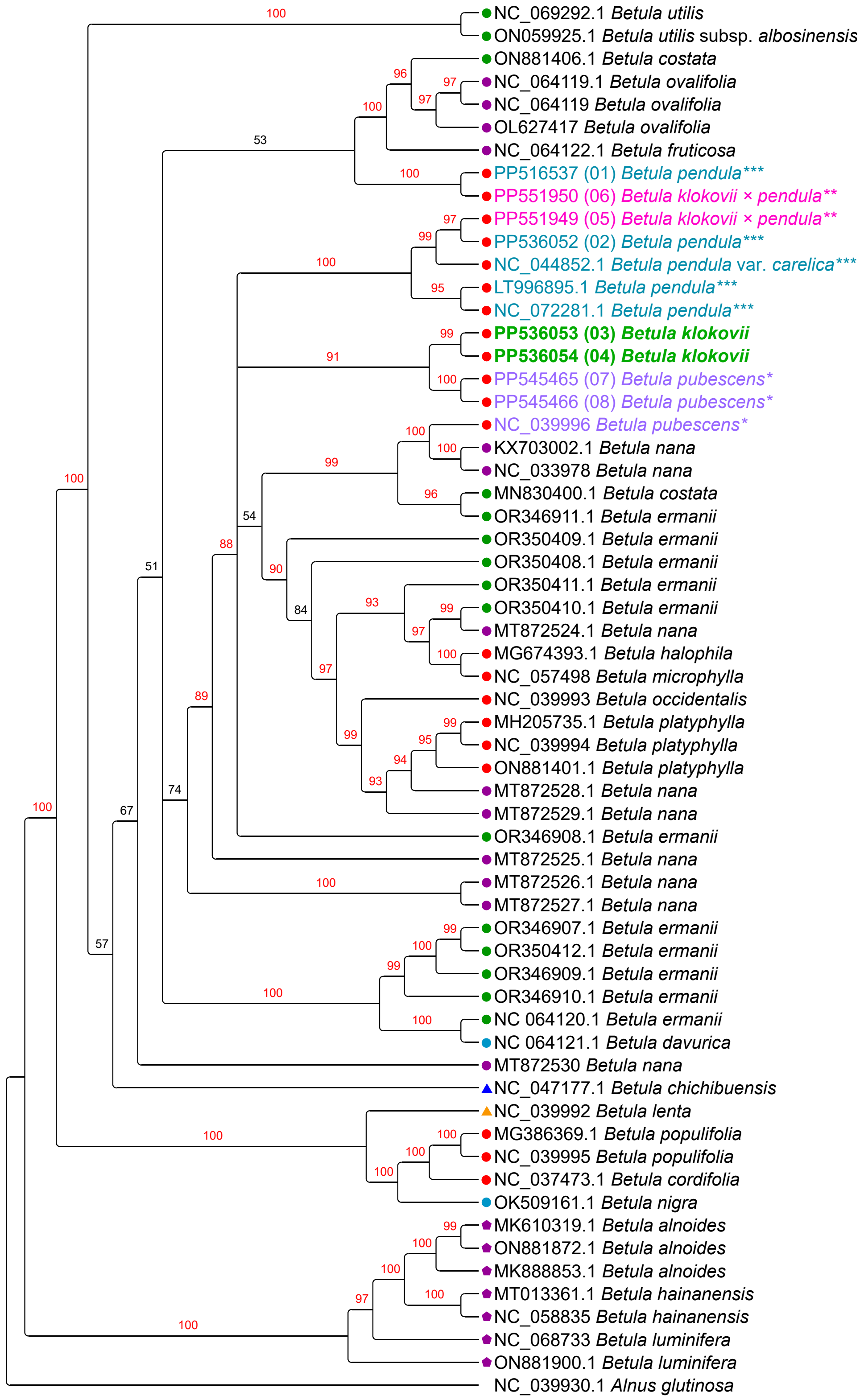


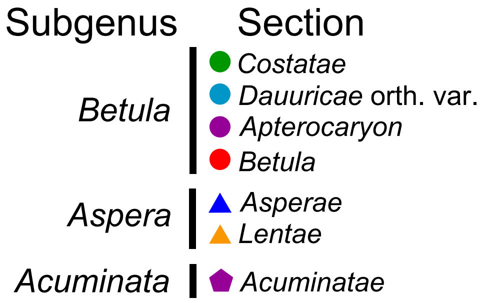


**Fig. S48** Plastome phylogeny based on unique plastome assemblies only (duplicates removed) with partitioning, models assignment according to Bayesian information criterion (BIC), Maximum Likelihood, IQ-TREE1, 10 000 iterations ultrafast bootstrap. Bootstrap values for highly supported clades are marked red. *B. klokovii* is colored in green and written in bold, presumed *B. pendula* × *B. klokovii* hybrids marked by double asterisk (**) and colored in magenta, *B. pubescens* is marked by asterisk (*) and colored in violet, *B. pendula* and its intraspecific taxa is marked by triple asterisk (***) and colored in cyan. Newly sequenced samples are indicated by numbers 01–08. Subgenera and sections are marked according to Wang & al., 2016.

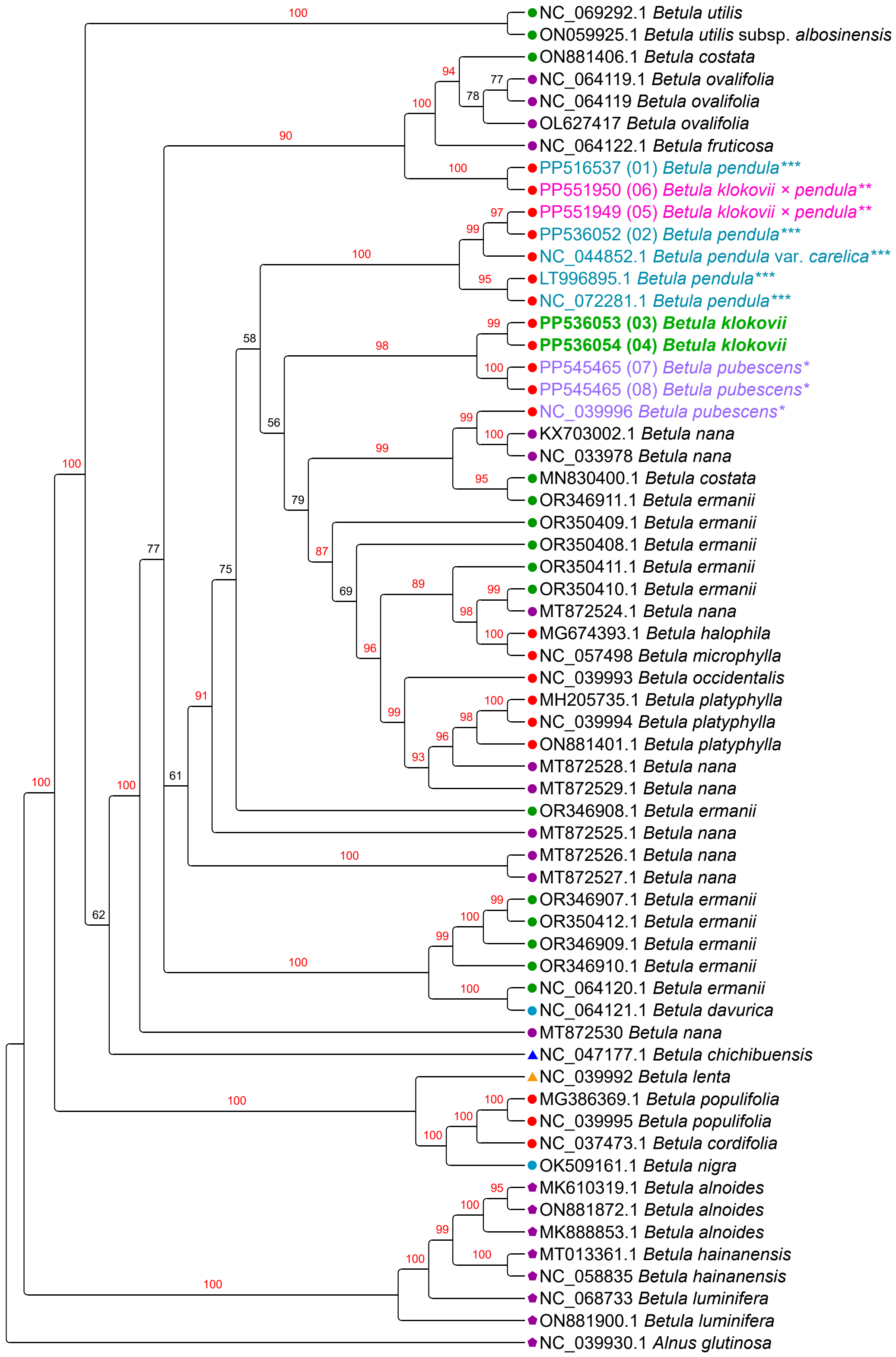


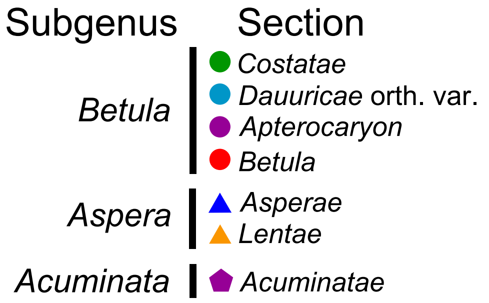


**Fig. S49** Plastome phylogeny based on unique plastome assemblies only (duplicates removed) without partitioning, Maximum Likelihood, IQ-TREE2, TVM+I+G, 10 000 iterations ultrafast bootstrap. Bootstrap values for highly supported clades are marked red. *B. klokovii* is colored in green and written in bold, presumed *B. pendula* × *B. klokovii* hybrids marked by double asterisk (**) and colored in magenta, *B. pubescens* is marked by asterisk (*) and colored in violet, *B. pendula* and its intraspecific taxa is marked by triple asterisk (***) and colored in cyan. Newly sequenced samples are indicated by numbers 01–08. Subgenera and sections are marked according to Wang & al., 2016.

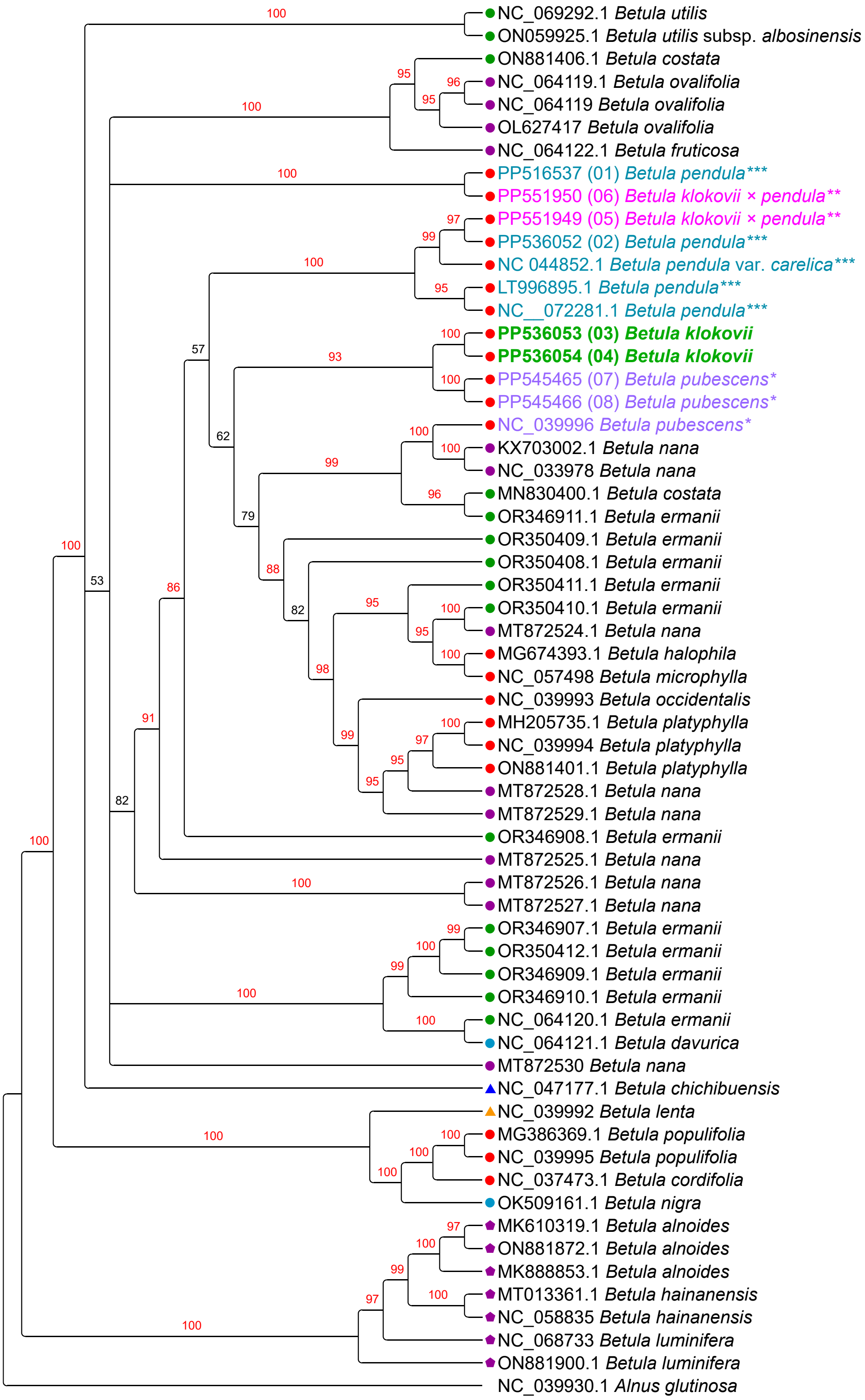


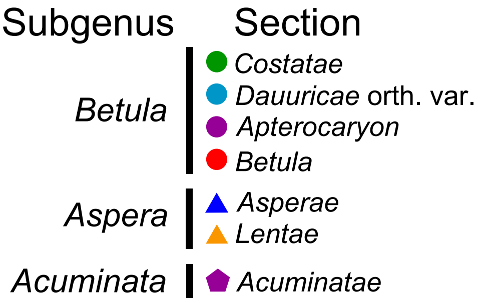


**Fig. S50** Plastome phylogeny based on unique plastome assemblies only (duplicates removed) with partitioning, models assignment according to Akaike information criterion (AIC), Maximum Likelihood, IQ-TREE2, 10 000 iterations ultrafast bootstrap. Bootstrap values for highly supported clades are marked red. *B. klokovii* is colored in green and written in bold, presumed *B. pendula* × *B. klokovii* hybrids marked by double asterisk (**) and colored in magenta, *B. pubescens* is marked by asterisk (*) and colored in violet, *B. pendula* and its intraspecific taxa is marked by triple asterisk (***) and colored in cyan. Newly sequenced samples are indicated by numbers 01–08. Subgenera and sections are marked according to Wang & al., 2016.

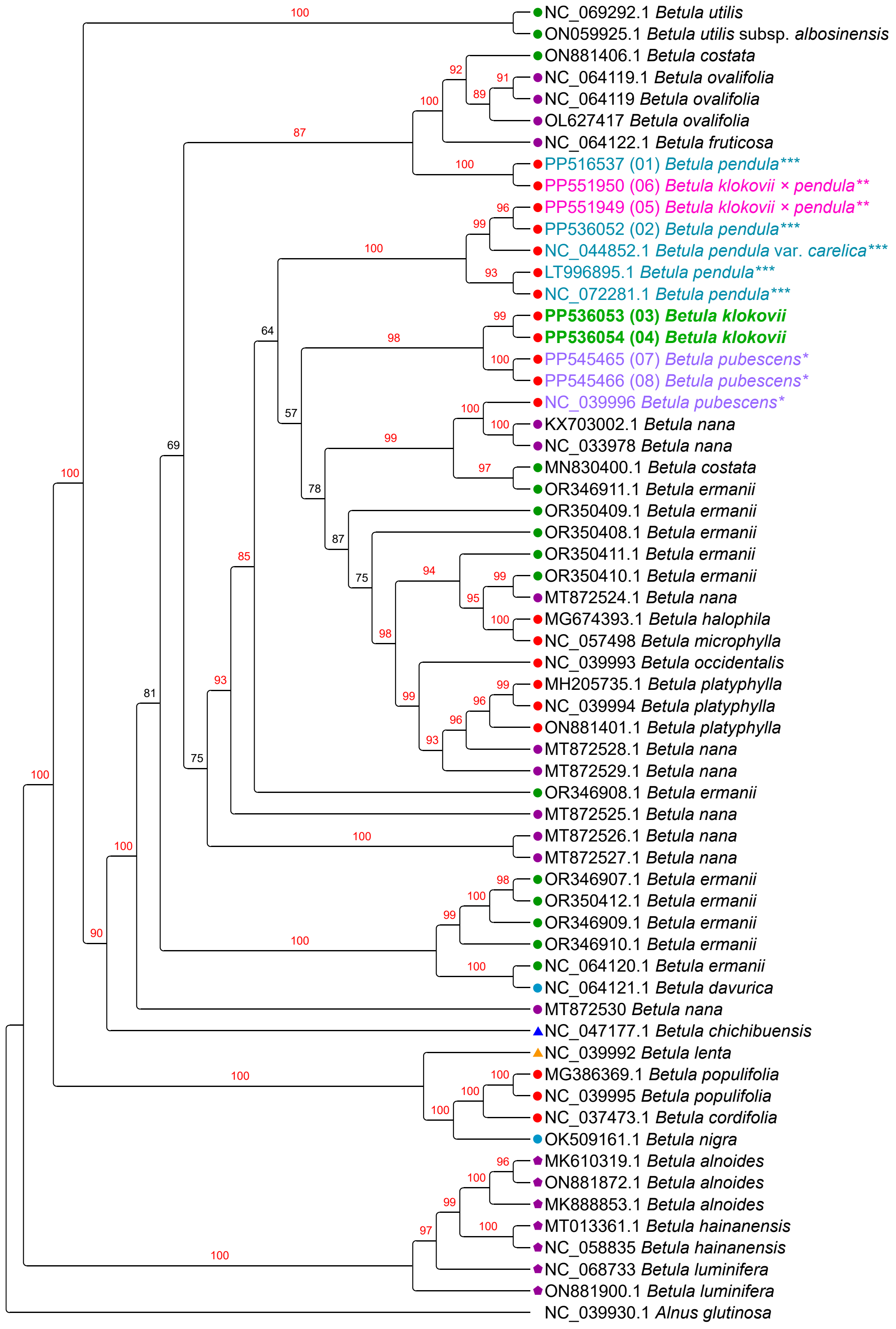

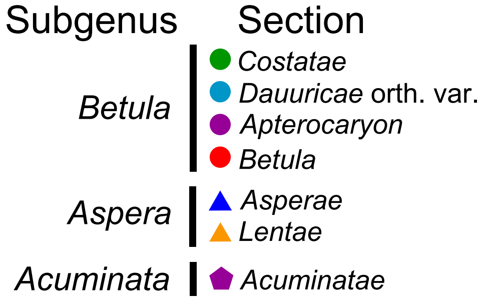


**Fig. S51** Plastome phylogeny based on unique plastome assemblies only (duplicates removed) with partitioning, models assignment according to Akaike information criterion with small sample correction (AICc), Maximum Likelihood, IQ-TREE2, 10 000 iterations ultrafast bootstrap. Bootstrap values for highly supported clades are marked red. *B. klokovii* is colored in green and written in bold, presumed *B. pendula* × *B. klokovii* hybrids marked by double asterisk (**) and colored in magenta, *B. pubescens* is marked by asterisk (*) and colored in violet, *B. pendula* and its intraspecific taxa is marked by triple asterisk (***) and colored in cyan. Newly sequenced samples are indicated by numbers 01–08. Subgenera and sections are marked according to Wang & al., 2016.

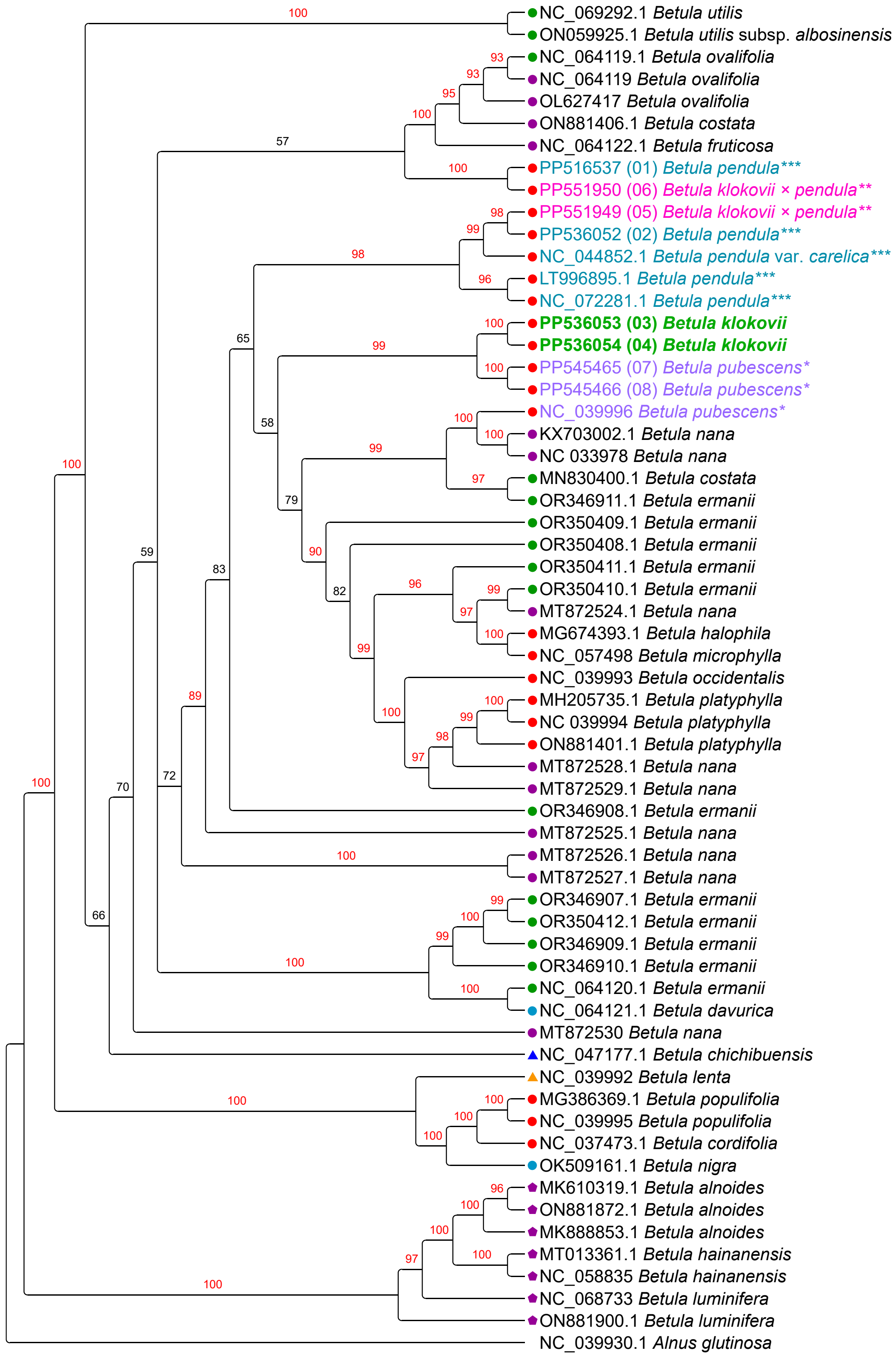

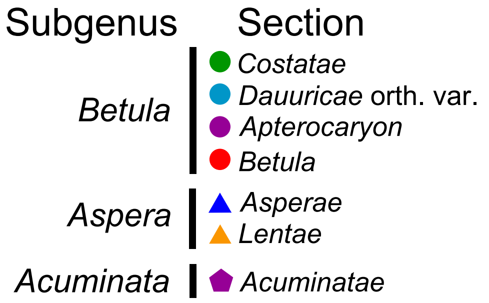


**Fig. S52** Plastome phylogeny based on unique plastome assemblies only (duplicates removed) with partitioning, models assignment according to Bayesian information criterion (BIC), Maximum Likelihood, IQ-TREE2, 10 000 iterations ultrafast bootstrap. Bootstrap values for highly supported clades are marked red. *B. klokovii* is colored in green and written in bold, presumed *B. pendula* × *B. klokovii* hybrids marked by double asterisk (**) and colored in magenta, *B. pubescens* is marked by asterisk (*) and colored in violet, *B. pendula* and its intraspecific taxa is marked by triple asterisk (***) and colored in cyan. Newly sequenced samples are indicated by numbers 01–08. Subgenera and sections are marked according to Wang & al., 2016.

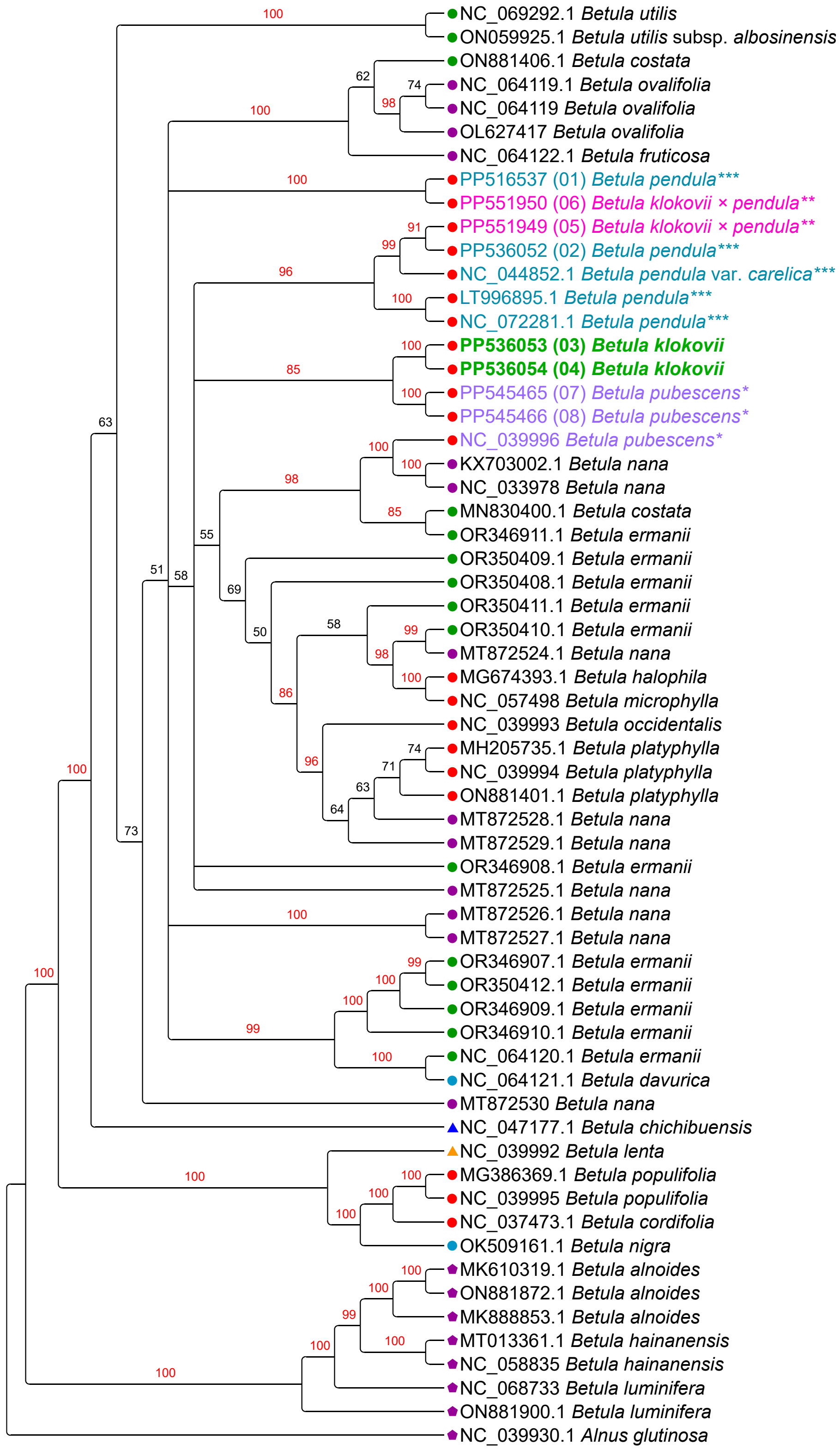


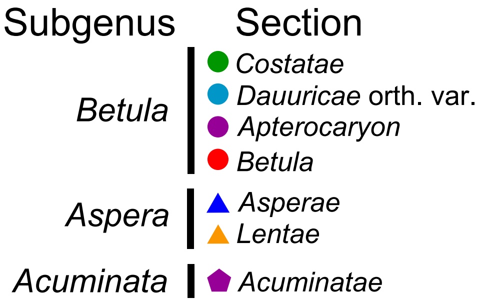


**Fig. S53** Plastome phylogeny based on unique plastome assemblies only (duplicates removed) without partitioning, Maximum Likelihood, IQ-TREE2, TVM+I+G, 1000 iterations nonparametric bootstrap. Bootstrap values for highly supported clades are marked red. *B. klokovii* is colored in green and written in bold, presumed *B. pendula* × *B. klokovii* hybrids marked by double asterisk (**) and colored in magenta, *B. pubescens* is marked by asterisk (*) and colored in violet, *B. pendula* and its intraspecific taxa is marked by triple asterisk (***) and colored in cyan. Newly sequenced samples are indicated by numbers 01–08. Subgenera and sections are marked according to Wang & al., 2016.

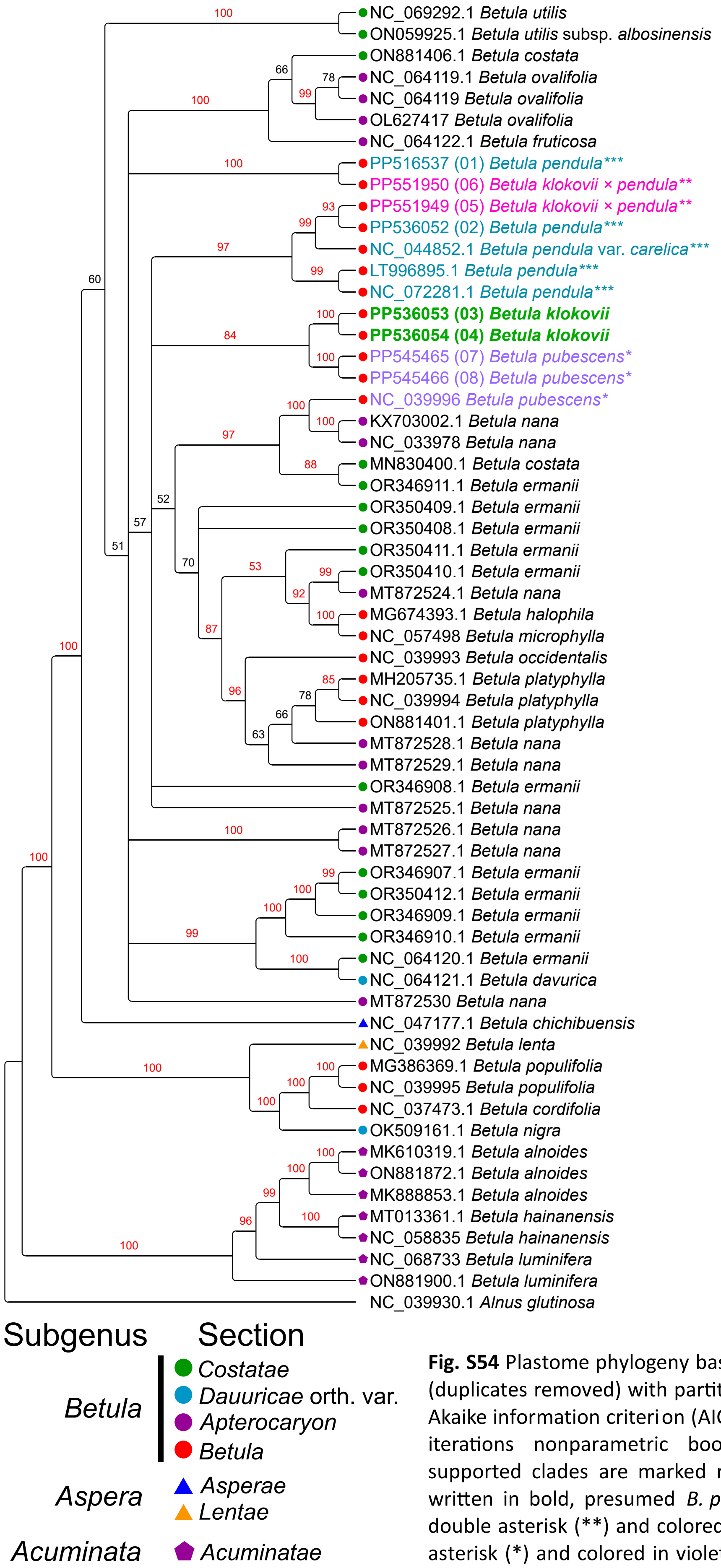


**Fig. S54** Plastome phylogeny based on unique plastome assemblies only (duplicates removed) with partitioning, models assignment according to Akaike information criterion (AIC), Maximum Likelihood, IQ-TREE2, 1000 iterations nonparametric bootstrap. Bootstrap values for highly supported clades are marked red. *B. klokovii* is colored in green and written in bold, presumed *B. pendula* × *B. klokovii* hybrids marked by double asterisk (**) and colored in magenta, *B. pubescens* is marked by asterisk (*) and colored in violet, *B. pendula* and its intraspecific taxa is marked by triple asterisk (***) and colored in cyan. Newly sequenced samples are indicated by numbers 01–08. Subgenera and sections are marked according to Wang & al., 2016.

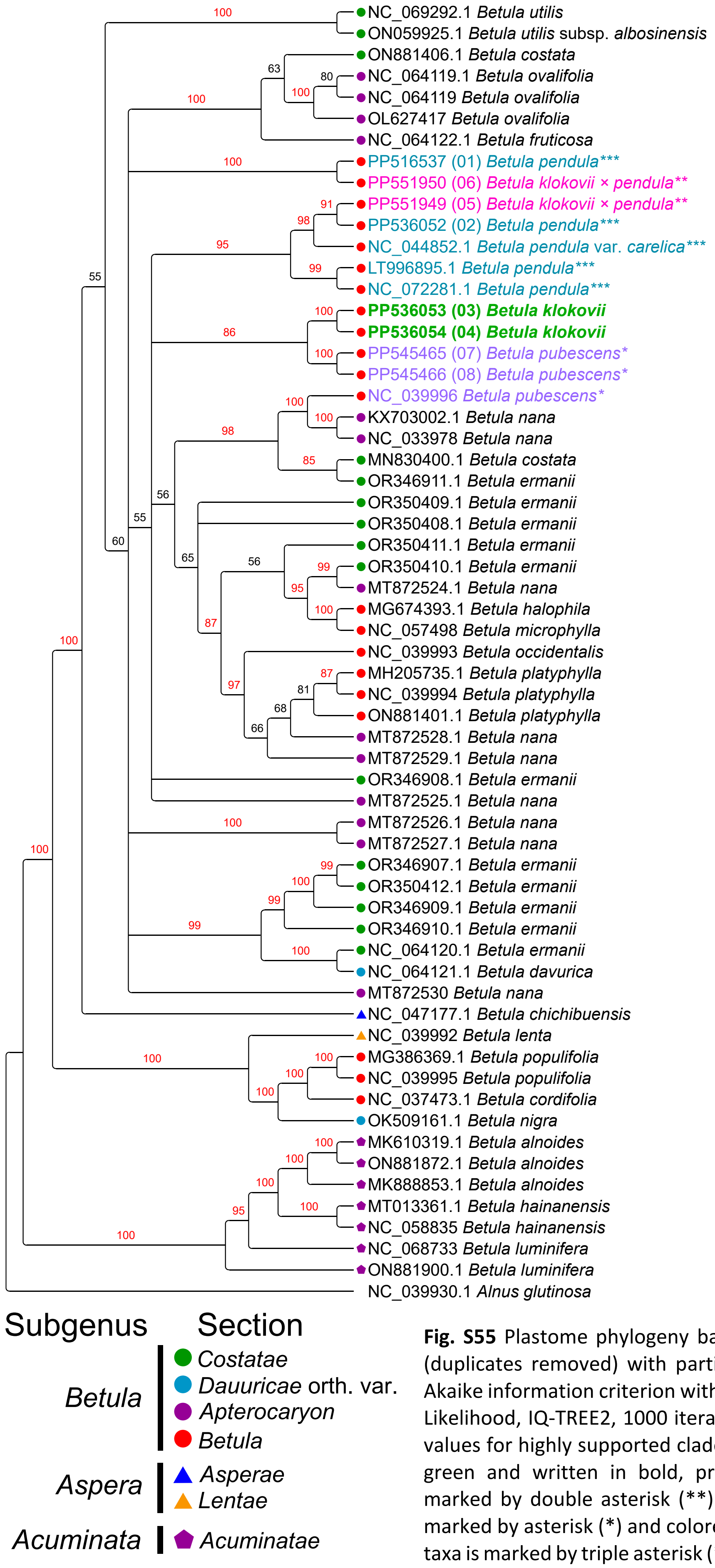


**Fig. S55** Plastome phylogeny based on unique plastome assemblies only (duplicates removed) with partitioning, models assignment according to Akaike information criterion with small sample correction (AICc), Maximum Likelihood, IQ-TREE2, 1000 iterations nonparametric bootstrap. Bootstrap values for highly supported clades are marked red. *B. klokovii* is colored in green and written in bold, presumed *B. pendula* × *B. klokovii* hybrids marked by double asterisk (**) and colored in magenta, *B. pubescens* is marked by asterisk (*) and colored in violet, *B. pendula* and its intraspecific taxa is marked by triple asterisk (***) and colored in cyan. Newly sequenced samples are indicated by numbers 01–08. Subgenera and sections are marked according to Wang & al., 2016.

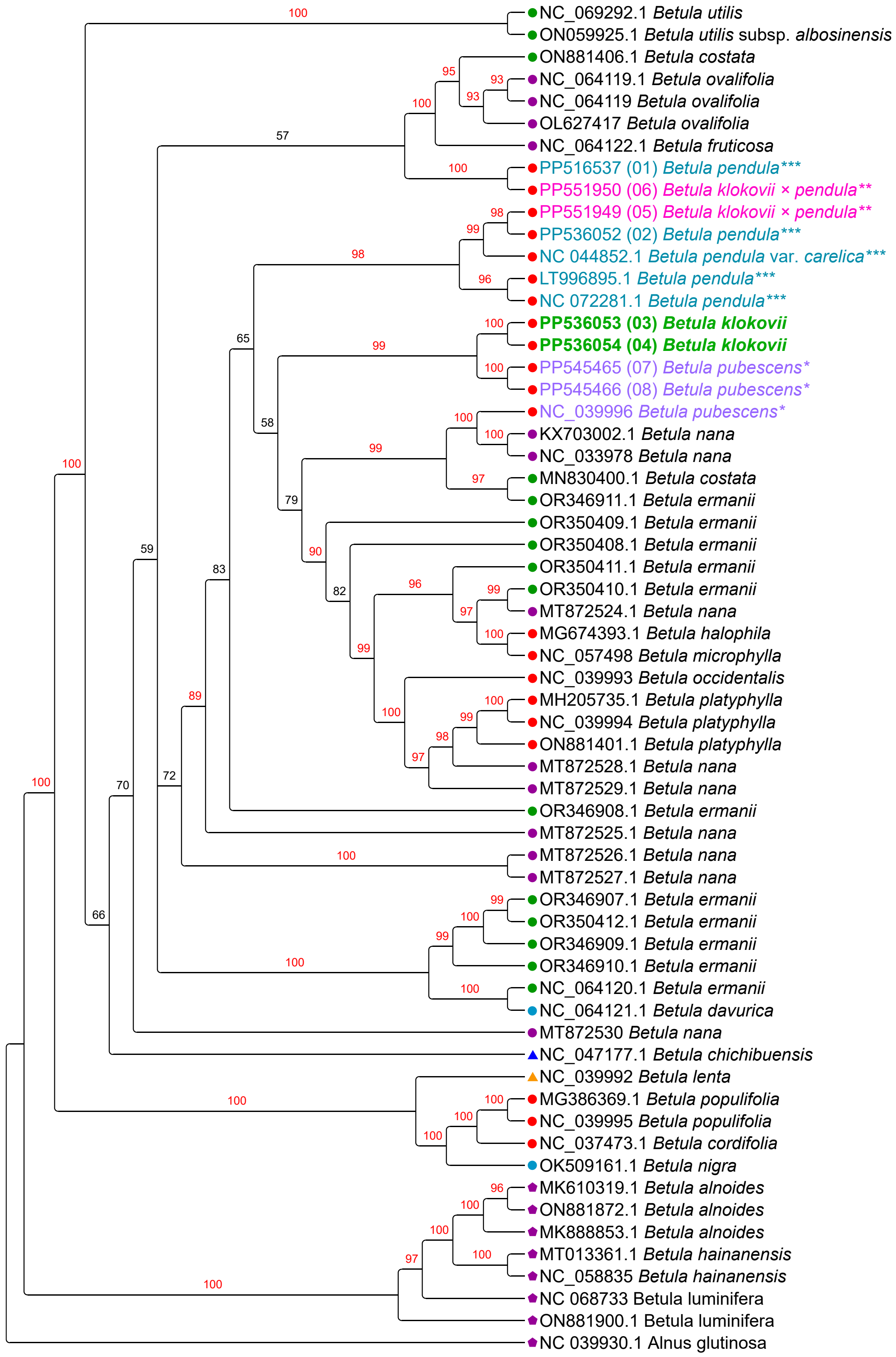


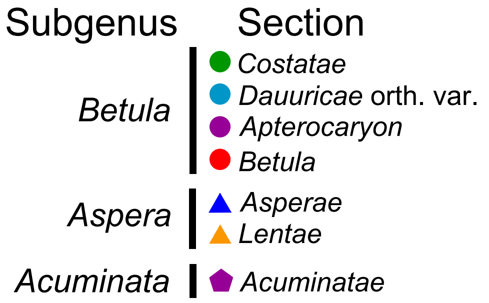


**Fig. S56** Plastome phylogeny reconstruction based on unique plastome assemblies only (duplicates removed) with partitioning, models assignment according to Bayesian information criterion (BIC), Maximum Likelihood, IQ-TREE2, 1000 iterations nonparametric bootstrap. Bootstrap values for highly supported clades are marked red. *B. klokovii* is colored in green and written in bold, presumed *B. pendula* × *B. klokovii* hybrids marked by double asterisk (**) and colored in magenta, *B. pubescens* is marked by asterisk (*) and colored in violet, *B. pendula* and its intraspecific taxa is marked by triple asterisk (***) and colored in cyan. Newly sequenced samples are indicated by numbers 01–08. Subgenera and sections are marked according to Wang & al., 2016.
