## Supporting information - Tables S1-S4 for "Phylogenetic and taxonomic insights into *Betula*: low-coverage whole genome sequencing and plastome analysis with focus on the rare Ukrainian endemic species *Betula klokovii* Zaverucha"

**Table S1** Information about analyzed samples

| # | Species | Internal identifier | Collection place | GPS coordinates | Collection date | Collector | Herbarium identifier |
|---|---|---|---|---|---|---|---|
| 01 | *Betula pendula* | UA-KM-3Bp | Ukraine, Ternopil region., Kremenets distr., Natural Nature Park "Kremenets Mountains", mt. Maslyatyn | N50°04.770' E025°39.706' | 16.07.2019 | A. Tarieiev, A. Shtogun, I. Skorobogata | |
| 02 | *Betula pendula* | UA-KS-8Bp | Ukraine, Ternopil region., Kremenets distr., Natural Nature Park "Kremenets Mountains", mt. Strakhova | N50°05.081' E025°40.475' | 16.07.2019 | A. Tarieiev, A. Shtogun, I. Skorobogata | |
| 03 | *Betula klokovii* | UA-KS-9Bk | Ukraine, Ternopil region., Kremenets distr., Natural Nature Park "Kremenets Mountains", mt. Strakhova, old bent tree | N50°05.094' E025°40.456' | 16.07.2019 | A. Tarieiev, A. Shtogun, I. Skorobogata | |
| 04 | *Betula klokovii* | UA-KM-8Bk | Ukraine, Ternopil region., Kremenets distr., Natural Nature Park "Kremenets Mountains", mt. Maslyatyn | N50°04.793' E025°39.680' | 16.07.2019 | A. Tarieiev, A. Shtogun, I. Skorobogata | |
| 05 | *Betula klokovii* × *pendula* | UA-KM-5Bkp | Ukraine, Ternopil region., Kremenets distr., Natural Nature Park "Kremenets Mountains", mt. Maslyatyn | N 50°04.784' E 025°38.681' | 16.07.2019 | A. Tarieiev, A. Shtogun, I. Skorobogata | |
| 06 | *Betula klokovii* × *pendula* | UA-KS-4Bkp | Ukraine, Ternopil region., Kremenets distr., Natural Nature Park "Kremenets Mountains", mt. Strakhova, old tree on the top | N 50°05.491' E 025°40.406' | 15.07.2019 | A. Tarieiev, A. Shtogun, I. Skorobogata | |
| 07 | *Betula pubescens* | UA-DS-3Bp | Ukraine, Sumy region, Desniansko-Starogutsky NNP | | | S.M. Panchenko | |
| 08 | *Betula pubescens* | UA-HL-2Bp | Ukraine, Sumy region, Okhtyrka distr., National Nature Park "Hetmanskyi", "Lytovsky bir" (Національний природний парк «Гетьманський»б литовський бір) | | 08.08.2019 | S.M. Panchenko | |

**Table S2** DNA concentration measurements by NanoDrop

| Sample | Identifier | Species | DNA concentration, ng/µL |
|---|---|---|---|
| 01 | UA-KM-3Bp | *B. pendula* | 9.2 |
| 02 | UA-KS-8Bp | | 6.0 |
| 03 | UA-KS-9Bk | *B. klokovii* | 4.5 |
| 04 | UA-KM-8Bk | | 3.6 |
| 05 | UA-KM-5Bkp | *B. klokovii* × *B. pendula* | 8.2 |
| 06 | UA-KS-3Bkp | | 7.1 |
| 07 | UA-DS-3Bp | *B. pubescens* | 3.6 |
| 08 | UA-HL-2Bp | | 4.8 |

**Table S3** Basic information on mapping using CLC Genomic Workbench 23.0.2

| # sample | Species | Reference (NCBI accession) | Reference length, bp | Contig length, bp | | Average coverage | |
|---|---|---|---|---|---|---|---|
| | | | | Raw reads | Cleaned reads | Raw reads | Cleaned reads |
| 01 | *Betula pendula* | NC_072281.1 | 160552 | 160458 | 151267 | 107,30 | 21,77 |
| 02 | | | | 160511 | 152126 | 135,93 | 31,53 |
| 03 | *Betula klokovii* | NC_039996.1 | 158647 | 158531 | 150179 | 166,13 | 25,63 |
| 04 | | | | 158543 | 146285 | 57,71 | 13,49 |
| 05 | *Betula pendula* × *klokovii* | NC_072281.1 | 160552 | 160524 | 151473 | 127,06 | 23,68 |
| 06 | | | | 160461 | 153437 | 225,45 | 46,20 |
| 07 | *Betula pubescens* | NC_039996.1 | 158647 | 158491 | 149953 | 143,14 | 28,24 |
| 08 | | | | 158418 | 152152 | 470,51 | 76,09 |

**Table S4** Basic information on NOVOPlasty plastid genome assemblies

| # sample | Species | Assembly parameters | | | | | | | | |
|---|---|---|---|---|---|---|---|---|---|---|
| | | k=23 | | Circularized genome assembly, bp | k=33 | | Circularized genome assembly, bp | k=45 | | Circularized genome assembly, bp |
| | | Seed | Reference | | Seed | Reference | | Seed | Reference | |
| *01* | *B. pendula* | OP820513.1 | NC_072281.1 | 160554 | OP820513.1 | NC_072281.1 | 160554 | OP820513.1 | NC_072281.1 | 160554 |
| | | NC_072281.1 | OP820513.1 | 160554 | NC_072281.1 | OP820513.1 | 160554 | NC_072281.1 | OP820513.1 | 160554 |
| | | NC_072281.1 | no reference | - | NC_072281.1 | no reference | - | NC_072281.1 | no reference | - |
| *02* | *B. pendula* | OP820513.1 | NC_072281.1 | 160535 | OP820513.1 | NC_072281.1 | 160535 | OP820513.1 | NC_072281.1 | 160536 |
| | | NC_072281.1 | OP820513.1 | 160535 | NC_072281.1 | OP820513.1 | 160535 | NC_072281.1 | OP820513.1 | 160536 |
| | | NC_072281.1 | no reference | 160535 | NC_072281.1 | no reference | 160535 | NC_072281.1 | no reference | 160536 |
| *03* | *B. klokovii* | MG386370.1 | NC_039996.1 | 160602 | MG386370.1 | NC_039996.1 | 160605 | MG386370.1 | NC_039996.1 | 160603 |
| | | NC_039996.1 | MG386370.1 | 160602 | NC_039996.1 | MG386370.1 | 160605 | NC_039996.1 | MG386370.1 | 160603 |
| | | NC_039996.1 | no reference | 160602 | NC_039996.1 | no reference | 160605 | NC_039996.1 | no reference | 160603 |
| *04* | *B. klokovii* | MG386370.1 | NC_039996.1 | 160602 | MG386370.1 | NC_039996.1 | 160602 | MG386370.1 | NC_039996.1 | 160602 |
| | | NC_039996.1 | MG386370.1 | 160602 | NC_039996.1 | MG386370.1 | 160602 | NC_039996.1 | MG386370.1 | 160602 |
| | | NC_039996.1 | no reference | 160602 | NC_039996.1 | no reference | 160602 | NC_039996.1 | no reference | 160602 |
| 05 | *B. klokovii* × *B. pendula* | OP820513.1 | NC_072281.1 | - | OP820513.1 | NC_072281.1 | 160553 | OP820513.1 | NC_072281.1 | 160553 |
| | | NC_072281.1 | OP820513.1 | - | NC_072281.1 | OP820513.1 | 160553 | NC_072281.1 | OP820513.1 | 160553 |
| | | NC_072281.1 | no reference | - | NC_072281.1 | no reference | - | NC_072281.1 | no reference | - |
| 06 | *B. klokovii* × *B. pendula* | OP820513.1 | NC_072281.1 | 160555 | OP820513.1 | NC_072281.1 | 160555 | OP820513.1 | NC_072281.1 | 160555 |
| | | NC_072281.1 | OP820513.1 | 160555 | NC_072281.1 | OP820513.1 | 160555 | NC_072281.1 | OP820513.1 | 160555 |
| | | NC_072281.1 | no reference | - | NC_072281.1 | no reference | - | NC_072281.1 | no reference | - |
| 07 | *B. pubescens* | MG386370.1 | NC_039996.1 | - | MG386370.1 | NC_039996.1 | - | MG386370.1 | NC_039996.1 | - |
| | | NC_039996.1 | MG386370.1 | - | NC_039996.1 | MG386370.1 | - | NC_039996.1 | MG386370.1 | - |
| | | NC_039996.1 | no reference | - | NC_039996.1 | no reference | - | NC_039996.1 | no reference | - |
| 08 | *B. pubescens* | MG386370.1 | NC_039996.1 | 160621 | MG386370.1 | NC_039996.1 | 160621 | MG386370.1 | NC_039996.1 | 160621 |
| | | NC_039996.1 | MG386370.1 | 160621 | NC_039996.1 | MG386370.1 | 160621 | NC_039996.1 | MG386370.1 | 160621 |
| | | NC_039996.1 | no reference | 160621 | NC_039996.1 | no reference | 160621 | NC_039996.1 | no reference | 160621 |
